# Uncovering the mechanistic basis of intracellular Raf inhibitor sensitivity reveals synergistic cotreatment strategies

**DOI:** 10.1101/2024.09.18.613772

**Authors:** Ethan G. Stoddard, Linglan Fang, Yuhao Zhong, Zachary E. Potter, Daniel S. Brush, Jessica J. Simon, Martin Golkowski, Dustin J. Maly

## Abstract

Raf kinases are crucial effectors in the Ras-Raf-Mek-Erk signaling pathway, making them important targets for the development of cancer therapeutics. This study investigates the variable potency of DFG-out-stabilizing Raf inhibitors in mutant KRas-expressing cell lines. We demonstrate that inhibitor potency correlates with basal Raf activity, with more active Raf being more susceptible to inhibition. We further show that DFG-out-stabilizing inhibitors disrupt high-affinity Raf-Mek interactions, promoting the formation of inhibited Raf dimers. Furthermore, we identify cobimetinib as a Mek inhibitor that uniquely sensitizes Raf to DFG-out inhibitors by disrupting autoinhibited Raf-Mek complexes. Building on this insight, we developed cobimetinib analogs with enhanced sensitization properties. Our findings provide a mechanistic framework for understanding the cellular determinants of DFG-out-stabilizing inhibitor sensitivity and offer strategies for optimizing synergistic Raf-Mek inhibitor combinations.

## Introduction

The Raf family protein kinases, consisting of ARaf, BRaf, and CRaf (Raf1) in humans, are major downstream effectors of active Ras (Ras-GTP)^1^. In the absence of active Ras, wild-type (WT) Raf kinases are primarily maintained in an autoinhibited monomeric state as part of a complex with inactive Mek and two 14-3-3 proteins^2–4^. Activation of Ras recruits Raf homo- and heterodimers to the plasma membrane, which phosphorylate the activation loop of Mek (Mek1/2)^5^. Activation loop-phosphorylated Mek1/2 then activates its sole substrate, Erk1/2, through phosphorylation of its activation loop. Erk1/2, in turn, phosphorylates numerous substrates, resulting in the modulation of signaling and transcriptional events that influence diverse cellular processes such as growth, differentiation, proliferation, and survival^6, 7^.

The prevalence of somatic mutations in BRaf and the upstream Ras GTPases that promote oncogenic signaling has provided motivation for the development of inhibitors that block Raf kinase activity^8, 9^. To date, three first-generation, ATP-competitive Raf inhibitors–dabrafenib, encorafenib, and vemurafenib–have been approved as single agents or as combination therapies with Mek inhibitors by the FDA^10^. These first-generation inhibitors stabilize a conformation of Raf’s ATP-binding site where a helix in the N-terminal lobe of the kinase domain–called the αC helix–is outwardly displaced from an active conformation. While effective in blocking the intracellular activity of V600 mutants of BRaf, these αC helix-out-stabilizing inhibitors have been found to promote “paradoxical activation” of downstream Mek and Erk across a wide concentration range in cells expressing non-V600 BRaf mutants or mutant Ras^11–13^. The phenomenon of paradoxical activation is a consequence of the fact that αC helix-out-stabilizing inhibitors promote Raf dimers that demonstrate negative cooperativity between the ATP-binding sites of inhibitor-bound protomers, resulting in the formation of catalytically active, single inhibitor-occupied dimers (**Fig. 1a**)^14^.

**Figure 1.**
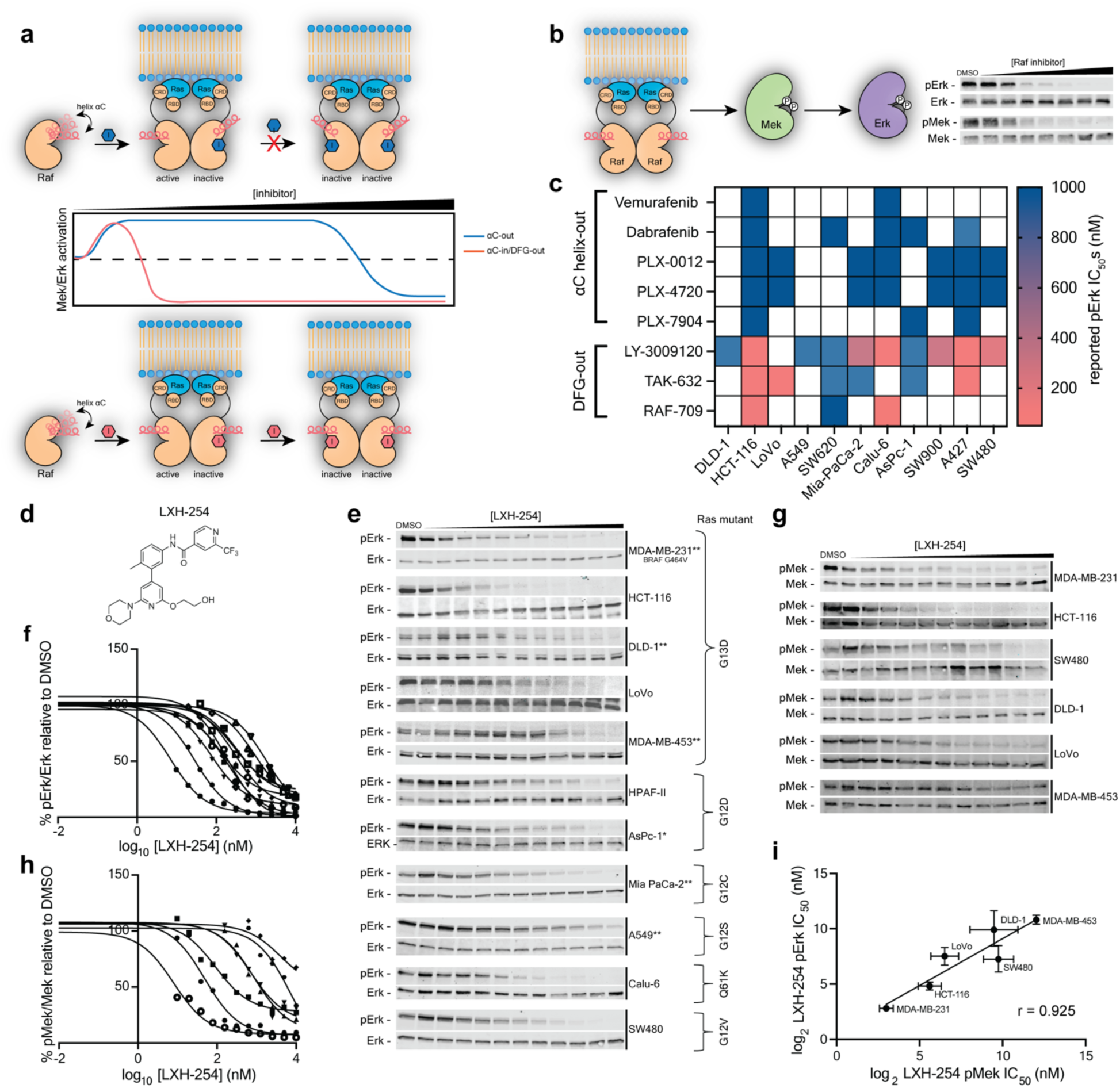
Raf inhibitor potency in various mutant KRas-expressing cell lines. **a,** Schematic of how Raf inhibitors that stabilize the αC helix-out and DFG-out conformations affect downstream Mek and Erk activation at different cellular concentrations. αC helix-out-stabilizing inhibitors exhibit “paradoxical activation” across a broad concentration range, while DFG-out-stabilizing inhibitors do not. **b,** Downstream Mek and Erk activation (pMek and pErk, respectively) as measured by western blot analysis. **c,** Heatmap representing previously reported pErk IC_50_ values for various αC helix-out- and DFG-out-stabilizing inhibitors in cells lines that express WT Raf and mutant KRas. Boxes in white indicate that no pErk IC_50_ values were reported. Darkest blue boxes denote a pErk IC_50_ value > 1 μM. **d,** Structure of the DFG-out-stabilizing inhibitor LXH-254. **e,** Representative western blots showing pErk levels in mutant KRas-expressing cells treated with a range of LXH-254 doses for 4 h (n=3, *n=1, **n=2). **f,** Quantification of normalized pErk percentages and determination of pErk IC_50_ values from the western blots obtained in (e). The percent pErk value shown at each LXH-254 concentration relative to DMSO is the mean of all replicates performed. IC_50_ curves were fit to percent pErk values obtained from all replicates. pErk IC_50_ values and 95% confidence intervals (CIs) for each cell line are listed in **Table S1**. **g,** Representative western blots showing pMek levels in mutant KRas-expressing cells treated with a range of LXH-254 doses for 4 h (n=2). **h,** Quantification of normalized pMek percentages and determination of pMek IC_50_ values from the western blots obtained in (g). The percent pMek value shown at each LXH-254 concentration relative to DMSO is the mean of both replicates. IC_50_ curves were fit to percent pMek values obtained from both replicates and pMek IC_50_ values are listed in Table S1. **i,** Correlation plot of LXH-254 pErk and pMek IC_50_ values for a comparative panel of six mutant KRas-expressing cell lines. Pearson’s *r* value is shown. Error bars represent 95% CIs for IC_50_ value curve fits.

Raf inhibitors that target alternative ATP-binding site conformations have also been developed. Inhibitors that stabilize the DFG-out conformation–where the Asp-Phe-Gly (DFG) motif in Raf’s activation loop is flipped–have been shown to be efficacious in blunting downstream signaling in multiple cellular contexts^15–19^. While DFG-out-stabilizing inhibitors also promote dimer formation, they potently block Raf’s kinase activity because they do not exhibit negative cooperativity between inhibitor-bound protomers^20^. Consequently, this class of inhibitors show promise for treating a broad spectrum of cancers with dysregulated Ras and Raf activity. Despite their broader inhibitory profile of DFG-out-stabilizing inhibitors, a number of basic mechanistic questions remain unanswered. The impact of cellular signaling state on these inhibitors’ potency remains incompletely defined, as does the mechanistic basis for observed differences in their effectiveness. Further, whether the same Mek inhibitors that have proven to be effective as combination therapies with αC helix-out-stabilizing inhibitors will show similar effectiveness with DFG-out stabilizing inhibitors is also unclear.

Here, we demonstrate that the potency of inhibitors stabilizing the DFG-out conformation of Raf’s ATP-binding is highly signaling state dependent in cells expressing mutant KRas and WT Raf kinases. We find that, counterintuitively, the more active that Raf is within a cell, the easier it is to inhibit, with differences in basal Raf activity accounting for observed variations in potency. By quantifying intracellular levels of target engagement of Raf protomers, we show that the potency of a DFG-out-stabilizing inhibitor in blocking downstream signaling is determined by the ease with which it recruits autoinhibited, monomeric Raf into dimers. A screen for synergistic Mek inhibitor cotreatment strategies revealed that cobimetinib uniquely sensitizes Raf kinases to inhibition by DFG-out-stabilizing inhibitors. This sensitization was found to result from cobimetinib’s ability to disrupt autoinhibited Raf kinase complexes and promote active dimers. Leveraging this mechanistic insight, we show that analogs of cobimetinib can be generated that provide enhanced sensitization to DFG-out-stabilizing inhibitors. Together, our findings provide a mechanistic framework for understanding the intracellular determinants of DFG-out-stabilizing inhibitor sensitivity and offer insight into the development of synergistic cotreatment strategies.

## Results

### LXH-254 demonstrates variable potency in mutant KRas-expressing cell lines

To understand how cellular context influences Raf’s sensitivity to inhibition, we compiled a list of reported Raf inhibitor potencies in blocking downstream Erk activation (hereafter referred to as pErk half-maximal inhibitory concentration (IC_50_) values) in several cells lines that express WT Raf and mutant KRas (**Fig. 1b**)^11, 21–27^. We found that αC helix-out-stabilizing inhibitors exhibited limited potency in cells expressing WT Raf, which is consistent with their propensity to promote single inhibitor-occupied Raf dimers that paradoxically activate downstream signaling (**Fig. 1c**). In contrast, inhibitors that stabilize the DFG-out conformation were able to potently block Erk activation. However, reported pErk IC_50_ values for the same DFG-out-stabilizing inhibitor ranged from double-digit nanomolar to micromolar, demonstrating cell context-dependent potency.

Reported pErk IC_50_ values of DFG-out-stabilizing inhibitors were obtained under varied experimental conditions, and systematic comparisons in different mutant KRas-expressing cell lines are lacking. Therefore, we conducted our own analysis of Raf-Mek-Erk pathway inhibition with LXH-254, a highly selective DFG-out-stabilizing clinical candidate (**Fig. 1d**)^8, 28^. Before performing full dose-responses with LXH-254 across multiple mutant KRas-expressing cell lines, we conducted a time course inhibition study in HCT-116 (G13D KRas/WT Raf) cells to determine the shortest treatment time that provided maximal decreases in pErk levels, thereby minimizing confounding secondary effects such as pathway rewiring and transcriptional reprogramming. We observed that, following a period of transient paradoxical activation, maximal and stable inhibition of downstream signaling was achieved after 4 h of LXH-254 treatment (**Extended Data Fig. 1a**). We next measured pErk IC_50_ values after a 4 h LXH-254 treatment across a diverse panel of eleven mutant KRas-expressing cancer cell lines (**Fig. 1e, f**). We observed a range of pErk IC_50_ values, from double-digit nanomolar (MDA-MB-231 and HCT-116 cells) to micromolar (MDA-MB-453) (**Table S1**). A comparative panel of six mutant KRas-expressing lines that represents a full range of measured pErk IC_50_ values was selected and LXH-254’s potency in blocking the phosphorylation of Raf’s direct substrate–Mek’s activation loop (pMek)–was determined (**Fig. 1g, h**). Consistent with pErk levels accurately reflecting Raf activity, intracellular pMek and pErk IC_50_ values were well correlated (**Fig. 1i**). Finally, we explored whether the relative potencies of LXH-254 determined at 4 h were representative of pathway inhibition at later time points by measuring pErk IC_50_ values in MDA-MB-453s (insensitive), Mia Paca-2s (moderate sensitivity), and HCT-116s (sensitive) after 24 h of LXH-254 treatment (**Extended Data Fig. 1b**). While we observed increased pErk IC_50_ values for all three lines, the relative differences in potencies were consistent with 4 h LXH-254 treatment, indicating that cell context-dependent sensitivity to DFG-out-stabilizing inhibitors is maintained after pathway rewiring and other secondary effects have taken hold.

### Variable potency in mutant KRas-expressing cell lines is a common characteristic of DFG-out-stabilizing inhibitors

We were curious whether the variable potency that LXH-254 demonstrated in mutant KRas-expressing cells is a common characteristic shared by inhibitors that stabilize the DFG-out conformation (**Fig. 1c**). To facilitate a comparative analysis, we developed LF-268, a pan-Raf inhibitor that is based on an entirely different scaffold than LXH-254 and features a trifluoromethylphenyl-benzamide moiety that stabilizes the DFG-out conformation (**Fig. 2a**). We determined LF-268’s IC_50_ values for purified ARaf, BRaf, and CRaf and found that it potently inhibited all three isoforms (**Fig. 2a** and **Extended Data Fig. 2a**). We next generated a click handle-derivatized version of LF-268, 268-TCO (**Fig. 2a** and **Extended Data Fig. 2a**), and used it for intracellular target profiling^29^. HCT-116 cells were first treated with DMSO or LF-268 as a competitor, followed by 268-TCO, and label free quantification of captured protein targets was performed after cell lysis and click handle enrichment (**Fig. 2b**). We observed that CRaf (Raf1) was the most LF-268-competed target, followed by a 14-3-3 isoform (YWHAB) and the Eph receptor tyrosine kinases EPHB4 and EPHA1/2. Competitive kinobead-based inhibitor profiling^30–32^ of LF-268 in HCT-116 lysates (**Fig. 2c**) confirmed that the 268-TCO competition experiment accurately reflected the kinase selectivity of LF-268 (**Table S2**). We next conducted global phosphoproteomic^33^ analysis of HCT-116 cells treated with LF-268, and discovered that only 54 of the >1500 phospho-peptides quantified decreased in abundance relative to DMSO. These 54 phospho-peptides correspond to 47 proteins, 35 of which constitute a STRING protein:protein interaction network^34^ centered around Erk1 and Erk2. (**Extended Data Fig. 2b** and **Table S3**). Phosphoproteomic analysis of kinobead-enriched lysate from LF-268-treated HCT-116 cells showed only two kinase phosphosites–the activation loop of Erk1 and the Y897 phospho-site of EPHA2–decreased by at least four-fold in response to LF-268 treatment (**Extended Data Fig. 2c** and **Table S4**) Together, our data show that LF-268 is a potent pan Raf inhibitor with a limited kinase target profile that mainly perturbs Erk-mediated signaling in mutant KRas-expressing cells.

**Figure 2.**
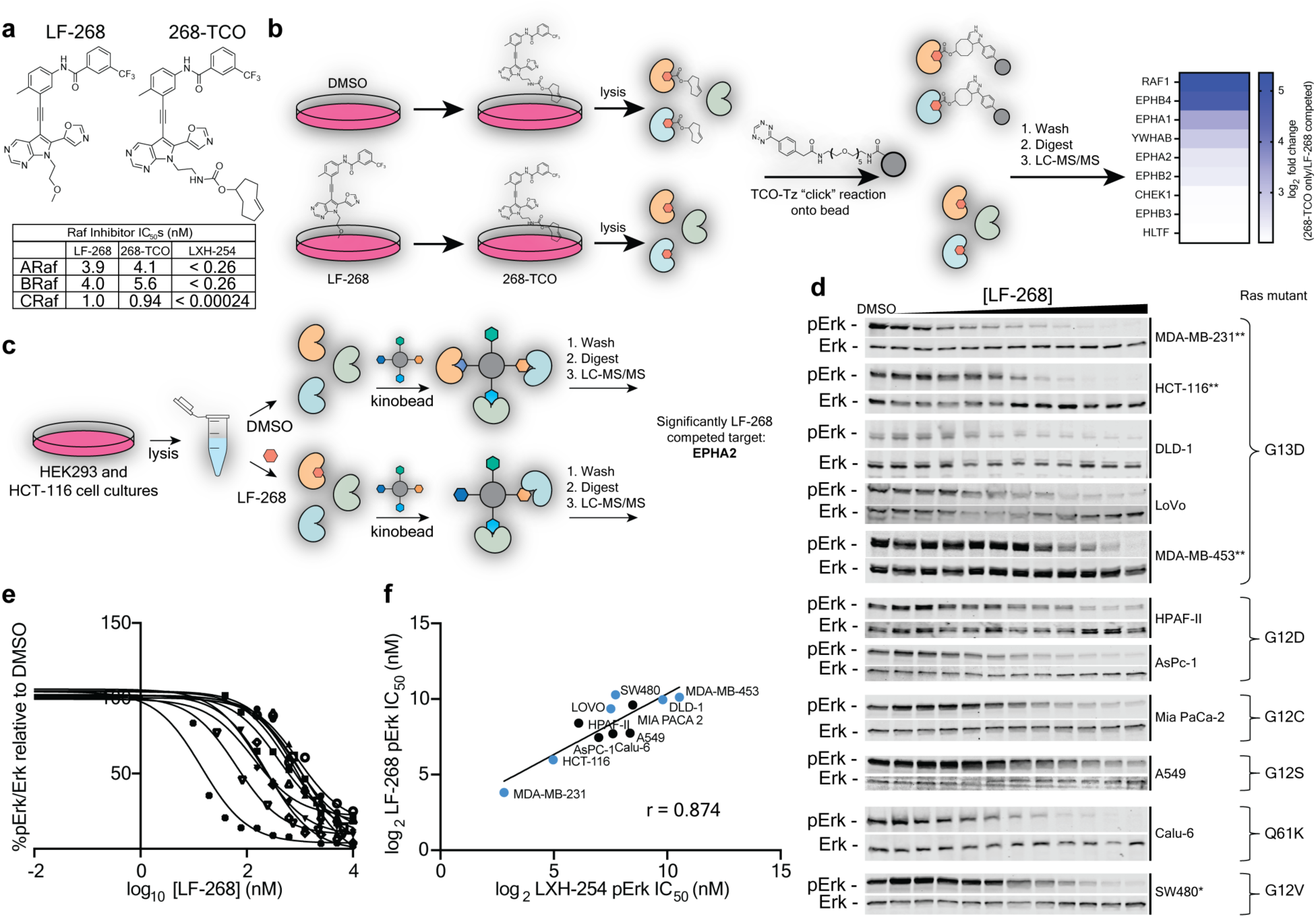
Characterization of an alternative DFG-out-stabilizing inhibitor of Raf. **a,** Structures of LF-268 and 268-TCO (*top*) and biochemical IC_50_ values (bottom) of LF-268, 268-TCO, and LXH-254 for purified Y301D/Y302D ARaf, WT BRaf, and Y340D/Y341D CRaf (LF-268: n=3, 268-TCO and LXH-254: n=1). **b,** Schematic of the competitive 268-TCO pulldown experiment performed with HCT-116 cells using tetrazine-linked beads for enrichment (*left*) and a heatmap (*right*) showing the proteins that were significantly competed (log_2_ ratio > 2 and *p* < 0.05) by 5 μM LF-268 compared to DMSO-treated cells (n=3). **c,** Schematic depicting the competitive kinobead-based inhibitor profiling workflow. Lysates were either incubated with 10 μM LF-268 or DMSO and only EPHA2 was significantly competed with a log_2_ ratio > 2 for 10 μM LF-268 relative to DMSO (n=2). **d,** Representative western blots showing pErk levels in mutant KRas-expressing cells treated with a range of LF-268 doses for 4 h (n=2, **n=3). **e,** Quantification of normalized pErk percentages and determination of pErk IC_50_ values from the western blots obtained in (d). The percent pErk value shown at each LF-268 concentration relative to DMSO is the mean of all replicates performed. IC_50_ curves were fit to the percent pErk values obtained from all replicates. pErk IC_50_ values and 95% confidence intervals (Cis) for each cell line are listed in **Table S1**. **f,** Correlation plot of LF-268 and LXH-254 pErk IC_50_ values for eleven mutant KRas-expressing cell lines. Pearson’s *r* value is shown. Values shown in blue indicate the six cell lines that comprise the comparative panel of mutant KRas-expressing cell lines.

Having confirmed that LF-268 possesses sufficient selectivity to provide confidence that observed decreases in pErk levels are mainly due to direct Raf inhibition, we quantified its potency in inhibiting downstream signaling in our panel of eleven mutant KRas-expressing cell lines. We observed a range of pErk IC_50_ values for LF-268 (**Fig. 2d, e**) that showed good correlation with LXH-254’s pErk IC_50_ values (**Fig. 2f**). Thus, the variable sensitivity of mutant KRas-expressing cell lines to LXH-254 appears to be a common characteristic of inhibitors that stabilize the DFG-out conformation.

### Elevated Raf activity correlates to increased sensitivity to DFG-out-stabilizing inhibitors

We next sought to identify the mechanistic basis behind Raf’s variable sensitivity to inhibition by LXH-254 and LF-268 in different mutant KRas-expressing lines. We found no apparent correlation between DFG-out inhibitor sensitivity and the identity or zygosity of the KRas mutant expressed in each line. For example, the most (HCT-116 and MDA-MB-231) and least (MDA-MB-453) sensitive cell lines to DFG-out-stabilizing inhibitors are both heterozygous for G13D KRas (**Fig. 1e** and **Fig 2d**). To determine if the basal signaling state of cells influence Raf’s sensitivity to inhibition, we measured the expression and phosphorylation levels of Raf-Mek-Erk pathway components across our comparative panel of six mutant KRas-expressing lines and looked for any potential correlations with pErk IC_50_ values. We found limited correlation between relative expression and phosphorylation levels of ARaf, BRaf, and CRaf and pErk IC_50_ values (**Extended Data Fig. 3a, b**). There was, however, a general trend towards elevated basal pErk levels and DFG-out inhibitor sensitivity (**Fig. 3a**). Strikingly, a strong inverse correlation was observed between basal pMek levels and pErk IC_50_ values in mutant KRas-expressing cells (**Fig. 3b, c**). Thus, it appears that the more active Raf is within a mutant KRas-expressing cell line, the more sensitive it is to inhibition by a DFG-out-stabilizing inhibitor.

**Figure 3.**
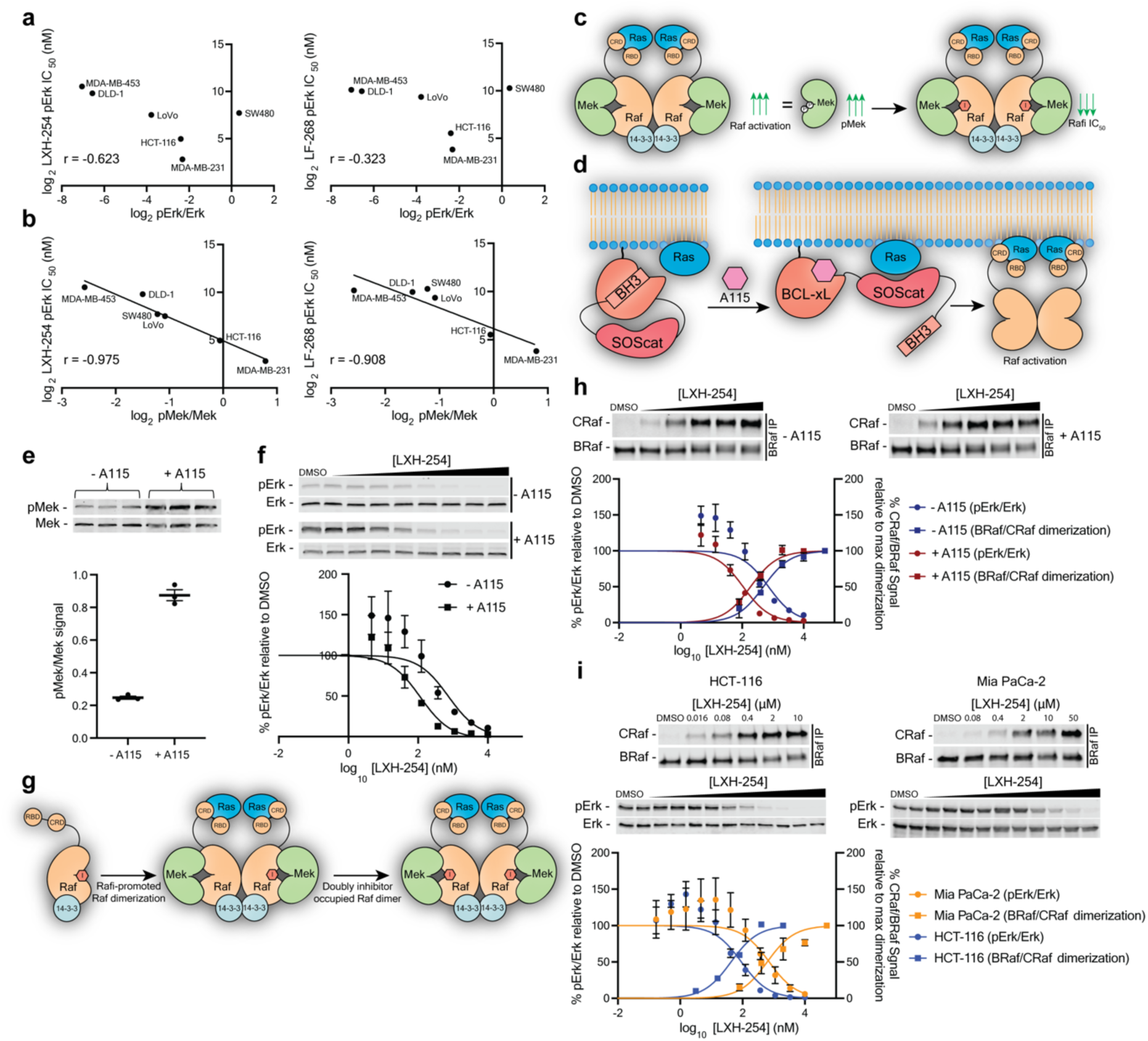
DFG-out-stabilizing inhibitors are more potent in cells lines with elevated Raf activity. **a,** Correlation plot of normalized pErk levels (n=1) determined by western blot versus LXH-254 (*left*) and LF-268 (*right*) pErk IC_50_ values across the comparative panel of mutant KRas-expressing cell lines. Pearson’s *r* values are shown. **b,** Correlation plot of normalized pMek levels (n=1) measured by western blot versus LXH-254 (*left*) and LF-268 (*right*) pErk IC_50_ values across the comparative panel of mutant KRas-expressing cell lines. Pearson’s *r* values are shown. **c,** Schematic depicting the relationship between increased Raf activity within cells, as measured by increased pMek, and enhanced sensitivity to inhibitors that stabilize the DFG-out conformation. **d,** Schematic depicting the activation of WT Ras with A115 in the CIAR-293 cell line. **e,** Western blot (*top*) and quantification (*bottom*) of pMek levels in CIAR-293 cells treated with 100 nM A115 or DMSO for 1 h. Bars equal the normalized pMek levels from n=3 replicates and error bars represent standard error of the mean (s.e.m.) **f,** Representative western blots (*top*) and quantification (*bottom*) of pErk levels in CIAR-293 cells pre-treated with A115 or DMSO for 1 h, followed by incubation with various doses of LXH-254 for 3 h. Normalized pErk percentages at each LXH-254 concentration (relative to DMSO-treated cells) equal the mean of n=3 replicates and error bars represent s.e.m. IC_50_ curves were fit to the percent pErk values obtained from all three replicates and pErk IC_50_ values and 95% confidence intervals (CIs) are listed in **Table S1**. **g,** Schematic depicting the conversion of monomeric Raf into a single inhibitor-occupied dimer and double inhibitor-occupied dimer. **h,** Representative western blots (*top*) and quantification (*bottom*) of CRaf levels co-immunoprecipitated with BRaf from CIAR-293 cells pre-treated with DMSO or A115 for 1 h, followed by incubating with a range of LXH-254 doses for 3 h. The percent CRaf levels (normalized to immunoprecipitated BRaf) shown at each LXH-254 concentration are the mean of n=3 replicates and error bars represent s.e.m. DC_50_ curves were fit to percent CRaf levels obtained from all three replicates and BRaf:CRaf dimerization DC_50_ values and 95% CIs are listed in **Table S1**. pErk IC_50_ curves from (f) are overlayed to demonstrate the concentration of LXH-254 where pErk IC_50_ and DC_50_ values intersect. **i,** Representative western blots (*top*) and quantification (*bottom*) of CRaf levels (co-immunoprecipitated with BRaf) and pErk levels in HCT-116 and Mia PaCa-2 cells treated with a range of LXH-254 doses for 3 h. The percent CRaf levels (normalized to immunoprecipitated BRaf) shown at each LXH-254 concentration are the mean of n=3 replicates and error bars represent s.e.m. DC_50_ curves were fit to percent CRaf levels obtained from all three replicates. The percent pErk value (normalized to Erk levels) shown at each LXH-254 concentration relative to DMSO is the mean of n=3 replicates performed. IC_50_ curves were fit to percent pErk values obtained from all replicates. BRaf:CRaf dimerization DC_50_ and pErk IC_50_ curves are overlayed. BRaf:CRaf dimerization DC_50_ values, pErk IC_50_ values, and 95% CIs are listed in **Table S1**.

To determine whether the correlation between elevated Raf activity and DFG-out inhibitor sensitivity holds within the same cellular context, we measured LXH-254’s potency in Chemically Inducible Activator of Ras (CIAR)-293 cells. CIAR-293 cells facilitate the rapid and direct activation of Ras, and subsequently Raf, in response to a small molecule, A115, that disrupts an autoinhibitory interaction within a membrane-localized CIAR protein switch (**Fig. 3d**)^35, 36^. We observed that the pErk IC_50_ value for LXH-254 was 4-fold lower in A115-treated CIAR-293 cells compared to DMSO-treated CIAR-293 cells (**Fig. 3f**). This reduction in LXH-254’s IC_50_ value directly corresponded to the four-fold higher pMek levels in A115-treated CIAR-293 cells (**Fig. 3e, f**). Our results suggest that the more active Raf is within mutant KRas-expressing cells, the more potently inhibitors that stabilize the DFG-out conformation are able to engage its ATP-binding site.

### Characterization of Inhibitor-Promoted Isoform-Specific Heterodimerization

Inhibitor-promoted Raf dimer formation is an important consideration in assessing the potency of intracellular pathway inhibition. Similar to αC helix-out-stabilizing inhibitors, inhibitors that stabilize the DFG-out conformation promote single inhibitor-occupied Raf dimers, and only after occupation of the second protomer is downstream pathway activation suppressed (**Fig. 1a**)^11^. Thus, any residual pMek and pErk observed at sub-saturating concentrations of DFG-out-stabilizing inhibitors could arise from either uninhibited Raf dimers, single inhibitor-occupied Raf dimers, or a combination of both. Therefore, we assessed direct engagement of Raf by measuring inhibitor-promoted half-maximal dimerization constant (DC_50_) values for endogenous BRaf and CRaf with co-immunoprecipitations (**Fig. 3g**). We first investigated whether LXH-254’s lower pErk IC_50_ value in CIAR-293 cells with elevated Raf activity, as measured by increased pMek levels (**Fig. 3e**), corresponded to a similar decrease in heterodimerization DC_50_ value. Consistent with the notion that the ATP-binding site of more active Raf is easier to engage with DFG-out-stabilizing inhibitors, we observed that lower concentrations of LXH-254 were required to promote 50% BRaf:CRaf heterodimerization in A115-treated CIAR-293 cells compared to untreated cells, although maximal heterodimerization levels were similar (**Extended Data Fig. 3c**). We also found that measured 50% pErk and 50% heterodimerization levels intersected at the same concentration of LXH-254 in both A115-treated and untreated CIAR-293 cells (**Fig. 3h**), suggesting that heterodimerization DC_50_ values represent inhibitor engagement of the second protomer of BRaf:CRaf heterodimers.

Next, we examined whether LXH-254’s variable potency in inhibiting the Raf-Mek-Erk pathway across different mutant KRas-expressing cell lines corresponded to its ability to promote BRaf:CRaf heterodimerization. We treated Mia PaCa-2 and HCT-116 cells with varying concentrations of LXH-254, and DC_50_ values were determined in each line by immunoprecipitating BRaf and quantifying the amount of CRaf co-enriched. We observed that LXH-254’s DC_50_ value in Mia PaCa-2 cells was ∼20-fold higher than in HCT-116 cells (**Fig. 3i**), corresponding to its ∼10-fold higher pErk IC_50_ value in Mia PaCa-2s relative to HCT-116 (**Fig. 1e-h**). As in CIAR-293 cells, the concentrations of LXH-254 that provided 50% co-immunoprecipitation of CRaf with BRaf intersected with a 50% reduction in pErk levels in each cell line. Size exclusion chromatography (SEC) analysis of Mia PaCa-2 cell lysates revealed that a saturating concentration of LXH-254 led to increased recruitment of BRaf, but not ARaf, into higher-size fractions with CRaf, likely representing dimeric signaling complexes (**Extended Data Fig. 4a**). We determined an CRaf:ARaf DC_50_ value in Mia PaCa-2 cells by immunoprecipitating CRaf and quantifying the amount of ARaf coenriched in the presence of varying concentrations of LXH-254. We found ARaf was only recruited into heterodimers with CRaf at LXH-254 concentrations (10-50 µM) where pErk levels decreased by >80% (**Extended Data Fig. 4b**). Thus, the variable ability of DFG-out-stabilizing inhibitors to promote doubly-occupied BRaf:CRaf heterodimers appears to be the major contributor to observed differences in Raf’s sensitivity to inhibition in mutant KRas-expressing cell lines.

### DFG-out-stabilizing inhibitors potently engage the second protomer of Raf dimers

When assessing the potency of inhibitors in suppressing signaling downstream of Raf homo- and heterodimers, two binding interactions–first and second protomer occupancy–must be considered. Inhibitors that stabilize the αC helix-out conformation, such as encorafenib and vemurafenib, exhibit a broad concentration range for paradoxical activation because, due to negative cooperativity, much higher drug concentrations are required to occupy the second protomer of active Raf dimers^11, 13^. Consistent with a lack of negative cooperativity, and the potential presence of positive cooperativity^20^, for DFG-out-stabilizing inhibitors in occupying the second protomer of Raf dimers, we observed that cells treated with LXH-254 for 4 h demonstrated limited paradoxical activation compared to αC helix-out-stabilizing encorafenib (**Extended Data Fig. 4c, d**). However, we found that robust but transient paradoxical activation occurred at shorter LXH-254 treatment times (**Extended Data Fig. 4e**), which is consistent with LXH-254 initially promoting single inhibitor-occupied, active Raf dimers from autoinhibited monomers, followed by more potent engagement and inhibition of the second protomer.

To determine whether the potency of DFG-out-stabilizing inhibitors for the second protomer of Raf dimers varies in different mutant KRas-expressing cell lines, we generated single inhibitor-occupied Raf dimers using encorafenib and measured LXH-254’s pErk IC_50_ value as a readout of second protomer engagement. Encorafenib exhibits strong negative cooperativity between Raf protomers and slow dissociation kinetics (t_½_ > 24 hours)^37–39^, which allows the quantitative formation of single inhibitor-occupied Raf dimers by performing inhibitor washout following cell treatment (**Fig. 4a**). To determine the optimal concentration of encorafenib for promoting maximal formation of single inhibitor-occupied Raf dimers, we treated HCT-116 and Mia Paca-2 cells with a dose range of encorafenib and measured pErk levels following washout (**Fig. 4b, c**). With these titrations, we calculated 50% maximal paradoxical activation (PA_50_) values and identified the concentration of encorafenib that provided maximal paradoxical activation for each cell line. Notably, measured PA_50_ values were seven-fold lower in HCT-116s than in Mia PaCa-2 cells (**Fig. 4b**), reflecting the relative sensitivities of these cell lines to inhibition by DFG-out-stabilizing inhibitors (**Fig. 2f**). We then quantified LXH-254’s pErk IC_50_ values for single encorafenib-occupied Raf dimers in HCT-116, Mia Paca-2, and MDA-MB-453 cells (**Fig. 4d, e and Extended Data Fig. 5a, b**). We observed that encorafenib pretreatment and washout resulted in a modest 2-3-fold decrease in LXH-254’s pErk IC_50_ value in HCT-116 cells (**Fig. 4d**) but more dramatic ∼30-fold and ∼20-fold decreases in Mia Paca-2 and MDA-MB-453 cells, respectively (**Fig. 4e** and **Extended Data Fig. 5a, b**). Despite the disparate sensitivity of these cell lines to DFG-out-stabilizing inhibitors alone, the potency of LXH-254 in all three encorafenib-pretreated cell lines converged within a 2-fold range (pErk IC_50_ values = 28-50 nM). Collectively, encorafenib’s ability to more potently promote single inhibitor-occupied Raf dimers in HCT-116s than Mia PaCa-2s, and the similar sensitivity of single encorafenib-occupied Raf dimers to inhibition by LXH-254 in different mutant KRas-expressing cell lines suggests that the binding event that recruits monomeric Raf into dimers–first protomer occupation–determines the sensitivity of a cell line to Raf inhibition by DFG-out-stabilizing inhibitors.

**Figure 4.**
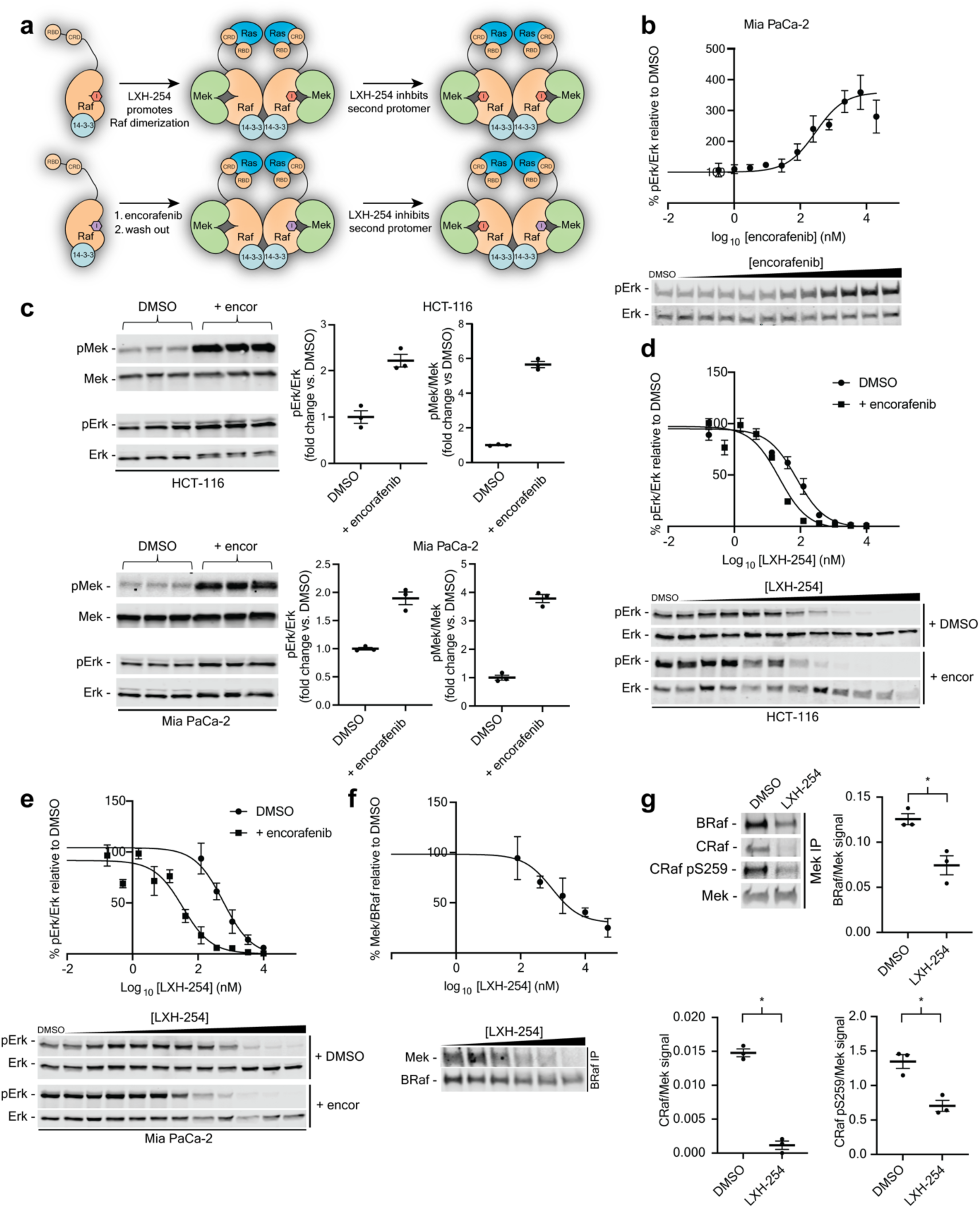
The potency of DFG-out-stabilizing inhibitors is determined by first protomer occupation of Raf dimers. **a,** Schematic showing how encorafenib, an αC helix-out-stabilizing inhibitor, can be used to quantitatively generate single inhibitor-occupied Raf dimers following wash out (*bottom*). Second protomer occupation by DFG-out-stabilizing inhibitors is quantified by measuring inhibition of Erk activation. **b,** Representative western blot (*bottom*) and quantification (*top*) of pErk levels in Mia PaCa-2 cells treated with a range of encorafenib doses for 1 h followed by a 1 h washout. The percent pErk values (relative to DMSO-treated cells) shown at each encorafenib concentration are the mean of n=3 replicates and error bars represent standard error of the mean (s.e.m.). The curve was fit to percent pErk values obtained from all three replicates and the pErk PA_50_ value and 95% CI are listed in **Table S1**. **c,** Western blots (*left*) and quantification (*right*) of pMek and pErk levels in HCT-116 (*top*) and Mia PaCa-2 (*bottom*) cells treated with 1 μM encorafenib or DMSO for 1 h followed by a 1 h washout. Bars equal the mean of n=3 replicates and error bars represent s.e.m. **d and e,** Representative western blots (*bottom*) and quantification (*top*) of pErk levels in HCT-116 (**d**) and Mia PaCa-2 (**e**) cells pre-treated with 1 μM encorafenib for 1 h, followed by washout and subsequent incubation with a range of LXH-254 doses for 3 h. The normalized percent pErk values (relative to DMSO-treated cells) shown at each LXH-254 concentration are the mean of n=3 replicates and error bars represent s.e.m. IC_50_ curves were fit to the percent pErk values obtained from all three replicates and pErk IC_50_ values and 95% confidence intervals (CIs) are listed in **Table S1**. pErk IC_50_ curves shown for DMSO-treated cells are from **Figure 3i**. **f,** Representative western blot (*bottom*) and quantification (*top*) of Mek co-immunoprecipitated with BRaf from Mia PaCa-2 cells treated with a range of LXH-254 doses for 3 h. The percent Mek (normalized to co-immunoprecipitated BRaf) values shown at each LXH-254 concentration are the mean of n=3 replicates and error bars represent s.e.m. A half-maximal competition (CC_50_) curve was fit to the percent Mek values obtained from all three replicates and the CC_50_ value and 95% CI is listed in **Table S1**. **g,** Representative western blots and quantification BRaf, CRaf, and CRaf pS259 co-immunoprecipitated with Mek from Mia PaCa-2 cells treated with 10 μM LXH-254 for 3 h. The bars shown are the Mek-normalized mean of n=3 replicates and the error bars represent s.e.m. *denotes significance as determined by two-sided Student’s t-test (*p* value < 0.05).

The low pErk IC_50_ values that LXH-254 and **LF-268** exhibited in MDA-MB-231 cells (**Fig. 1f** and **Fig. 2e**), which is the only mutant KRas-expressing cell line we profiled that expresses a Raf mutant (G464V), is consistent with the observed ability of DFG-out-stabilizing inhibitors to potently engage the second protomer of single encorafenib-occupied dimers. Previous studies have demonstrated that the G466V mutation in BRaf promotes basal Raf dimerization^40^, which most likely enables DFG-out-stabilizing inhibitors to engage both protomers of Raf dimers without the initial, potency-limiting step of recruiting monomeric Raf into dimers.

### DFG-out-stabilizing inhibitors disrupt high affinity Raf-Mek interactions

In addition to recruiting BRaf into high-size fractions with CRaf, our SEC analysis showed that LXH-254 treatment led to less Mek co-eluting with BRaf in high-size fractions (**Extended Data Fig. 6a**) relative to untreated Mia PaCa-2 cells. Consistent with this observation, we found that LXH-254 treatment dose-dependently decreased the amount of Mek that co-immunoprecipitated with BRaf (**Fig. 4f**) and that the 50% competition constant (CC_50_) for Mek co-enrichment was similar to the BRaf:CRaf heterodimerization DC_50_ and pErk IC_50_ values in the same cell line (**Fig. 3i**). Furthermore, LXH-254 treatment also significantly decreased the amount of BRaf and CRaf that co-enriched with immunoprecipitated Mek (**Fig. 4g**), suggesting a competitive relationship between DFG-out-stabilizing inhibitor binding and high affinity BRaf:Mek and CRaf:Mek interactions.

While we observed that LXH-254 treatment led to a dramatic decrease in the total amount of CRaf that co-immunoprecipitated with Mek, CRaf that is phosphorylated at the S259 autoinhibitory site^41^ was only minimally competed. LXH-254’s modest disruption of the pS259 CRaf:Mek complex is consistent with our observations that it did not potently engage the ATP-binding site of pS259 CRaf (**Extended Data Fig. 6b, c**) or recruit pS259 CRaf into heterodimers with BRaf (**Extended Data Fig. 6d**). LXH-254 treatment also did not modulate phosphorylation levels at the pS259 site (**Extended Data Fig. 6e**). Together, our data show that DFG-out inhibitor binding to BRaf and CRaf antagonizes their high affinity interactions with Mek but that Raf species phosphorylated at autoinhibitory 14-3-3 binding sites represent a population that is not recruited into DFG-out inhibitor-promoted Raf dimers.

### Pharmacologically modulating Raf:Mek complexes influences Raf’s sensitivity to DFG-out-stabilizing inhibitors

Diverse effects of Mek inhibitors on Raf:Mek complexes have been reported^42–44^, motivating us to investigate whether a synergistic co-treatment strategy could be devised. To this end, we identified doses of five diverse Mek inhibitors (**Fig. 5a**) that achieved partial (∼85%) Mek inhibition, measured as a decrease in pErk levels, in Mia Paca-2 cells (**Extended Data Fig. 7a and Fig. 5b**). As expected, these partial inhibitory doses yielded varying effects on pMek levels (**Extended Data Fig. 7b**). Next, we quantified LXH-254’s potency in Mia Paca-2 cells pretreated with partial (∼85%) inhibitory dose of each Mek inhibitor (**Fig. 5c, d**) and found that four of the five Mek inhibitors tested either increased or had little effect on pErk IC_50_ values. However, cobimetinib pretreatment led to a ∼4-fold increase in LXH-254’s potency. A similar trend was observed for the partial (∼85%) inhibitory doses of the five Mek inhibitors on LXH-254’s BRaf:CRaf DC_50_ values, with only cobimetinib pretreatment resulting in lower doses of LXH-254 promoting heterodimerization relative to DMSO (**Extended Data Fig. 8a-c**). Thus, cobimetinib appears to enhance LXH-254’s ability to block downstream signaling by promoting the formation of double inhibitor-occupied dimers, while other Mek inhibitors do not. To quantify the full extent of cobimetinib’s sensitization of cells to Raf inhibition, we measured LXH-254’s ability to block pErk formation in the least sensitive mutant KRas-expressing cell line (MDA-MB-453) pretreated with a range of cobimetinib concentrations (**Fig. 5e, f**). Bliss analysis of the titration data revealed a significant level of synergy between LXH-254 and cobimetinib, with an average Bliss score of 8.72 (*p* < 0.05, **Fig. 5g**).

**Figure 5.**
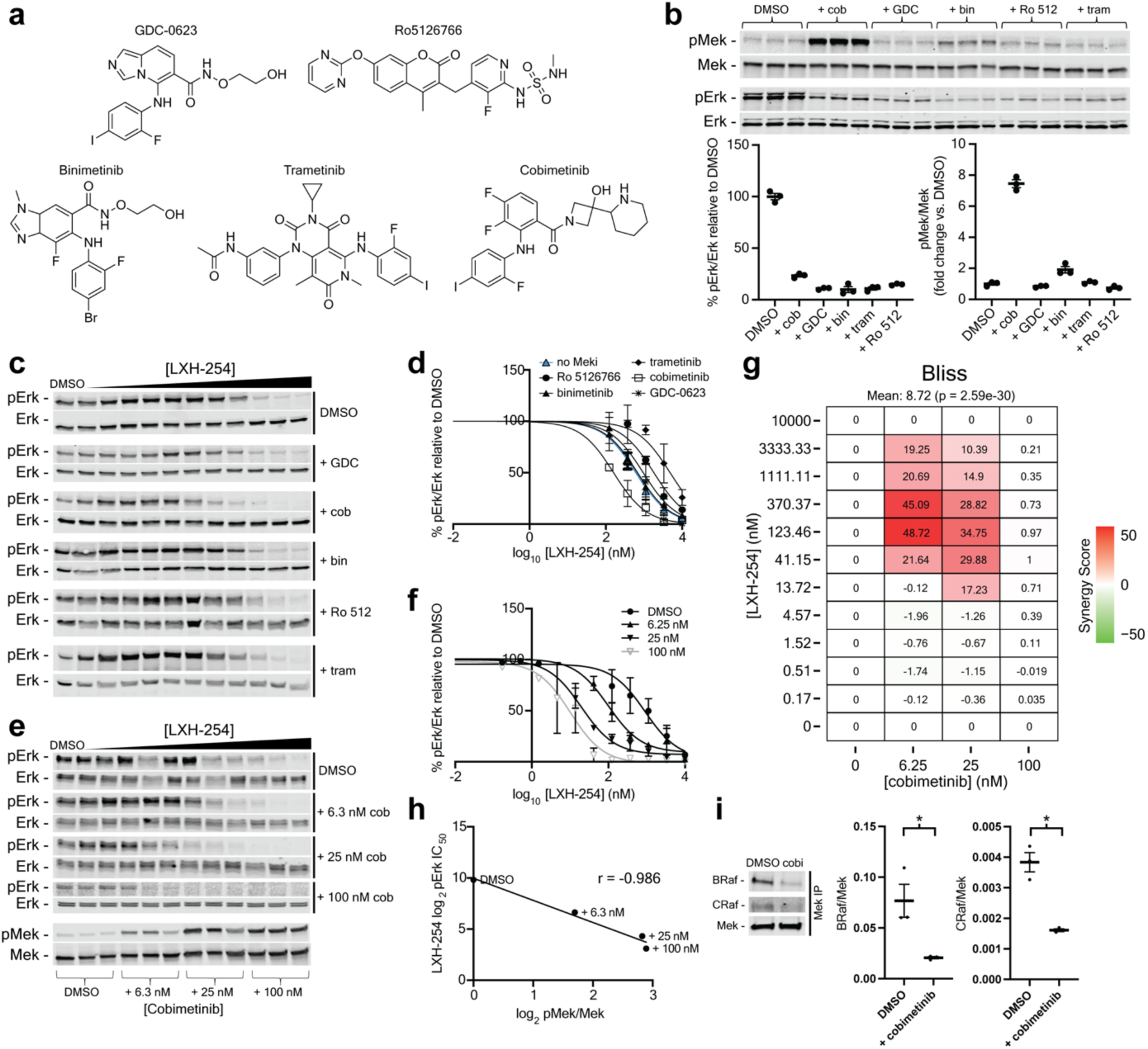
Cobimetinib sensitizes mutant KRas-expressing cells to LXH-254 by disrupting high affinity Mek/Raf complexes. **a,** Structures of clinical Mek inhibitors used in this study. **b,** Western blot (*top*) and quantification (bottom) of pMek and pErk levels in Mia PaCa-2 cells treated with partial (∼85%) inhibitory doses of the Mek inhibitors shown in (a) or DMSO for 4 h. Bars equal the mean of n=3 replicates and error bars represent standard error of the mean (s.e.m.). Mek inhibitor concentrations used: cobimetinib = 40 nM, Ro 5126766 = 6 nM, binimetinib = 16 nM, trametinib = 2 nM, GDC-0623 = 2 nM. **c and d,** Representative western blots (**c**) and quantification (**d**) of pErk levels in Mia PaCa-2 cells treated with DMSO or partial inhibitory doses of Mek inhibitors for 1 h, followed by incubation with a range of LXH-254 doses for 3 h. The normalized percent pErk values (relative to DMSO-treated cells) shown at each LXH-254 concentration are the mean of n=3 replicates and error bars represent s.e.m. IC_50_ curves were fit to the percent pErk values obtained from all three replicates and pErk IC_50_ values and 95% confidence intervals (CIs) are listed in **Table S1**. pErk IC_50_ curves shown for DMSO-treated cells are from **Figure 3i**. **e,** Representative western blots of pErk levels (*top*) in Mia PaCa-2 cells pre-treated with DMSO or a range (6.3, 25, or 100 nM) of cobimetinib doses for 1 h, followed by incubation with a range of LXH-254 doses for 3 h. The bottom western blot shows pMek levels for cells treated only with DMSO or a range (6.3, 25, or 100 nM) of cobimetinib doses. **f,** Quantification of percent pErk (relative to DMSO-treated cells) levels from (e). The percent pErk levels shown at each LXH-254 concentration are the mean of n=3 replicates and error bars represent s.e.m. IC_50_ curves were fit to the percent pErk values obtained from all three replicates. Calculated pErk IC_50_ values and 95% CIs are listed in **Table S1**. pErk IC_50_ curves shown for DMSO-treated cells are from **Figure 3i**. **g,** Bliss synergy analysis of cobimetinib and LXH-254 using the percent pErk values from the data shown in (f) using SynergyFinder. Results are depicted as a heatmap of average Bliss scores. **h,** Correlation plot of the log_2_pErk IC_50_ values and normalized pMek levels determined in (e). Pearson’s *r* value is shown. **i,** Representative western blots (*left*) and quantification (*right*) of BRaf and CRaf levels co-immunoprecipitated with Mek from Mia PaCa-2 cells treated with 40 nM cobimetinib for 4 h. The bars shown are the Mek-normalized mean of n=3 replicates and the error bars represent s.e.m. *indicates significance as determined by two-sided Student’s t-test (*p* value < 0.05).

### Cobimetinib sensitizes Raf to LXH-254 inhibition by promoting active Raf dimer formation

Previous studies have demonstrated that cobimetinib promotes increased pMek levels and BRaf:CRaf heterodimerization in DFG-out inhibitor-sensitive HCT-116 cells^44^. We found that cobimetinib also led to dose-dependent increases in pMek levels in MDA-MB-453 and Mia Paca-2 cells (**Extended Data Fig. 7b**), suggesting it leads to the promotion of active Raf dimers in mutant KRas-expressing cell lines that are less sensitive to DFG-out-stabilizing inhibitors. Further confirming that cobimetinib promotes active Raf dimers, we observed that provision of cobimetinib, like LXH-254, increased proximity labeling of all Raf isoforms by TurboID-KRas in A115-activated CIAR-293 cells compared to DMSO or the non-sensitizing Mek inhibitor GDC-0623 (**Extended Data Fig. 8d, e**). Notably, we found that there was a strong inverse correlation between cobimetinib’s promotion of increased pMek levels and its capacity to lower LXH-254 pErk IC_50_ values (**Fig. 5h**). Thus, cobimetinib’s ability to promote active Raf dimers appears to sensitize Raf to inhibition by DFG-out-stabilizing inhibitors. A partial inhibitory dose of cobimetinib also promoted paradoxical activation of Raf at lower concentrations of αC helix-out-stabilizing encorafenib (**Extended Data Fig. 8f**), which is consistent with the notion that cobimetinib’s interaction with Mek destabilizes autoinhibited complexes of monomeric Raf to allow more facile promotion of single inhibitor-occupied dimers.

We next explored whether cobimetinib’s sensitization of monomeric Raf to inhibitors is due to a similar disruption of high affinity Raf:Mek interactions as LXH-254. First, we subjected the lysates of Mia Paca-2 cells treated with cobimetinib or non-sensitizing GDC-0623 to SEC and quantified Raf and Mek elution by western blot analysis (**Extended Data Fig. 9a, b**). Consistent with its ability to promote active Raf dimers^44^, we found that cobimetinib promoted CRaf’s and, to a lesser extent, ARaf’s elution at a higher molecular weight relative to GDC-0623. We also observed that cobimetinib led to less co-elution of Mek within high-size fractions containing BRaf and CRaf compared to GDC-0623, demonstrating the differential effects of these Mek inhibitors on high affinity Raf:Mek interactions. Next, we directly probed the effect of cobimetinib on Mek:Raf complexes by immunoprecipitating Mek and quantifying co-enriched BRaf and CRaf (**Fig. 5i**). Like LXH-254, we found that cobimetinib significantly decreased BRaf and CRaf co-enrichment with immunoprecipitated Mek, confirming the direct disruption of high affinity Raf:Mek complexes. Together, our results support a model in which cobimetinib’s attenuation of a high affinity Raf:Mek interaction leads to more facile recruitment of Raf into inhibitor-promoted Raf dimers, thereby increasing the potency of inhibitors that stabilize the DFG-out conformation.

### Mek inhibitors can be tuned to increase Raf inhibitor sensitization

Based on our model, we predicted that Mek inhibitors could be optimized to enhance Raf’s sensitivity to DFG-out-stabilizing inhibitors by promoting increased disruption of high affinity Raf:Mek interactions. Comparison of the co-crystal structures of sensitizing cobimetinib and non-sensitizing GDC-0623 bound to the autoinhibited BRaf:Mek complex (**Fig. 6a**) showed that the biaryl anilines of both inhibitors form comparable binding contacts with a hydrophobic pocket in Mek, but their alcohol-containing moieties form dissimilar polar interactions^42^. Because these aforementioned interactions appear to stabilize different conformations of an α-helix in the N-terminal region of Mek’s activation loop that is in close proximity to autoinhibited BRaf (**Fig. 6b**), we hypothesized that Mek inhibitors that differentially modulate this region would elicit varying levels of DFG-out-stabilizing inhibitor sensitization. To test this hypothesis, we generated a small panel of Mek inhibitors by replacing the 4-azetidinol moiety of cobimetinib with different substituents (**Fig. 6c**). We determined partial (∼85%) inhibitory doses of each Mek inhibitor (**Extended Data Fig. 10a, b**) and tested their ability to sensitize Raf to inhibition by LXH-254 in Mia Paca-2 cells. We found that partial inhibitory doses of these Mek inhibitors yielded a range of LXH-254 pErk IC_50_ values, with **3** providing the highest level of sensitization and **5** the least (**Fig. 6d, e**). Thus, the identity of the substituent that extends towards an α-helical region of Mek’s activation loop can have a substantial impact on the level of DFG-out inhibitor sensitization provided.

**Figure 6.**
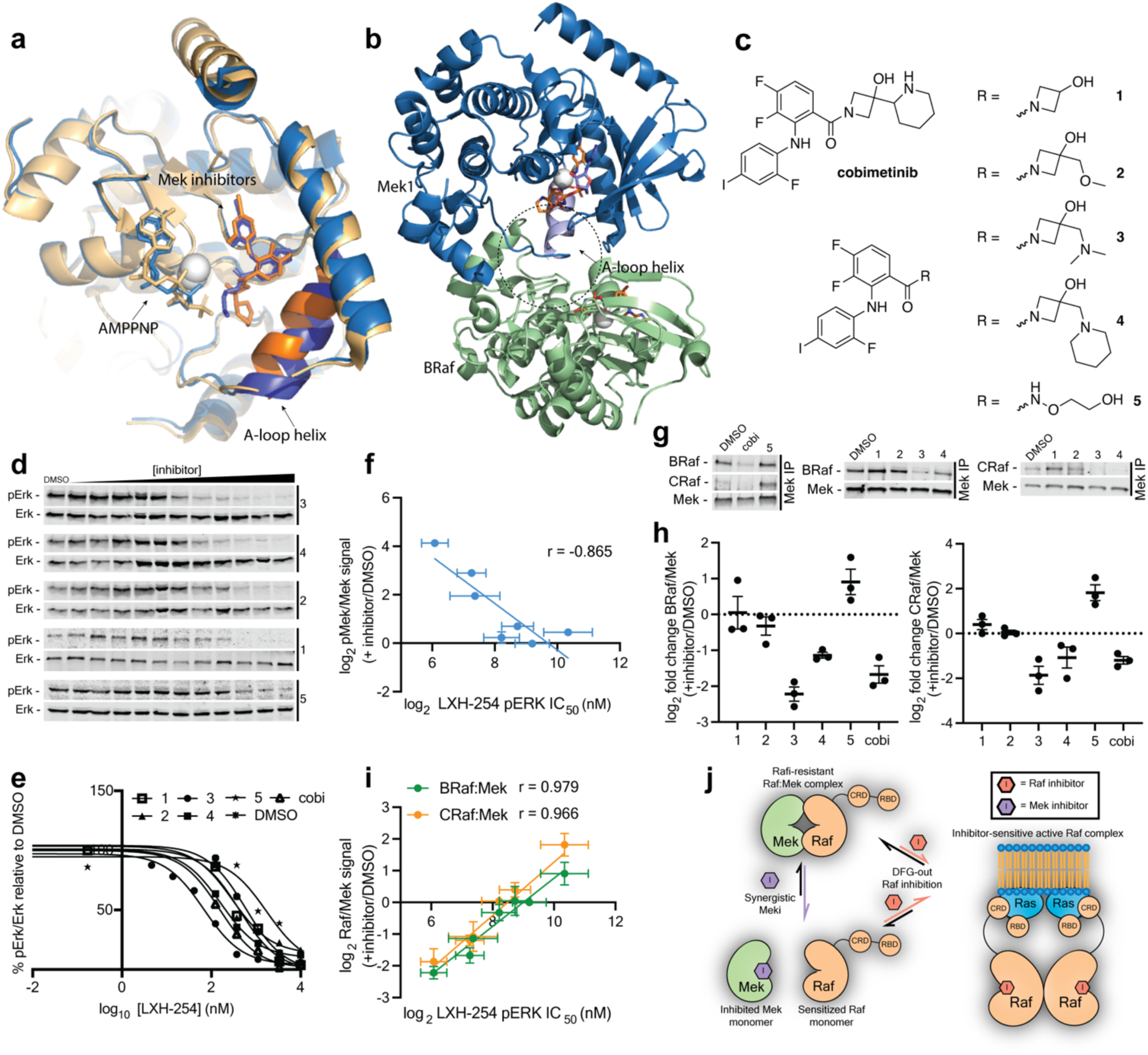
Mek inhibitors can be optimized to disrupt high affinity Raf:Mek interactions and provide sensitization to DFG-out-stabilizing Raf inhibitors. (a) Aligned co-crystal structures of cobimetinib (*orange*) and GDC-0623 (*blue*) bound to AMP-PNP:Raf:Mek complex (PDB IDs: 7M0V and 6NYB). (b) Co-crystal structure of the cobimetinib:AMP-PNP:Raf:Mek complex showing the α-helix in Mek’s activation loop (PDB ID: 7M0V). (c) Structures of Mek inhibitors **1-5** and cobimetinib. (d, e) Representative western blots (d) and quantification (e) of pErk levels in Mia PaCa-2 cells treated with DMSO or partial inhibitory doses of inhibitors **1-5** for 1 h, followed by incubation with a range of LXH-254 doses for 3 h. Cobimetinib cotreatment data are from the experiments in Fig. 5c, d. The normalized percent pErk values (relative to DMSO-treated cells) shown at each LXH-254 concentration are the mean of n=3 replicates. IC_50_ curves were fit to the percent pErk values obtained from all three replicates and pErk IC_50_ values and 95% confidence intervals (CIs) are listed in **Table S1**. Mek inhibitor concentrations: **1** = 80 nM, **2** = 360 nM, **3** = 120 nM, **4** = 600 nM, **5** = 10 nM. (f) Correlation plot of LXH-254 pErk IC_50_s from (e) and calculated pMek levels from **Extended Data Fig. 10c**. Pearson’s *r* value is shown. Error bars for pMek levels represent standard error of the mean (s.e.m.) and for pErk IC_50_ levels represent 95% CIs. (g, h) Representative western blots (g) and quantification (h) of BRaf and CRaf levels co-immunoprecipitated with Mek from Mia PaCa-2 cells treated with partial inhibitory doses of inhibitors **1-5** or cobimetinib for 4 h. The bars shown are the Mek-normalized mean of n=3 replicates and the error bars represent s.e.m. Mek inhibitor concentrations: **1** = 80 nM, **2** = 360 nM, **3** = 120 nM, **4** = 600 nM, **5** = 10 nM. (i) Correlation plot of the pErk IC_50_ values determined in (d, e) and levels of BRaf and CRaf co-immunoprecipitated as determined in (g, h). Pearson’s *r* values are shown. Error bars for BRaf and CRaf levels co-immunoprecipitated with MEK represent s.e.m. and for pErk IC_50_ levels represent 95% CIs. (j) Model showing how Mek inhibitors that destabilize high affinity Raf:Mek interactions sensitize Raf to DFG-out-stabilizing inhibitors.

We next determined whether the varying levels of sensitization provided by Mek inhibitors **1**-**5** is related to their ability to promote Raf dimerization through disruption of high affinity Raf:Mek interactions. We observed that partial inhibitory doses of **1**-**5** resulted in varying degrees of Raf dimerization, as indicated by measured pMek levels (**Extended Data Fig. 10c**), and that there was an inverse correlation between a Mek inhibitor’s ability to promote increased pMek levels at a partial inhibitory dose and capacity to lower LXH-254 pErk IC_50_ values (**Fig. 6f**). We next tested the ability of **1**-**5** to directly disrupt high affinity Raf:Mek interactions by immunoprecipitating Mek from Mia Paca-2 cells treated with partial inhibitory doses of each Mek inhibitor and probing for levels of co-enriched BRaf and CRaf. We observed that inhibitors **1**-**5** varied in their capacity to disrupt Mek’s interaction with BRaf and CRaf (**Fig. 6g, h**). Consistent with the disruption of high affinity Raf:Mek interactions serving as a mechanism for providing sensitization of Raf to DFG-out-stabilizing inhibitors, we observed that there was a strong correlation between Mek:Raf co-immunoprecipitation levels and LXH-254 pErk IC_50_ values in cells co-treated with inhibitors **1**-**5** (**Fig. 6i**). Thus, Mek inhibitors tuned to disrupt autoinhibitory Raf:Mek complexes both increase the activity of Raf and provide sensitization to inhibitors that stabilize the DFG-out conformation. These data point to a model in which the occupancy of Raf by an inhibitor that stabilizes the DFG-out conformation is antagonistic with a high affinity interaction between Raf and Mek (**Fig. 6j**), which Mek inhibitors can be tuned to disrupt.

## Discussion

Several Raf inhibitors that stabilize the αC helix-out conformation have been approved for treating Erdheim-Chester disease and melanomas driven by V600 mutants of BRaf^45, 46^.

However, these inhibitors have proven much less effective in blunting elevated Mek and Erk activation levels in cancer cell lines that express mutant Ras or other non-V600 BRaf mutants due to their promotion of Raf dimers with only one protomer potently engaged by an inhibitor, a phenomenon referred to as paradoxical activation^11–13^. Inhibitors that stabilize the DFG-out conformation have been demonstrated to suppress Mek and Erk activation in broader cellular contexts. While DFG-out-stabilizing inhibitors also promote Raf dimer formation, they are capable of potently engaging both protomers^47^. Here, we systematically explored the potency of DFG-out stabilizing inhibitors in mutant KRas-expressing cell lines (**Fig. 1**, **Fig. 2**) and determined how inhibitor engagement affected Raf’s formation of intermolecular complexes (**Fig. 3**, **Fig.4**). We also investigated whether specific Mek inhibitors could be combined with DFG-out stabilizing inhibitors to synergistically suppress downstream signaling (**Fig. 5**). By directly probing observed effects on pathway inhibition under short cellular treatment regimes, which minimized secondary effects, we gained mechanistic insight into why Raf is more easily inhibited under certain cellular signaling conditions. This insight was used to design Mek inhibitors that are more effective as part of co-treatment regimes with DFG-out-stabilizing inhibitors (**Fig. 6**).

The previous observation that LXH-254 and another inhibitor that stabilizes the DFG-out conformation (tovorafenib) demonstrate positive cooperativity between protomers within purified Raf dimers^20^ is consistent with our finding that DFG-out-stabilizing inhibitors more potently block Erk activation in cell lines with elevated Raf activity, and presumably higher levels of dimeric Raf (**Fig. 3b, e, f**). Our studies probing target engagement levels for endogenous Raf kinases are also concordant with nanoBRET assays performed with exogenously-expressed constructs demonstrating that DFG-out-stabilizing inhibitors more potently engage Raf kinases in cells with elevated RAS-GTP levels^48^. The aforementioned results, along with the demonstration that DFG-out-stabilizing inhibitors potently block the activity of the second protomer of single inhibitor-occupied Raf dimers (**Fig. 4d, e and Extended Data Fig. 5b**), highlight that DFG-out-stabilizing inhibitors are selective for dimeric Raf over other Raf species.

The reason for higher Raf activity in some mutant KRas-expressing cell lines compared to others is unclear. Neither the identity of the KRas mutant expressed in a cell line nor whether it is homo- or heterozygous correlated with basal pMek levels (**Fig. 1e**, **Fig. 2d and Fig. 3b**).

Because previous studies have shown that ARaf is less active and more resistant to inhibition by DFG-out-stabilizing inhibitors than BRaf or CRaf,^20, 28, 48, 49^ a model where cells are more resistant to Raf inhibition if ARaf makes a larger contribution to signaling downstream of activated Ras would be consistent with our results. However, we found that ARaf expression levels did not correlate with measured pErk IC_50_ values (**Extended Data Fig. 3a**). Moreover, ARaf was only recruited into doubly-occupied dimers with CRaf at inhibitor concentrations where most Erk activation was already suppressed (**Extended Data Fig. 4a,b**). In multiple cell lines, the same concentration of inhibitor that led to a 50% reduction in pErk levels also led to 50% maximal BRaf:CRaf heterodimerization, suggesting that the degree of basal dimerization of these Raf isoforms is a major determinant of pErk IC_50_ values (**Fig. 3h,i**). Given the demonstration that ARaf expression levels can affect cellular sensitivity to LXH-254^28^ and ARaf mutations can confer resistance in a clinically relevant setting^50^, this isoform likely plays a dominant role in some contexts.

We found that occupation of BRaf’s and CRaf’s ATP-binding sites with DFG-out-stabilizing inhibitors led to their recruitment into BRaf:CRaf heterodimers from autoinhibited monomeric complexes (**Fig. 3h,i**). Our results suggest that Mek forms higher affinity interactions with inactive, monomeric BRaf and CRaf than within dimeric Raf complexes.

Moreover, the overall sensitivity of a cell to pathway inhibition appears to be correlated to the ease with which these autoinhibited BRaf:Mek and CRaf:Mek complexes can be disrupted. An autoinhibited complex consisting of BRaf that is phosphorylated at S365 (which creates a high affinity 14-3-3-binding site)^41^, two 14-3-3 proteins, and Mek has been extensively characterized ^51–53^. While we were not able to probe how the S365 phospho-site affects the behavior of BRaf due to the lack of a sufficiently selective commercially available antibody, our results suggest that DFG-out-stabilizing inhibitors are not capable of efficiently engaging the ATP-binding site of CRaf that is phosphorylated at an equivalent 14-3-3-binding site (S259) (**Extended Data Fig. 6c**)^41^. We also found that pS259 CRaf was not recruited into inhibitor-promoted heterodimers and DFG-out-stabilizing inhibitor treatment did not alter pS259 CRaf levels (**Extended Data Fig. 6d**,e). Thus, our results suggest that there is a significant cellular population of autoinhibited, monomeric CRaf that is not phosphorylated at S259. We speculate that this is also true for BRaf, but this assertion remains to be validated.

We profiled how different Mek inhibitors directly affect Raf’s interactions and inhibition by DFG-out-stabilizing inhibitors by using a short co-treatment regime and concentrations of each that provided a similar level of partial downstream inhibition (**Fig. 5**). Of the clinical inhibitors we tested, cobimetinib was unique in sensitizing Raf to inhibition by LXH-254.

Cobimetinib’s ability to sensitize Raf to DFG-out-stabilizing inhibitors is consistent with its promotion of BRaf:CRaf heterodimers and disruption of high affinity Mek:Raf interactions^52^. We speculate that cobimetinib’s piperidine-derivatized 4-azetidinol moiety, which is unique amongst clinical Mek inhibitors, promotes the stabilization of a specific conformation of an α-helix in the N-terminal region of Mek’s activation loop that weakens interactions with autoinhibited Raf. This results in destabilization of the autoinhibited complex. Our demonstration that analogs of cobimetinib that provide increased disruption of high affinity Raf:Mek complexes provide greater sensitization to DFG-out-stabilizing inhibitors suggests that interactions with this region of Mek can be tuned further (**Figure 6**).

Together, our findings not only provide mechanistic insights into Raf inhibition but also suggest promising directions for developing more effective treatment strategies for cancers driven by Ras and Raf mutations. It further suggests that further optimization of Mek inhibitors as future research may focus on optimizing Mek inhibitors to enhance their synergy with DFG-out stabilizing Raf inhibitors and exploring the potential of this approach in clinical settings.

---

**Extended Data Figure 1.**
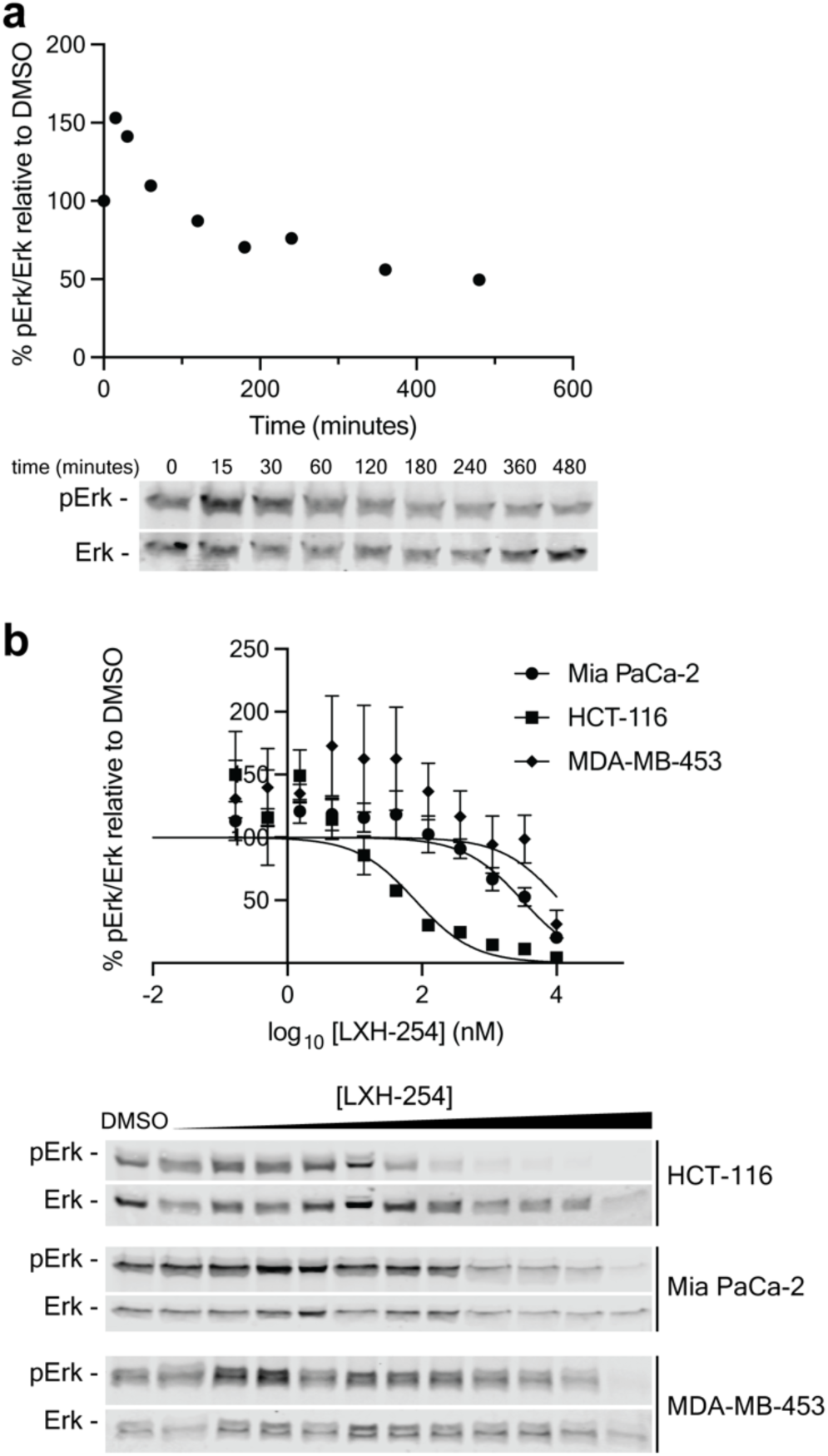
Time dependence of Raf inhibition. **a,** Western blot (*bottom*) and quantification (*top*) of percent normalized pErk levels in HCT-116 cells treated with 32 nM LXH-254 relative to DMSO for the times indicated (n=1). **b,** Representative western blots (*bottom*) showing pErk levels in HCT-116, Mia PaCa-2, and MDA-MB-453 cell lines treated with a range of LXH-254 doses for 24 h (n=3). Quantification (*top*) of normalized pErk percentages and determination of pErk IC_50_ values from the conditions described above. The percent pErk value shown at each LXH-254 concentration relative to DMSO is the mean of n=3 replicates. Error bars represent standard error of the mean (s.e.m.). IC_50_ curves were fit to percent pErk values obtained from all three replicates. pErk IC_50_ values and 95% confidence intervals (CIs) for each cell line are listed in **Table S1**.

**Extended Data Figure 2.**
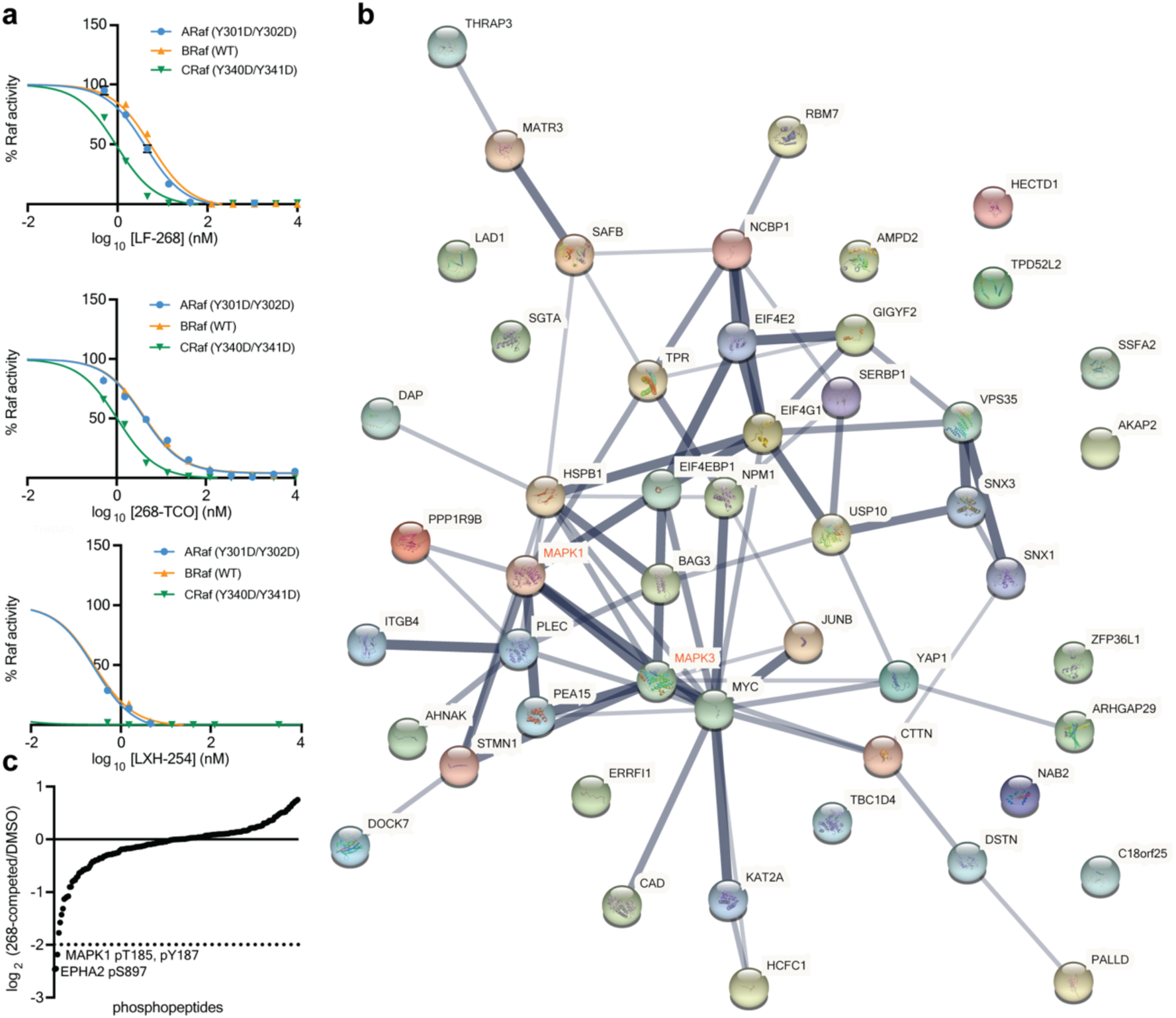
Profiling of LF-268 and 268-TCO. **a,** Percent activity values for purified Y301D/Y302D ARaf, WT BRaf, and Y340D/Y341D CRaf in the presence of a range of LF-268, 268-TCO, or LXH-254 concentrations (LF-268 n=3, 268-TCO n=1, LXH-254 n=1). For LF-268, individual data points represent the average of n=3 replicates and error bars represent standard error of the mean (s.e.m.). Calculated IC_50_ values are shown in Fig. 2a. **b,** STRING protein:protein interaction network mapping proteins with at least one phospho-site that exhibits a >2-fold average decrease in HCT-116 cells treated for 4 h with 5 μM LF-268 relative to DMSO (n=3). The thickness of the line indicates the relative strength of data that indicates a protein:protein interaction. MAPK1 and MAPK3 (Erk1 and Erk2) are labeled in red. Phospho-peptide quantifications are in **Table S3**. **c,** Histogram of log_2_ ratios of quantified phospho-peptides for HCT-116 cells treated with 5 μM of LF-268 for 4 h relative to DMSO following kinobead enrichment (n=3). Log_2_ ratios are plotted from lowest to highest and the dotted line indicates a log_2_ ratio < −2. Phospho-peptide quantifications are in **Table S4**.

**Extended Data Figure 3.**
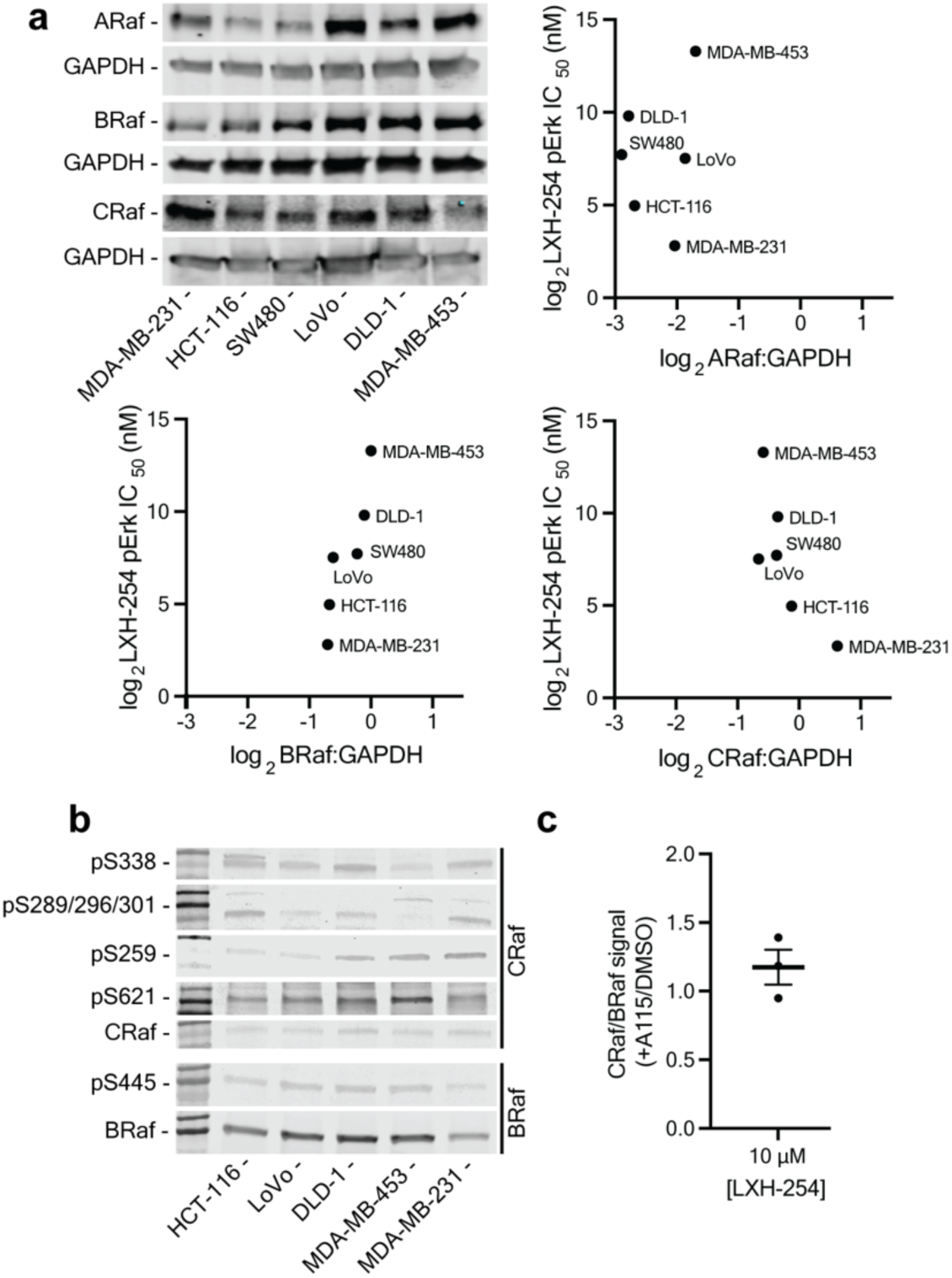
Expression and phosphorylation levels of Raf-Mek-Erk pathway components. **a,** Western blot (*top left*) of ARaf, BRaf, and CRaf expression levels across the comparative panel of mutant KRas-expressing cell lines. Correlation plots of GAPDH-normalized ARaf (*top right*), BRaf (*bottom left*), and CRaf (*bottom right*) levels as measured by western blot (n=1) versus LXH-254 pErk IC_50_ values. **b,** Western blots of phosphorylation levels of BRaf and CRaf regulatory phospho-sites (n=1). All images are from individual western blots. **c,** The ratio of CRaf co-immunoprecipitated with BRaf (normalized to immunoprecipitated BRaf) from CIAR-293 cells pre-treated with A115 relative to DMSO for 1 h, followed by incubation with LXH-254 for 3 h. CRaf co-immunoprecipitation levels for DMSO and A115 pre-treated CIAR-293 cells were probed on the same western blot, allowing relative levels to be compared. The values shown are from the experiments described in **Fig. 3h**. Bars represent the mean of n=3 replicates and error bars represent standard error of the mean (s.e.m.).

**Extended Data Figure 4.**
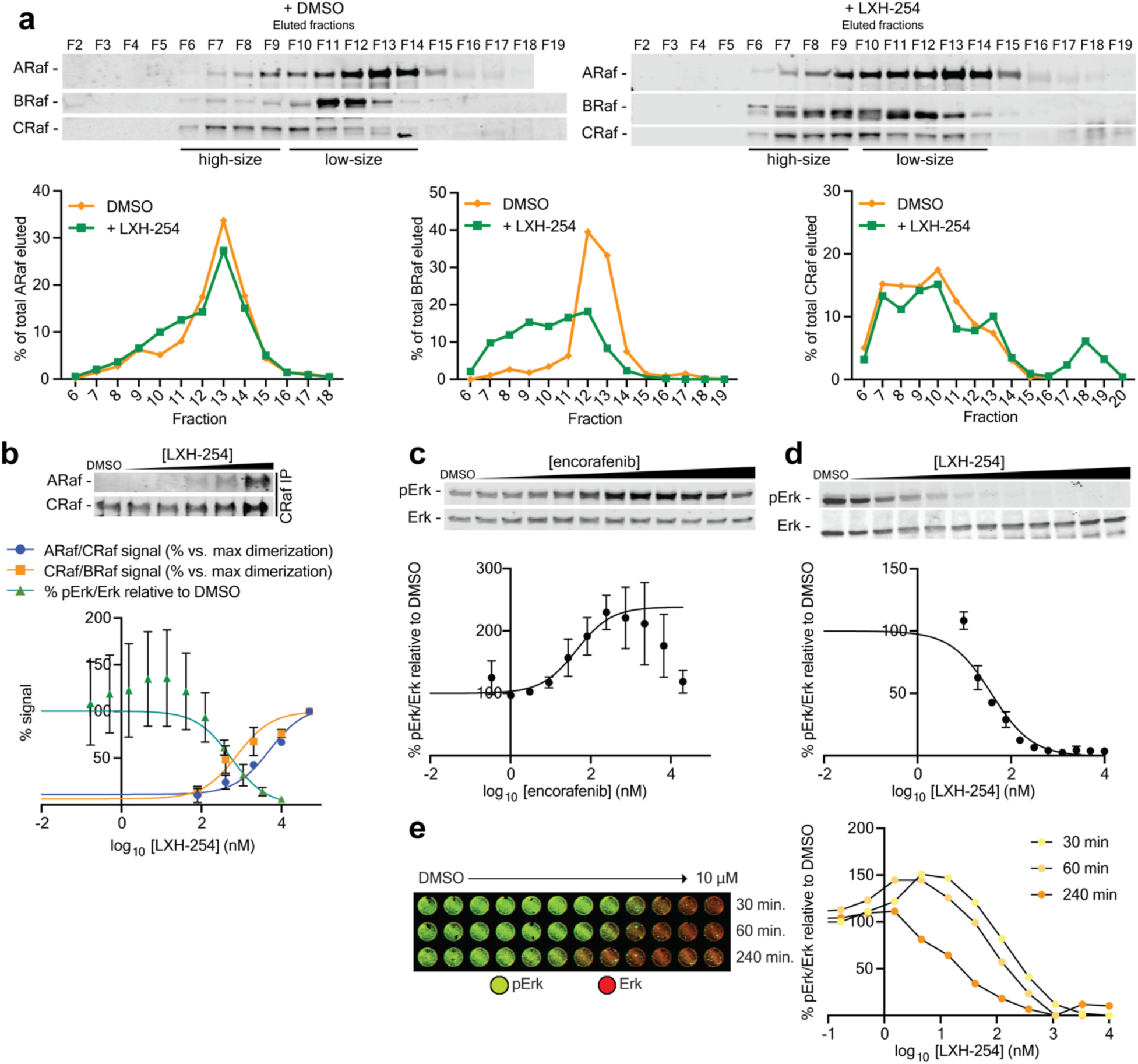
LXH-254 preferentially promotes BRaf:CRaf heterodimerization. **a,** Western blots (*top*) of ARaf, BRaf, and CRaf levels in size exclusion chromatography fractions from the lysates of Mia PaCa-2 cells treated with 10 μM LXH-254 or DMSO (n=1). Quantification indicates the amount of each Raf isoform eluted in each fraction as a percentage of the total isoform in all fractions (*bottom*). **b,** Representative western blots (*top*) and quantification (*bottom*) of ARaf levels co-immunoprecipitated with CRaf from Mia PaCa-2 treated with a range of LXH-254 doses for 3 h. The percent CRaf levels (normalized to immunoprecipitated BRaf) shown at each LXH-254 concentration are represent singlicate data. DC_50_ curves were fit to percent ARaf levels obtained from all three replicates and ARaf:CRaf dimerization DC_50_ values and 95% CIs are listed in **Table S1**. The Mia PaCa-2 pErk IC_50_ and BRaf:CRaf dimerization DC_50_ curves from **Figure 3f** are overlayed to demonstrate at which LXH-254 concentrations pErk IC_50_s and dimerization DC_50_ values intersect. Error bars represent standard error of the mean (s.e.m.). **c,** Representative western blot (*top*) of pErk levels in HCT-116 cells treated with range of encorafenib doses for 1 h followed by a 1 h washout. Quantification (bottom) of normalized pErk percentages and determination of pErk PA_50_ value. The percent pErk value shown at each LXH-254 concentration relative to DMSO is the mean of n=3 replicates. PA_50_ curves were fit to percent pErk values obtained from all replicates. pErk PA_50_ values and 95% confidence intervals (CIs) are listed in **Table S1**. **d,** Representative western blot (*top*) and quantification (bottom) of pErk levels in HCT-116 cells treated with a range of LXH-254 doses for 4 h (same data as shown in **Figure 1e,f**). **e,** In-cell western assay (*top*) of pErk levels in HCT-116 cells treated with a range of LXH-254 doses for various treatment times. Quantification (*bottom*) of normalized pErk percentages (relative to DMSO-treated cells) observed at each LXH-254 concentration from the in-cell western assay (n=1).

**Extended Data Figure 5.**
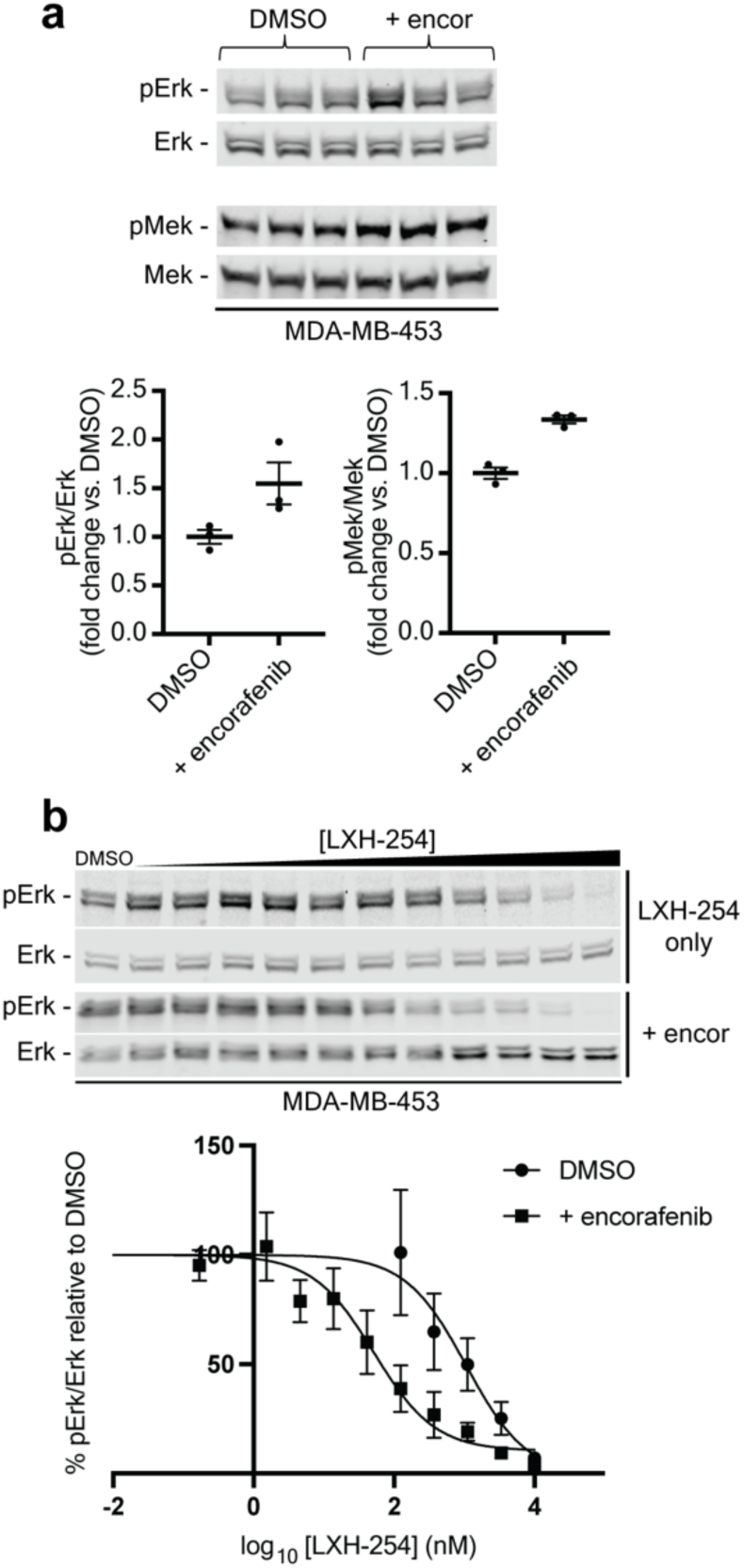
First protomer occupation sensitizes MDA-MB-453 cells to LXH-254 inhibition. **a,** Western blots (*top*) and quantification of percent pMek and pErk levels) in MDA-MB-453 cells treated with 1 μM encorafenib or DMSO for 1 h followed by washout. The percent pErk and percent pMek values (relative to DMSO-treated cells) are shown. Bars equal the mean of n=3 replicates and error bars represent standard error of the mean (s.e.m.). **b,** Representative western blots (*top*) and quantification of pErk levels in MDA-MB-453 cells pre-treated with 1 μM encorafenib for 1 h, followed by washout and subsequent incubation with a range of LXH-254 doses for 3 h. The normalized percent pErk values (relative to DMSO-treated cells) shown at each LXH-254 concentration are the mean of n=3 replicates and error bars represent s.e.m. IC_50_ curves were fit to the percent pErk values obtained from all three replicates and pErk IC_50_ values and 95% confidence intervals (CIs) are listed in **Table S1**.

**Extended Data Figure 6.**
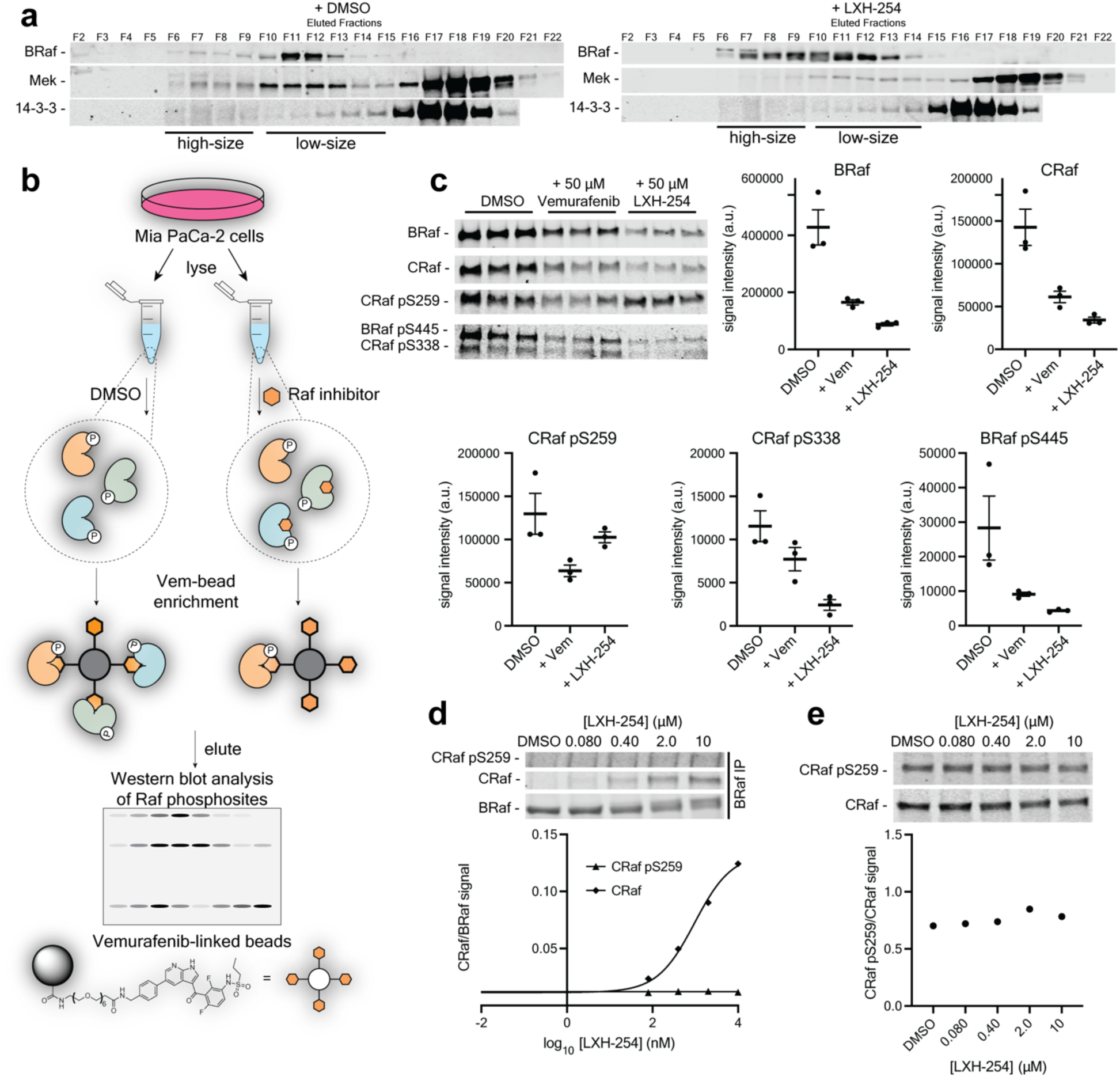
LXH-254 effectively engages non-pS259 CRaf:Mek complexes. **a,** Western blots of BRaf, Mek, and 14-3-3 levels in size exclusion chromatography fractions from the lysates of Mia PaCa-2 cells treated with 10 μM LXH-254 or DMSO (n=1). **b,** Schematic of the competitive Vemurafenib-conjugated bead enrichment to quantify inhibitor engagement of BRaf and CRaf. **c,** Representative western blots (*top left*) showing levels of BRaf, CRaf, BRaf pS445, CRaf pS338, and CRaf pS259 from Mia PaCa-2 lysates treated with Vemurafenib, LXH-254, or DMSO for 3 h and enriched with Vemurafenib-conjugated beads. Quantification of enriched BRaf, CRaf, BRaf pS445, CRaf pS338, and CRaf pS259 shown. Bars equal the mean of n=3 replicates and error bars represent standard error of the mean (s.e.m.). **d,** Western blots (*top*) and quantification (*bottom*) of CRaf and CRaf pS259 co-immunoprecipitated with BRaf from Mia PaCa-2 cells treated with a range of LXH-254 doses for 3 h. CRaf and CRaf pS259 levels were normalized to BRaf levels. **e,** Western blots (top) and quantification (bottom) of CRaf and CRaf pS259 levels in Mia PaCa-2 cells treated with a range of LXH-254 doses for 3 h (n=1).

**Extended Data Figure 7.**
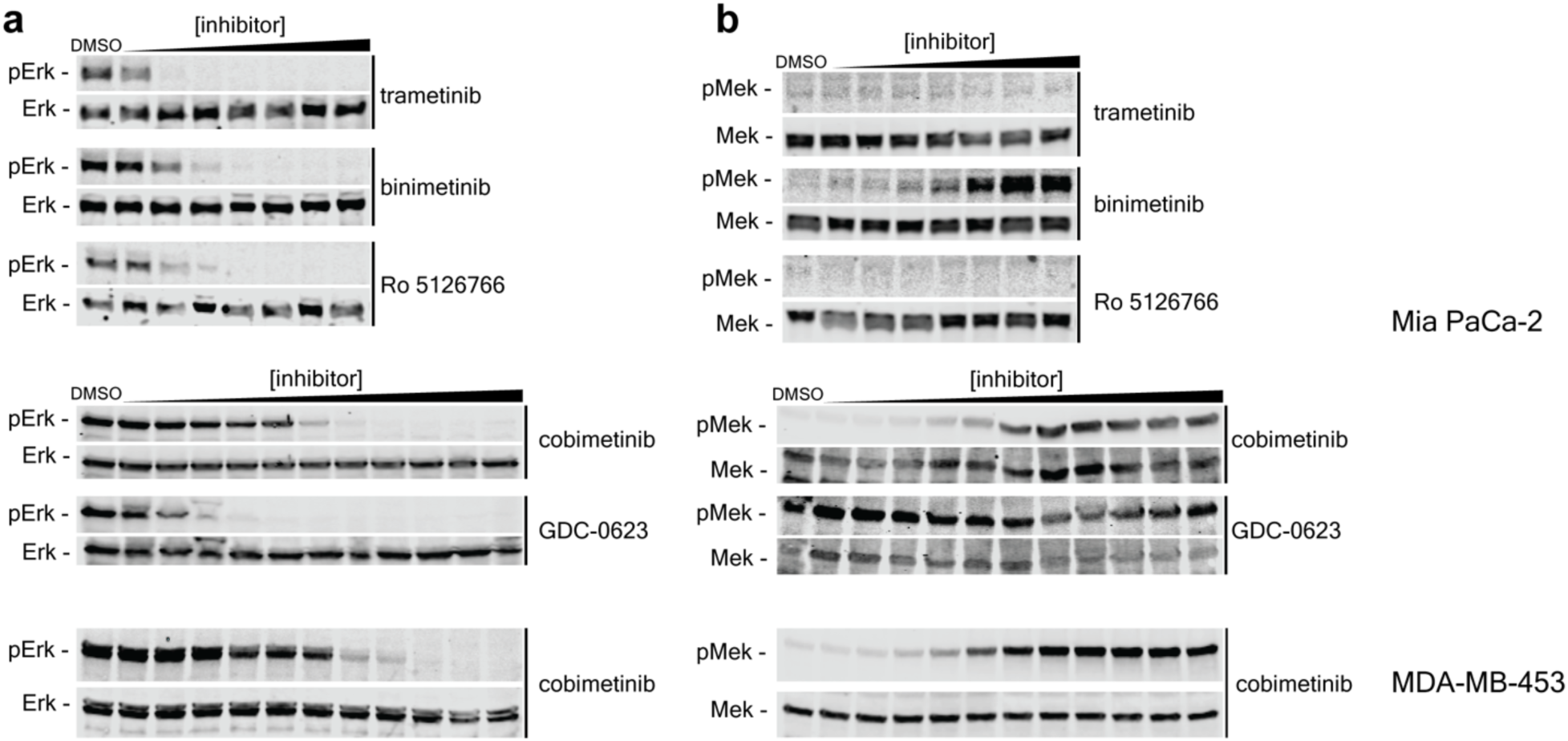
Clinical Mek inhibitor titrations in Mia PaCa-2 and MDA-MB-453 cell lines. **a and b,** Representative western blots showing pErk (**a**) and pMek (**b**) levels in Mia PaCa-2 and MDA-MB-453 cells treated with a range of Mek inhibitor doses for 4 h (n=1).

**Extended Data Figure 8.**
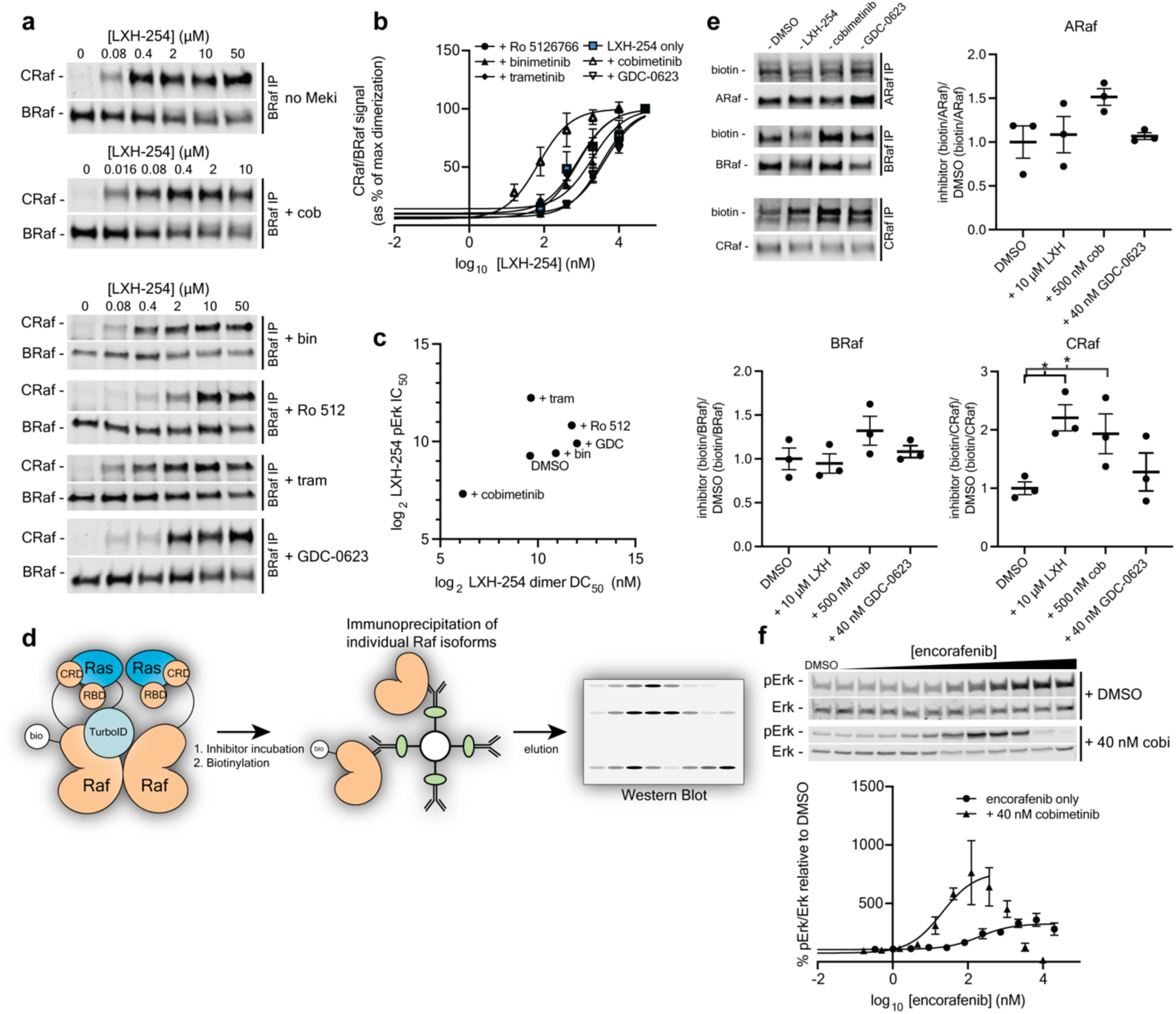
Cobimetinib facilitates LXH-254-mediated Raf dimer formation, increases Ras:Raf interactions, and sensitizes Raf to inhibition. **a and b,** Representative western blots (**a**) and quantification (**b**) of CRaf levels co-immunoprecipitated with BRaf from Mia PaCa-2 cells pre-treated with DMSO or a partial (∼85%) inhibitory dose of a Mek inhibitor for 1 h, followed by treatment with a range of LXH-254 doses for 3 h. The BRaf-normalized percent CRaf levels shown at each LXH-254 concentration relative to maximum CRaf levels are the mean of n=3 replicates and error bars represent standard error of the mean (s.e.m.). DC_50_ curves were fit to percent CRaf levels obtained from all three replicates. BRaf:CRaf dimerization DC_50_ values and 95% confidence intervals (CIs) for each cell line are listed in **Table S1**. BRaf:CRaf DC_50_ curves shown for LXH-254 only treated cells are from **Figure 3i**. Mek inhibitor pre-treatment concentrations used: cobimetinib = 200 nM, Ro 5126766 = 30 nM, binimetinib = 80 nM, trametinib = 10 nM, GDC-0623 = 10 nM. **c,** Correlation plot of LXH-254 BRaf:CRaf DC_50_s from (b) and pErk IC_50_s calculated from **Figure 5c**, d. Pearson’s *r* value = 0.595. **d,** Schematic depicting workflow for quantifying biotinylation of individual Raf isoforms by KRas-TurboID in CIAR-293 cells. **e,** Representative western blots (*top left*) showing levels of biotinylation of immunoprecipitated ARaf, BRaf, and CRaf in CIAR-293 cells transfected with a KRas-TurboID construct, treated with DMSO, LXH-254 (10 μM), cobimetinib (500 nM), or GDC-0623 (40 nM) for 4 h, followed by treatment with 100 μM biotin for 15 min. Quantified biotinylation levels of ARaf, BRaf, and CRaf (normalized to total immunoprecipitated ARaf, BRaf, and CRaf, respectively, and represented as a fraction relative to DMSO) were obtained from n=3 replicates. Bars equal the mean of all replicates and error bars represent standard error of the mean (s.e.m.). **f,** Representative western blots (*top*) of pErk levels in Mia PaCa-2 cells pre-treated with 40 nM cobimetinib or DMSO for 1 h, followed by incubation with a range of encorafenib doses for 3 h. The percent pErk values shown at each encorafenib concentration relative to DMSO are the mean of n=3 replicates. Error bars represent s.e.m. PA_50_ curves were fit to percent pErk values obtained from all three replicates. pErk PA_50_ values and 95% CIs for each cell line are listed in **Table S1**.

**Extended Data Figure 9.**
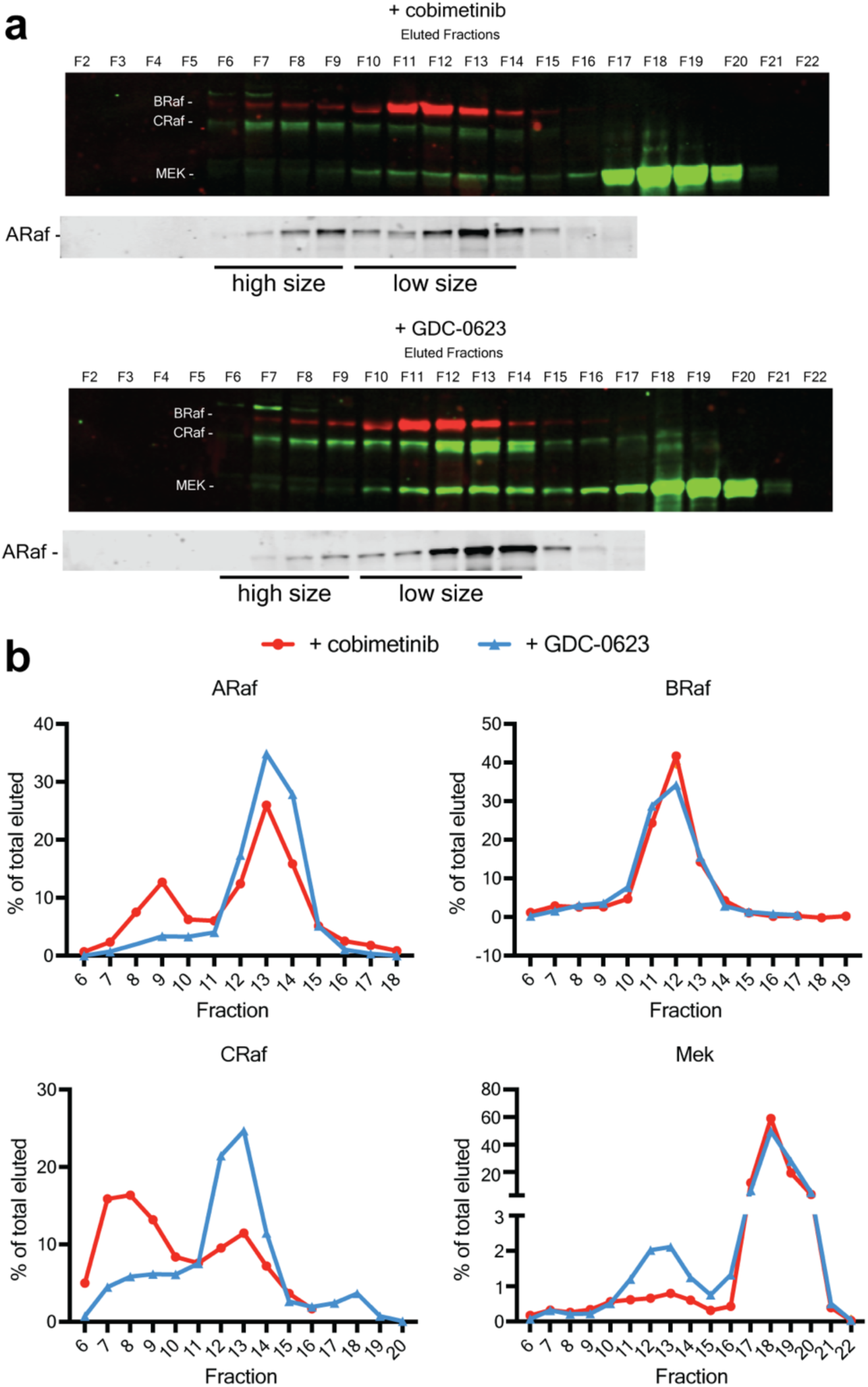
Cobimetinib destabilizes high affinity CRaf:Mek complexes. **a,** Western blots (*top*) of ARaf, BRaf, CRaf, and Mek levels in size exclusion chromatography fractions from the lysates of Mia PaCa-2 cells treated with cobimetinib or GDC-0623 (n=1). Quantification indicates the amount of each Raf isoform eluted in each fraction as a percentage of the total isoform in all fractions (*bottom*). **b,** Quantification shows the amount of each Raf isoform eluted in each fraction as a percentage of the total isoform in all fractions.

**Extended Data Figure 10.**
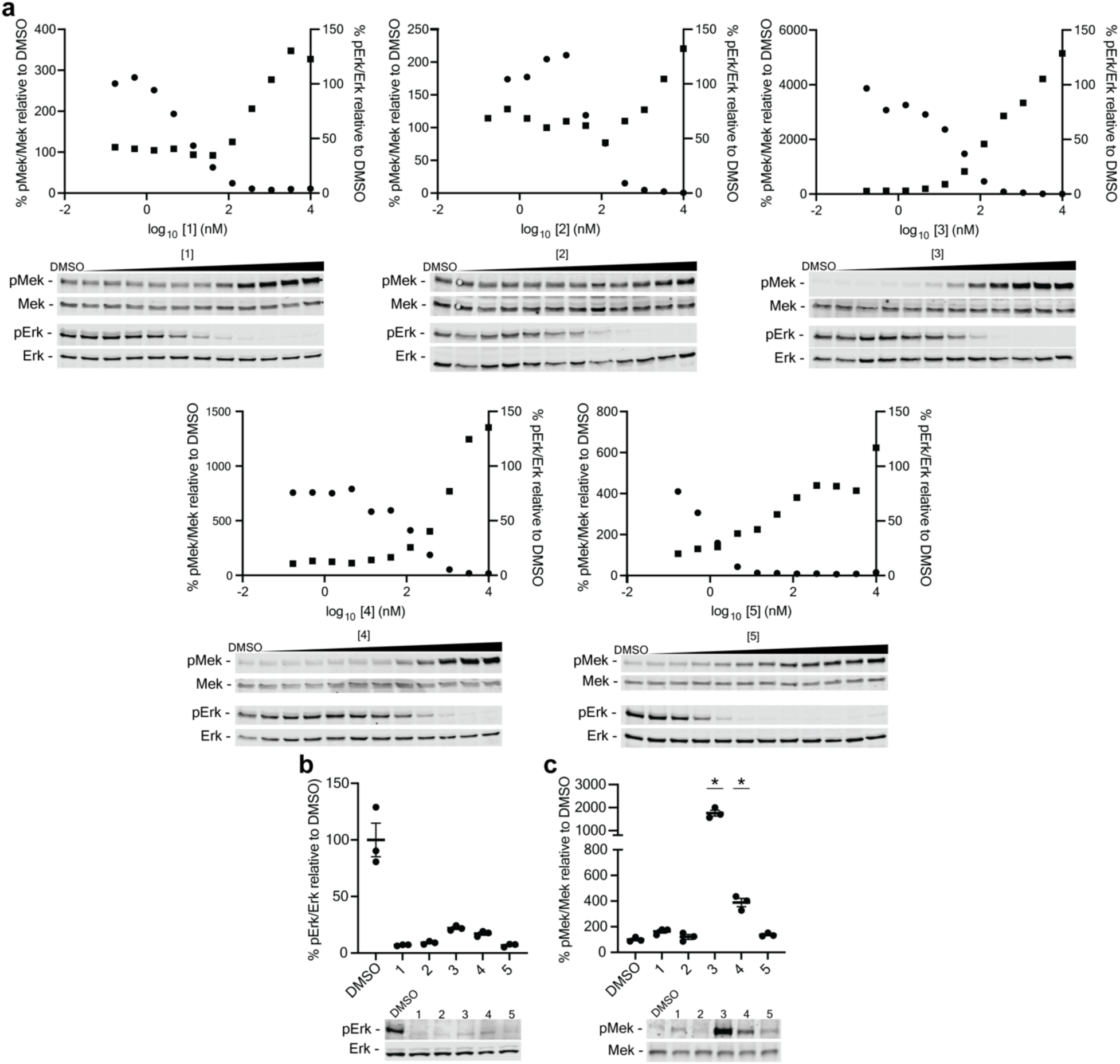
Titrations of Mek inhibitors 1-5. **a,** Western blots (*bottom*) and quantification (*top*) of pErk and pMek levels in Mia PaCa-2 cells treated with a range of **1-5** doses for 4 h (n=1). Percent pErk and pMek values were normalized to values in DMSO-treated cells. **b,** Representative western blot (*bottom*) and quantification of pErk levels in Mia PaCa-2 cells treated with DMSO or the partial inhibitory doses of **1-5** (**1** = 80 nM, **2** = 360 nM, **3** = 120 nM, **4** = 600 nM, **5** = 10 nM) that were used in **Fig. 6d, e, g, h** for 4 h (n=3). Bars equal the mean of n=3 replicates and error bars represent standard error of the mean (s.e.m.). **c,** Representative western blot (*bottom*) and quantification (*top*) of pMek levels in Mia PaCa-2 cells treated with DMSO or the partial inhibitory doses of **1-5** (**1** = 80 nM, **2** = 360 nM, **3** = 120 nM, **4** = 600 nM, **5** = 10 nM) for 4 h (n=3). Percent pErk and pMek values were normalized to values in DMSO-treated cells. Bars equal the mean of n=3 replicates and error bars represent s.e.m. *denotes significant difference between indicated samples versus DMSO-treated samples as determined by two-sided Student’s t-test (*p* value < 0.05).

## Methods

### Inhibitors

A-1155463 (ChemieTek, cat. no. CT-A115)

LXH-254 (Selleck Chemicals, cat. no. S8745)

Vemurafenib (Selleck Chemicals, cat. no. S1267)

Cobimetinib (MedChem Express, cat. no. HY-13064)

Binimetinib (MedChem Express, cat. no. HY-15202)

Ro 5126766 (MedChem Express, cat. no. HY-18652)

Trametinib (MedChem Express, cat. no. HY-10999)

Encorafenib (MedChem Express, cat. no. HY-15605)

GDC-0623 (MedChem Express, cat. no. HY-15610)

3,4-Difluoro-2-[(2-fluoro-4-iodophenyl)amino]-N-(2-hydroxyethoxy)benzamide (AABlocks, cat. no. AA01EMTY)

All commercial inhibitors were tested for purity via analytical HPLC and positively identified by mass spectrometry.

### Antibodies

MEK1 61B12 Mouse mAb (Cell Signaling Technology, cat. no. 2352)

Phospho-p44/42 MAPK (Erk1/2) D13.14.4E XP^®^Rabbit mAb (Cell Signaling Technology, cat. no. 4370)

p44/42 MAPK (Erk1/2) (3A7) Mouse mAb (Cell Signaling Technology, cat. no. 9107)

Phospho-MEK1/2 (Ser217/221) (41G9) Rabbit mAb (Cell Signaling Technology, cat. no. 9154) c-Raf (D4B3J) Rabbit mAb (Cell Signaling Technology, cat. no. 53745)

Phospho-c-Raf (Ser338) (56A6) Rabbit mAb (Cell Signaling Technology, cat. no. 9427)

Phospho-c-Raf (Ser259) Antibody (Cell Signaling Technology, cat. no. 9421)

Phospho-c-Raf (Ser289/296/301) Antibody (Cell Signaling Technology, cat. no. 9431)

Phospho-B-Raf (Ser445) Antibody (Cell Signaling Technology, cat. no. 2696)

Phospho-c-Raf (Ser621) Polyclonal Antibody (Invitrogen, cat. no. 44-504G)

Protein A Agarose Beads (Cell Signaling Technology, cat. no. 9863)

Protein G Agarose Beads (Cell Signaling Technology, cat. no. 37478)

14-3-3 (pan) E9S9M Rabbit mAb (Cell Signaling Technology, cat. no. 95422)

BRaf Antibody (F-7) (Santa Cruz Biotechnology, cat. no. 5284)

ARaf Antibody (A-5) Santa Cruz Biotechnology, cat. no. 166771)

Anti-Mek1 Antibody (Sigma-Aldrich, cat. no. 07-641)

Purified Mouse Anti-MEK1 Clone 25/MEK1 (BD Biosciences, cat. no. 610122)

GAPDH Antibody (A-3) (Santa Cruz Biotechnology, cat. no. 137179)

IRDye® 680RD Goat anti-Mouse IgG Secondary Antibody (LI-COR Biosciences, cat. no. 926-68070)

IRDye® 800CW Goat anti-Rabbit IgG Secondary Antibody (LI-COR Biosciences, cat. no. 926-32211)

#### Cell Culture

All cancer cell lines used in this study including Mia PaCa-2, MDA-MB-453, SW480, DLD-1, MDA-MB-231, LoVo, HPAF-II, AsPc-1, A549, and Calu-6 were purchased from ATCC and cultured in DMEM (ThermoFisher) + 10% FBS (Millipore Sigma) or, for HCT-116 cells, in McCoy’s media (ThermoFisher) supplemented with 10% FBS. All cell lines were maintained at 37 °C with 5% CO_2_. CIAR-293 expressing cell lines were generated as previously described.^33^

#### SDS-PAGE and Western Blotting

Gel-ready samples were quickly centrifuged and loaded onto Any kDa SDS-PAGE gels (Bio-Rad). 2 µL PAGEruler Plus Prestained Protein Ladder (Thermo Scientific) was also loaded onto each gel. SDS PAGE was performed at 200V for 40 min. Gels were then placed into Trans-Blot Turbo Nitrocellulose Midi Transfer Packs (Bio-Rad), loaded into a Trans-Blot Turbo (Bio-Rad), and protein transfer was performed (16V, 14 min). Membranes were then incubated in blocking buffer (5% dry milk in 1X TBS and 0.1% Tween-20) for 1 hour at room temperature followed by an overnight incubation at 4 °C with primary antibodies. Blots were then washed three times with TBSt (1X TBS + 0.1% Tween-20) and incubated with IRDye® 680RD Goat anti-Mouse IgG Secondary Antibody (LI-COR Biosciences) and IRDye® 800CW Goat anti-Rabbit IgG Secondary Antibody (LI-COR Biosciences) for 1 hour in blocking buffer at room temperature. Blots were washed three times with TBSt and imaged on LiCor Odyssey Dlx. Western blot analysis and quantification were done using ImageStudio software.

#### Raf inhibitor titrations in KRas mutant cell lines

Cells were plated on 12 well plates and allowed to adhere and grow overnight. Inhibitor incubations were started when cells were 80-90% confluent for 4 hour incubations or at 50% confluence for 24 hour incubations. All inhibitor titrations were done in FBS containing media. Cells were treated with LXH-254 or LF-268 (initial concentration 10 µM, 3-fold serial dilutions, final [DMSO] = 0.05%) for four or 24 hours at 37 °C. Cells were lysed using a modified RIPA buffer containing 50 mM Tris-HCl (pH = 7.5), 150 mM NaCl, 10% glycerol, 1% IGEPAL, phosphatase inhibitor cocktails 2 + 3 (Sigma Aldrich), and protease inhibitor tablet (ThermoScientific). Cells were incubated in the buffer for 30 min at 4 °C with agitation. The lysate was cleared via centrifugation at 17,000 × g for 10 min at 4 °C and the supernatant was collected. 4X SDS loading dye containing 200 mM Tris (pH 6.8), 8% SDS, 40% glycerol, 4% beta-mercaptoethanol, and 0.08% bromophenol blue was added to each sample followed by vortexing and heating at 95 °C for 5 min. Eluted proteins were subjected to SDS-PAGE and Western blot analysis using p44/42 MAPK (Erk1/2) (3A7) Mouse mAb (Cell Signaling Technology) /Phospho-p44/42 MAPK (Erk1/2) D13.14.4E XP^®^Rabbit mAb (Cell Signaling Technology) or Phospho-MEK1/2 (Ser217/221) (41G9) Rabbit mAb (Cell Signaling Technology)/Purified Mouse Anti-MEK1 Clone 25/MEK1 (BD Biosciences) antibodies at 4 °C.

### 268-TCO Pulldowns

#### Materials & Reagents

- Lysis Buffer and Washing Buffer 1: [Tris]= 50 mM, pH= 7.8; [NaCl]= 120 mM; [NaF]= 10 mM; [Na3VO4]= 1 mM; [EDTA]= 1 mM; [IGEPAL CA-630]= 1% (by volume);
- Washing Buffer 2: [Tris]= 50 mM, pH= 7.8; [NaCl]= 120 mM; [EDTA]= 1 mM. Denaturing Buffer: [Tris]= 50 mM, pH= 8.5; [Guanidinium Chloride]= 6 M; [TCEP]= 5 mM (add freshly); [CAM]= 10 mM (add freshly).
- Pierce Protease Inhibitor Tablets (Thermo Scientific; Catalog number: A32963)
- NHS-Activated Sepharose 4 Fast Flow (GE Healthcare; Catalog number: 17090601)
- Pierce 660 nm Protein Assay (Thermo Scientific)
- Lys-c Lysyl Endopeptidase, MS Grade (Wako)
- MS Grade Trypsin (Pierce)

#### Probe treatment and cell lysis

Cells were grown on 10-cm plates in Dulbecco’s Modified Eagle’s Medium (DMEM) supplemented with 10% (by volume) Fetal Bovine Serum (FBS) at 37 °C in a humidified incubator containing 5% CO2 until 90% confluence. The 268-TCO probe was added to the cells by replacing medium with fresh DMEM (5 mL) containing 10% FBS and 268-TCO (5 µM) or DMSO (DMSO%= 0.1% in the final medium). Cells were incubated with 268-TCO for 30 min at 37 °C in the incubator and the medium was removed immediately after incubation. The cells were quickly washed with ice-cold Dulbecco’s Phosphate-Buffered Saline (DPBS) (5 mL, one time) and immediately lysed with ice-cold Lysis Buffer (500 µL). Cells were scraped into pre-chilled 1.5mL Eppendorf tubes and were centrifuged at 17,000 × g for 20 min at 4 °C. The supernatant of lysates was collected, and protein concentration was measured using Pierce 660 nm Protein Assay. The protein concentration of lysates was diluted to 1.5 mg/mL with ice-cold lysis buffer.

#### Capture of probe-bound kinase complexes using Tz-Beads

Tetrazine beads were generated according to previously described protocol^54^. To reduce background contaminants, lysates were pre-cleared by incubating with 15 µL (volume of drained beads) of sepharose beads coupled to ethanolamine for 1 hour at 4 °C prior to incubation with Tz-beads. For each pulldown experiment, 15 µL (volume of drained beads) of Tz-Beads (PMID: 31274292) were added to a 1.5-mL Eppendorf tube and were washed with ice-cold lysis buffer (three times; 400 µL for each wash). 750 µg of protein lysates (1.5 mg/mL) was added to Tz-Beads (15 µL) and was gently incubated on an end-to-end rotator for 30 mins at 4 °C. After incubation with Tz-Beads, the flow-through was quickly removed by aspiration. The beads were then washed with ice-cold Washing Buffer 1 by gently inverting the tube four times (three times; 400 µL for each wash), followed by washing with Washing Buffer 2 by gently inverting the tube for four times (three times; 400 µL for each wash). The target protein should be enriched and bound to the Tz-Beads by the end of this step.

#### On-bead tryptic digestion of protein targets for tandem mass spectrometry analysis

To detect enriched targets using LC/MS, the washed beads were resuspended in 50 µL of Denaturing Buffer and were boiled at 95 °C for 5 min. After cooling down, 100 mM TEAB (50 µL) was added. The pH value was adjusted to 8 to 9 by adding NaOH (1 M). Lys-c Lysyl Endopeptidase (1 µg) was then added and incubated with the beads at 37 °C for 2 hours in a thermal mixer agitating at 1,400 rpm. After incubation, the suspension was diluted with 100 mM TEAB (100 µL) and MS Grade Trypsin (1 µg) was added. pH was adjusted to 8 to 9. The beads were incubated with trypsin overnight at 37 °C in a thermal mixer agitating at 1,400 rpm. After incubation, Buffer A (200 µL) was added to the beads and trypsin was quenched by adjusting the S58 pH to 2 to 3 using formic acid. The resulting supernatant was desalted on C18 StageTip and washed once by buffer A (50 µL). The tryptic peptides were eluted off the C18 StageTip using buffer B (50 µL). The eluted solution was dried in speed-vac and then reconstituted with buffer A (20 µL) for LC/MS analysis.

#### Proteomic Data Acquisition

Peptides were separated on a nanoAcquity UPLC instrument with 12 cm long fused silica capillary column (Polymicro Technologies Flexible Fused Silica Capillary Tubing, Inner Diameter 75 µm, Outer Diameter 375 µm, cat. no. 1068150019) made in-house with a laser puller and packed with 5 µm 120 Å reversed phase C18 beads (ReproSil-Pur 120 C18-AQ, Dr. Maisch GmbH HPLC, cat. no. r15.aq.). The LC gradient was 90 min long with 10-35% B at 200 nL/min. LC solvent A was 0.1% acetic acid and LC solvent B was 0.1% acetic acid, 99.9% acetonitrile. MS data were collected with a Thermo Scientific Orbitrap Fusion Tribrid mass spectrometer.

#### Proteomic Data Processing

Raw data were analyzed using MaxQuant and search engine Andromeda using the following default settings: Multiplicity was set to 1; Digestion was set to trypsin/P; Variable modifications were set to Oxidation (M) and Acetyl (Protein N-term); LFQ minimal ratio count was set to 2; LFQ minimal number of neighbors set to 3; LFQ average number of neighbors set to 6; MS/MS spectra were searched against the UniProt human database (updated July 22nd, 2015); False discovery rate (FDR) was set to 0.01; First search and MS/MS search mass tolerance was set to 10 ppm; The data in the Protein Groups result table was processed with Perseus software. MaxQuant intensity values of identified proteins were used for all downstream analysis. We only consider proteins identified if MaxQuant was able to compute corresponding protein intensity values. Identified proteins that were classified as “only identified by site”, “reverse”, or “contaminant” were filtered and removed from the table. Proteins with 1 or fewer peptide counts were filtered and removed. All intensity values were transformed based on log2 algorithm. Missing protein intensity values for identified proteins were imputed using Perseus by sampling from a distribution downshifted by 1.3 and having a width of 0.25. The *P*-values were calculated using two-tailed, two-sample *t*-test. Enrichment scores (log_2_[intensity (268-TCO)/intensity (DMSO)]) were calculated using the relative intensity values from probe-treated and DMSO-treated samples with confidence values (log_10_(P-value)) for each identified protein. We defined significant targets as protein hits with an enrichment score ≥ 1.0 and a confidence value ≥ 1.4.

### Kinobead Enrichment

#### Materials & Reagents

- Lysis Buffer and Washing Buffer 1: [Tris]= 50 mM, pH= 7.8; [NaCl]= 120 mM; [NaF]= 10 mM; [Na3VO4]= 1 mM; [EDTA]= 1 mM; [IGEPAL CA-630]= 1% (by volume);
- Washing Buffer 2: [Tris]= 50 mM, pH= 7.8; [NaCl]= 120 mM; [EDTA]= 1 mM. Denaturing Buffer: [Tris]= 50 mM, pH= 8.5; [Guanidinium Chloride]= 6 M; [TCEP]= 5 mM (add freshly); [CAM]= 10 mM (add freshly).
- Pierce Protease Inhibitor Tablets (Thermo Scientific; Catalog number: A32963)
- NHS-Activated Sepharose 4 Fast Flow (GE Healthcare; Catalog number: 17090601)
- Pierce 660 nm Protein Assay (Thermo Scientific)
- Lys-c Lysyl Endopeptidase, MS Grade (Wako)
- MS Grade Trypsin (Pierce)

#### Generation of kinobeads

Kinobeads were generated as previously described^31^. 150 μL of cellular lysates (in mod. RIPA buffer) were incubated with kinobeads for 3 hours at 4 °C. Beads were then washed sequentially with twice with cold Mod. RIPA buffer and three times with cold 1X TBS. Proteins on beads were then denatured by resuspending in denaturing buffer (6M guanidinium chloride, 50 mM Tris, 5 mM TCEP, 10 mM CAM) and heating at 95 °C for 5 min. The slurry was then diluted with 100 mM TEAB to a final concentration of 50 mM. After adjusting pH to 8-9, LysC was added to the beads and incubated at 37 °C for 2 hours. Samples were diluted 2x with 100mM TEAB and incubated overnight with trypsin. The resulting peptides were subjected to a C18 cleanup, dried, and resuspended in 80:20 ACN:water mixture and subsequently analyzed by LC-MS/MS.

#### Preparation of cell lysates

In a 10-cm plate, cells were washed with ice-cold DPBS for three times and lysed with 500 µL of ice-cold mod RIPA buffer ([Tris]=50 mM, [NaCl]=150 mM, [NaF]=10 mM, [Igepal-CA630]=1.0%, [Sodium deoxycholate]= 0.25%, [glycerol]=5%, pH=7.8, containing protease inhibitor, 1 mM PMSF, and phosphatase inhibitor cocktail 2&3). Cell lysates were centrifuged at 17,000 × g for 10 min at 4 °C. The supernatant was collected and quantified using Pierce 660 nm Protein Assay Reagent and diluted to 2.0 mg/mL with ice-cold lysis buffer.

#### Inhibitor treatment and kinase enrichment with kinobeads

The lysates (160 µL) were then incubated DMSO or 10 µM of inhibitor for 30 min on an end-to-end rotator at 4°C. Meanwhile, 10 µL of a 50% kinobeads slurry was washed with 500 µL of mod RIPA buffer twice. The inhibitor-treated lysates were then added to the drained kinobeads and incubated on an end-to-end rotator for 3 hours at 4°C. After incubation, the beads were spun down and the supernatant was aspirated off. The beads were then washed with 500 µL of ice-cold mod RIPA buffer twice and with 500 µL of ice cold TBS buffer ([Tris]=50 mM, pH= 7.8, [NaCl]=150 mM) for three times to remove the detergent.

#### On-bead trypsinization and LC/MS sample preparation

The washed beads were then resuspended in 25 µL of denaturing buffer ([TFE]=20%, [Tris]=5 mM, pH=8.5, [TCEP]=5 mM (add freshly), [CAM]=10 mM (add freshly)). The beads slurry was vortexed briefly and then boiled at 95 °C for 5 min (release pressure frequently). The beads slurry was diluted with 25 µL of 10% TFE in water. The pH of the resulting suspension was adjusted to 8-9 by adding 1 µL of 1N NaOH at a time. 0.4 µg of LysC was added to the slurry and agitated on a thermomixer at 37 °C for 2 hours at 1,400 rpm. The slurry was then diluted with 50 µL of 10% TFE. 0.4 µg of sequencing grade trypsin was added to the slurry and the sample was agitated overnight at 37 °C at 800 rpm on a thermomixer. After trypsinization, the slurry was diluted with 100 µL of Buffer A (5% acetonitrile in water, 0.1% TFA) containing 1% formic acid. The resulting slurry was agitated at room temperature for 5 min. The resulting supernatant was then loaded onto a C-18 column made in-house, washed with 50 µL of Buffer A, and eluted to a low-retention Eppendorf tube using 50 µL of Buffer B (80% acetonitrile in water, 0.1% TFA). The peptides were then resuspended in 20 µL of Buffer A to render the final samples.

#### IMAC Enrichment

Fe^3+^-NTA loaded beads were washed with 10 bead volumes of 80% aqueous acetonitrile with 0.1% TFA. 20 µL peptides (in 80% aq. acetonitrile with 0.1% TFA) were incubated with the beads and phosphopeptides were enriched. Phosphopeptides were eluted twice with 50 mM KH_2_PO_4_/NH_4_OH (pH=10), combined, pH neutralized, and dried by vacuum centrifugation. The phosphopeptides were then subjected to a C18 cleanup, dried, and resuspended in 80:20 ACN:water mixture followed by LC-MS/MS.

#### Proteomic Data Acquisition

Peptides were separated on a nanoAcquity UPLC instrument with a 12-cm long fused silica capillary column (Polymicro Technologies Flexible Fused Silica Capillary Tubing, Inner Diameter 75 µm, Outer Diameter 375 µm, cat. no. 1068150019) made in-house with a laser puller and packed with 5 µm 120 Å reversed phase C18 beads (ReproSil-Pur 120 C18-AQ, Dr. Maisch GmbH HPLC, cat. no. r15.aq.). The LC gradient was 90-min long with 10-35% solvent B at 200 nL/min. LC solvent A was 0.1% acetic acid in water. LC solvent B was 0.1% acetic acid in acetonitrile. MS data were collected with a Thermo Scientific Orbitrap Fusion Tribrid mass spectrometer.

#### Proteomic Data Process and Analysis

The protocol was modified based on the protocol previously described in the supporting information of Journal of the American Chemical Society, 141(30), 11912-11922. Raw datasets were analyzed using MaxQuant and search engine Andromeda using the following default settings: Multiplicity was set to 1; Digestion was set to trypsin/P; Variable modifications were set to Oxidation (M) and Acetyl (Protein N-term); LFQ minimal ratio count was set to 2; LFQ minimal number of neighbors was set to 3; LFQ average number of neighbors was set to 6; MS/MS spectra were searched against the UniProt human database (updated July 22nd, 2015); False discovery rate (FDR) was set to 0.01; First search and MS/MS search mass tolerance was set to 10 ppm; The data in the Protein Groups result table was processed with Perseus software. The missing protein intensity values in LFQ analyses were imputed by sampling from a distribution downshifted by 1.3 with a width of 0.2. The imputed LFQ intensity values of identified proteins were used for all downstream analyses. We only consider proteins as quantified if MaxQuant was able to compute corresponding protein intensity values in DMSO-treated samples. Identified proteins that were classified as “only identified by site”, “reverse”, or “contaminant” were filtered and removed from the table. Proteins with one or fewer peptide counts were filtered and removed. All intensity values were transformed based on log2 algorithm. The *P*-values were calculated using two-tailed, two-sample *t*-tests. Competition scores (log_2_R)= (log_2_[Intensity(DMSO-treated)/Intensity(Inhibitor-treated)]) were calculated using the relative intensity values from DMSO- and inhibitor-treated lysates with confidence values=log_10_(*P*-value) for each identified protein. We defined significantly competed kinases as protein hits with a competition score ≥ 1.5 and a confidence value ≥ 1.5 from a two-sample t test with an FDR of 0.05 (n = 3).

#### Quantifying relative expression and phosphorylation status of MAPK pathway proteins in KRas mutant cell lines

KRas mutant cell lines were plated onto 6-well plates and allowed to adhere and grow overnight. The next morning cells were lysed using a modified RIPA buffer containing 50 mM Tris-HCl (pH = 7.5), 150 mM NaCl, 10% glycerol, 1% IGEPAL, phosphatase inhibitor cocktails 2 + 3 (Sigma Aldrich), and protease inhibitor tablet (ThermoScientific). Cells were incubated in the buffer for 30 min at 4 °C with agitation. The lysate was transferred to eppendorf tubes and cleared via centrifugation at 17,000 × g for 10 min at 4 °C and the supernatant was collected. 4X SDS loading dye containing 200 mM Tris (pH 6.8), 8% SDS, 40% glycerol, 4% beta-mercaptoethanol, and 0.08% bromophenol blue was added to each sample followed by vortexing and heating at 95 °C for 5 min. Eluted proteins were subjected to SDS-PAGE and Western blot analysis using ARaf Antibody (A-5) (Santa Cruz Biotechnology), BRaf Antibody (F-7) (Santa Cruz Biotechnology), c-Raf (D4B3J) Rabbit mAb (Cell Signaling Technology), GAPDH Mouse mAb (A-3) (Santa Cruz Biotechnology), Phospho-c-Raf (Ser338) (56A6) Rabbit mAb (Cell Signaling Technology), Phospho-c-Raf (Ser259) Antibody (Cell Signaling Technology), Phospho-c-Raf (Ser289/296/301) Antibody (Cell Signaling Technology), Phospho-B-Raf (Ser445) Antibody (Cell Signaling Technology), Phospho-c-Raf (Ser621) Polyclonal Antibody (Invitrogen), p44/42 MAPK (Erk1/2) (3A7) Mouse mAb (Cell Signaling Technology), Phospho-p44/42 MAPK (Erk1/2) D13.14.4E XP^®^Rabbit mAb (Cell Signaling Technology), Phospho-MEK1/2 (Ser217/221) (41G9) Rabbit mAb (Cell Signaling Technology), and Purified Mouse Anti-MEK1 Clone 25/MEK1 (BD Biosciences).

#### Generating CIAR-293 cell line

A CIAR construct containing the HVR of KRas 4a was generated in pcDNA5/FRT/TO (ThermoFisher). The construct was integrated into Flp-In T-Rex 293 cells. Transfection was done using turbofectin (Origene) followed by hygromycin selection (ThermoFisher) for 7-10 days.

#### Inhibitor titrations in CIAR-293 cells

CIAR-293 cells were plated on poly-d-lysine coated plates in DMEM + 10% FBS and allowed to grow and adhere overnight. Cells were then serum starved for 18 hours followed by a 1 hour incubation with 100 nM A-1155463 or DMSO (0.05%) (ChemieTek) at 37 °C. Cells were then exposed to LXH-254 (initial concentration 10 µM, 3-fold serial dilutions) in the presence of 100 nM A-115463 for 3 hours at 37 °C. Cells were lysed using a modified RIPA buffer containing 50 mM Tris-HCl (pH = 7.5), 150 mM NaCl, 10% glycerol, 1% IGEPAL, phosphatase inhibitor cocktails 2 + 3 (Sigma Aldrich), and protease inhibitor tablet (ThermoScientific). Cells were incubated in the buffer for 30 min at 4 °C with agitation. The lysate was transferred to eppendorf tubes and cleared via centrifugation at 17,000 × g for 10 min at 4 °C and the supernatant was collected. 4X SDS loading dye containing 200 mM Tris (pH 6.8), 8% SDS, 40% glycerol, 4% beta-mercaptoethanol, and 0.08% bromophenol blue was added to each sample followed by vortexing and heating at 95 °C for 5 min. Eluted proteins were subjected to SDS-PAGE and Western blot analysis using a combination of p44/42 MAPK (Erk1/2) (3A7) Mouse mAb (Cell Signaling Technology) /Phospho-p44/42 MAPK (Erk1/2) D13.14.4E XP^®^Rabbit mAb (Cell Signaling Technology) or Phospho-MEK1/2 (Ser217/221) (41G9) Rabbit mAb (Cell Signaling Technology)/Purified Mouse Anti-MEK1 Clone 25/MEK1 (BD Biosciences) antibodies.

#### Co-immunoprecipitation of CRaf with BRaf in CIAR-293 cells

CIAR-293 cells were plated on 6-well poly-d-lysine coated plates in DMEM + 10% FBS and allowed to grow and adhere overnight. Cells were then serum starved for 18 hours followed by a 1 hour incubation with 100 nM A-1155463 (ChemieTek) or DMSO (0.05%). Cells were then exposed to LXH-254 (initial concentration 50 µM, 5-fold serial dilutions) for 3 hours at 37 °C. Cells were lysed using a modified RIPA buffer containing 50 mM Tris-HCl (pH = 7.5), 150 mM NaCl, 10% glycerol, 1% IGEPAL, phosphatase inhibitor cocktails 2 + 3 (Sigma Aldrich), and protease inhibitor tablet (ThermoScientific). Cells were incubated in the buffer for 30 min at 4 °C with agitation. The lysate was transferred to eppendorf tubes and cleared via centrifugation at 17,000 × g for 10 min at 4 °C and the supernatant was collected. Meanwhile, Protein G agarose beads (Cell Signaling Technology) were loaded with BRaf Antibody (F-7, Santa Cruz Biotechnology), with the following protocol: beads were quickly washed by gentle centrifugation three times with modified RIPA buffer on ice. Beads were then resuspended in mod. RIPA buffer, antibody (5 µg antibody/20 µL 50% bead slurry) was added and incubated with beads for 2.5 hours at 4 °C with end-over-end rotation. Beads were quickly washed with mod. RIPA buffer followed by the addition of 200 µL of cell lysate (1 mg/mL protein concentration) and incubation at 4 °C for 2 hours with end-over-end rotation. Beads were quickly washed three times with mod. RIPA buffer and proteins were eluted with 1X SDS loading dye containing 50 mM Tris (pH 6.8), 2% SDS, 10% glycerol, 1% beta-mercaptoethanol, and 0.02% bromophenol blue. Eluted proteins were subjected to SDS-PAGE and Western blot analysis using BRaf Antibody (F-7, Santa Cruz Biotechnology) and c-Raf (D4B3J) Rabbit mAb (Cell Signaling Technology).

#### Co-immunoprecipitation of CRaf with BRaf in KRas mutant cells after LXH-254 incubation

MiaPaCa-2 cells were plated on 6 well plates in DMEM + 10% FBS at 75% confluence and allowed to adhere and grow overnight. HCT-116 cells were plated on 6 well plates in McCoy’s 5A Medium at 50% confluence and allowed to adhere and grow overnight. Cells were then exposed to LXH-254 (in MiaPaCa-2 cells: initial concentration 50 µM, 5-fold serial dilutions; in HCT-116 cells: initial concentration 10 µM, 5-fold serial dilutions) for 4 hours at 37 °C. Cells were lysed using a modified RIPA buffer containing 50 mM Tris-HCl (pH = 7.5), 150 mM NaCl, 10% glycerol, 1% IGEPAL, phosphatase inhibitor cocktails 2 + 3 (Sigma Aldrich), and protease inhibitor tablet (ThermoScientific). Cells were incubated in the buffer for 30 min at 4 °C with agitation. The lysate was cleared via centrifugation at 17,000 × g for 10 min at 4 °C and the supernatant was collected. Meanwhile, Protein G agarose beads (Cell Signaling Technology) were loaded with BRaf Antibody (F-7, Santa Cruz Biotechnology), with the following protocol: beads were quickly washed by gentle centrifugation three times with modified RIPA buffer on ice. Beads were then resuspended in mod. RIPA buffer, antibody (5 µg antibody/20 µL 50% bead slurry) was added and incubated with beads for 2.5 hours at 4 °C with end-over-end rotation. Beads were quickly washed once with mod. RIPA buffer, followed by the addition of 200 µL of cell lysate. Samples were then incubated at 4 °C for 2 hours with end-over-end rotation. Beads were quickly washed three times with mod. RIPA buffer via centrifugation and proteins were eluted with 1X SDS loading dye containing 50 mM Tris (pH 6.8), 2% SDS, 10% glycerol, 1% beta-mercaptoethanol, and 0.02% bromophenol blue. Eluted proteins were subjected to SDS-PAGE and Western blot analysis using BRaf Antibody (F-7, Santa Cruz Biotechnology), c-Raf (D4B3J) Rabbit mAb (Cell Signaling Technology), and Phospho-c-Raf (Ser259) Antibody (Cell Signaling Technology).

#### Size Exclusion Chromatography

Mia PaCa-2 cells were plated on 15 cm plates and allowed to adhere and grow overnight. For live cell treatments, cells were incubated with DMSO, LXH-254 (10 µM), cobimetinib (500 nM), or GDC-0623 (40 nM) at indicated concentrations for 4 hours at 37 °C. Cells were then lysed with mod. RIPA, transferred to eppendorf tubes, and cleared via centrifugation at 17,000 × g for 10 min at 4 °C and the supernatant was collected. Proteins in lysates were then separated using size exclusion chromatography with a Superdex 200 Increase 10/300 GL column on an FPLC. The elution buffer contained 20 mM Tris (pH 8.0), 150 mM NaCl, and 10% glycerol. Buffer was pumped through the column at 0.5 mL/min for 60 min with fractions collected every min. Samples were then collected, 4X SDS loading dye containing 200 mM Tris (pH 6.8), 8% SDS, 40% glycerol, 4% beta-mercaptoethanol, and 0.08% bromophenol blue was added, and heated at 95 °C for 5 min. Samples were then subjected to SDS-PAGE and Western blotting using ARaf Antibody (A-5) (Santa Cruz Biotechnology), BRaf Antibody (F-7, Santa Cruz Biotechnology), c-Raf (D4B3J) Rabbit mAb (Cell Signaling Technology), 14-3-3 (pan) E9S9M Rabbit mAb (Cell Signaling Technology), and Purified Mouse Anti-MEK1 Clone 25/MEK1 (BD Biosciences).

#### Co-immunoprecipitation of ARaf with CRaf in MiaPaCa-2 cells treated with LXH-254 dilution

MiaPaCa-2 cells were plated on 6 well plates in DMEM + 10% FBS at 75% confluence and allowed to adhere and grow overnight. Cells were then exposed to LXH-254 (initial concentration 50 µM, 5-fold serial dilutions) for 4 hours at 37 °C. Cells were lysed using a modified RIPA buffer containing 50 mM Tris-HCl (pH = 7.5), 150 mM NaCl, 10% glycerol, 1% IGEPAL, phosphatase inhibitor cocktails 2 + 3 (Sigma Aldrich), and protease inhibitor tablet (ThermoScientific). Cells were incubated in the buffer for 30 min at 4 °C with agitation. The lysate was transferred to eppendorf tubes and cleared via centrifugation at 17,000 × g for 10 min at 4 °C and the supernatant was collected. Meanwhile, Protein A agarose beads (Cell Signaling Technology) were loaded with c-Raf (D4B3J) Rabbit mAb (Cell Signaling Technology), with the following protocol: beads were quickly washed by gentle centrifugation three times with modified RIPA buffer on ice. Beads were then resuspended in mod. RIPA buffer, antibody (10 µg antibody/20 µL 50% bead slurry) was added and incubated with beads for 2.5 hours at 4 °C with end-over-end rotation. Beads were quickly washed followed by the addition of 200 µL of cell lysate. Samples were then incubated at 4 °C for 2 hours with end-over-end rotation. Beads were quickly washed three times with mod. RIPA buffer via centrifugation and proteins were eluted with 1X SDS loading dye containing 50 mM Tris (pH 6.8), 2% SDS, 10% glycerol, 1% beta-mercaptoethanol, and 0.02% bromophenol blue. Eluted proteins were subjected to SDS-PAGE and Western blot analysis using ARaf Antibody (A-5) (Santa Cruz Biotechnology) and c-Raf (D4B3J) Rabbit mAb (Cell Signaling Technology).

#### Encorafenib titrations in KRas mutant cell lines

HCT-116 cells were plated at 50% confluence on 12 well plates in McCoy’s 5A Medium + 10% FBS and allowed to grow and adhere overnight. MiaPaCa-2 cells were plated at 75% confluence on 12 well plates in DMEM + 10% FBS and allowed to grow and adhere overnight. Cells were then exposed to encorafenib (initial concentration 20 µM, 3-fold serial dilutions) for 4 hours at 37 °C. Cells were lysed using a modified RIPA buffer containing 50 mM Tris-HCl (pH = 7.5), 150 mM NaCl, 10% glycerol, 1% IGEPAL, phosphatase inhibitor cocktails 2 + 3 (Sigma Aldrich), and protease inhibitor tablet (ThermoScientific). Cells were incubated in the buffer for 30 min at 4 °C with agitation. The lysate was cleared via centrifugation at 17,000 × g for 10 min at 4 °C and the supernatant was collected. 4X SDS loading dye containing 200 mM Tris (pH 6.8), 8% SDS, 40% glycerol, 4% beta-mercaptoethanol, and 0.08% bromophenol blue was added to each sample followed by vortexing and heating at 95 °C for 5 min. Samples were then subjected to SDS-PAGE and Western blot analysis using a combination of p44/42 MAPK (Erk1/2) (3A7) Mouse mAb (Cell Signaling Technology)/Phospho-p44/42 MAPK (Erk1/2) D13.14.4E XP^®^Rabbit mAb (Cell Signaling Technology) or Phospho-MEK1/2 (Ser217/221) (41G9) Rabbit mAb (Cell Signaling Technology)/Purified Mouse Anti-MEK1 Clone 25/MEK1 (BD Biosciences).

#### In cell western protocol

HCT-116 cells were plated on 96 well plates in McCoy’s 5A Medium + 10% FBS at 50% confluence and allowed to grow and adhere overnight. Cells were then exposed to LXH-254 (initial concentration 10 µM, 3-fold serial dilutions) for 30, 60, and 240 min at 37 °C. Media was aspirated and replaced with 4% PFA in PBS solution and incubated at room temperature for 25 min. Samples were then incubated with 125 mM glycine in 1X PBS for 5 min followed by blocking with Intercept PBS Blocking Buffer (LicorBio) for 1 hour at 37 °C. p44/42 MAPK (Erk1/2) (3A7) Mouse mAb (Cell Signaling Technology)/Phospho-p44/42 MAPK (Erk1/2) D13.14.4E XP^®^Rabbit mAb (Cell Signaling Technology) were diluted in Intercept PBS Block Buffer (LicorBio) and added to the samples followed by incubation for 1 hour at room temperature. Samples were then washed three times with 1X PBS + 0.1% Tween-20 followed by addition of IRDye® 680RD Goat anti-Mouse IgG Secondary Antibody (LI-COR Biosciences) and IRDye® 800CW Goat anti-Rabbit IgG Secondary Antibody (LI-COR Biosciences) and incubation for 1 hour in blocking buffer at room temperature. Samples were then washed three times with 1X PBS + 0.1% Tween-20. Plates were then imaged on LiCor Odyssey Dlx.

#### Encorafenib and LXH-254 co-treatment protocol

HCT-116 cells were plated at 50% confluence on 12 well plates in McCoy’s 5A Medium + 10% FBS and allowed to grow and adhere overnight. MiaPaCa-2 cells and MDA-MB-453 cells were plated at 75% confluence on 12 well plates in DMEM + 10% FBS and allowed to grow and adhere overnight. Cells were then exposed to encorafenib (1 µM) for 1 hour at 37 °C followed by replacement with LXH-254 (initial concentration 10 µM, 3-fold serial dilutions) and incubation for 3 additional hours at 37 °C. Cells were lysed using a modified RIPA buffer containing 50 mM Tris-HCl (pH = 7.5), 150 mM NaCl, 10% glycerol, 1% IGEPAL, phosphatase inhibitor cocktails 2 + 3 (Sigma Aldrich), and protease inhibitor tablet (ThermoScientific). Cells were incubated in the buffer for 30 min at 4 °C with agitation. The lysate was cleared via centrifugation at 17,000 × g for 10 min at 4 °C and the supernatant was collected. 4X SDS loading dye containing 200 mM Tris (pH 6.8), 8% SDS, 40% glycerol, 4% beta-mercaptoethanol, and 0.08% bromophenol blue was added to each sample followed by vortexing and heating at 95 °C for 5 min. Samples were then subjected to SDS-PAGE and Western blot analysis using a combination of p44/42 MAPK (Erk1/2) (3A7) Mouse mAb (Cell Signaling Technology) and Phospho-p44/42 MAPK (Erk1/2) D13.14.4E XP^®^Rabbit mAb (Cell Signaling Technology).

#### Co-immunoprecipitation of BRaf, CRaf, and CRaf pS259 with Mek in MiaPaCa-2 cells treated with LXH-254

MiaPaCa-2 cells were plated on 6 well plates in DMEM + 10% FBS at 75% confluence and allowed to adhere and grow overnight. Cells were then exposed to LXH-254 (10 µM) for 4 hours at 37 °C. Cells were lysed using a modified RIPA buffer containing 50 mM Tris-HCl (pH = 7.5), 150 mM NaCl, 10% glycerol, 1% IGEPAL, phosphatase inhibitor cocktails 2 + 3 (Sigma Aldrich), and protease inhibitor tablet (ThermoScientific). Cells were incubated in the buffer for 30 min at 4 °C with agitation. The lysate was cleared via centrifugation at 17,000 × g for 10 min at 4 °C and the supernatant was collected. Meanwhile, Protein A agarose beads (Cell Signaling Technology) were loaded with Anti-Mek1 Antibody (Sigma-Aldrich), with the following protocol: beads were quickly washed by gentle centrifugation three times with modified RIPA buffer on ice. Beads were then resuspended in mod. RIPA buffer, antibody (10 µg antibody/20 µL 50% bead slurry) was added and incubated with beads for 2.5 hours at 4 °C with end-over-end rotation. Beads were quickly washed followed by the addition of 200 µL of cell lysate. Samples were then incubated at 4 °C for 2 hours with end-over-end rotation. Beads were quickly washed three times with mod. RIPA buffer via centrifugation and proteins were eluted with 1X SDS loading dye containing 50 mM Tris (pH 6.8), 2% SDS, 10% glycerol, 1% beta-mercaptoethanol, and 0.02% bromophenol blue. Eluted proteins were subjected to SDS-PAGE and Western blot analysis using BRaf Antibody (F-7) (Santa Cruz Biotechnology), c-Raf (D4B3J) Rabbit mAb (Cell Signaling Technology), Phospho-c-Raf (Ser259) Antibody (Cell Signaling Technology), and Purified Mouse Anti-MEK1 Clone 25/MEK1 (BD Biosciences).

#### Vemurafenib Bead Generation

1 mL NHS-ester Sepharose 4 FastFlow beads were washed three times with a 5 mL 1:1 mixture of DMF and ethanol on a vacuum manifold. 2.5 mg of Compound 6 (see synthetic methods) and 250 µL DIPEA were dissolved in 500 µL of a 1:1 DMF:ethanol mixture and added to the beads. The conjugation reaction was incubated at room temperature overnight with end-over-end rotation. The next morning beads were washed three times with 5 mL DMF:ethanol and resuspended in 500 DMF:ethanol containing 1M EDCI-HCl. 100 µL ethanolamine and 250 µL DIPEA were then added to the mixture and the reaction was allowed to proceed overnight with end-over-end rotation. Beads were then washed three times with 2 mL 1:1 DMF:ethanol, three times with 2 mL 0.5 M NaCl in water, three times with 2 mL 20% ethanol in water, and were resuspended with 500 µL 20% ethanol in water and stored at 4 °C until use.

#### Vemurafenib-bead Pulldowns

HCT-116 cells were plated on a 15 cm plate and allowed to grow to confluence and adhere overnight. Cells were lysed with mod. RIPA buffer. 200 µL 1 mg/mL lysate was aliquoted into separate eppies. Samples were then incubated with indicated concentrations of LXH-254, Vemurafenib, or DMSO for 1 hour at 4 °C with end-over-end rotation. Meanwhile, vemurafenib beads were washed three times with mod. RIPA. Inhibitor-incubated lysate was then added to 20 µL 50% vemurafenib bead slurry for 3 hours at 4 °C with end-over-end rotation. Beads were then washed three times with ice cold lysis buffer and eluted by incubating with 30 µL 1X SDS loading dye at 95 °C for 5 min. Eluted proteins were subjected to SDS-PAGE and Western blot analysis using BRaf Antibody (F-7) (Santa Cruz Biotechnology), Phospho-B-Raf (Ser445) Antibody (Cell Signaling Technology) c-Raf (D4B3J) Rabbit mAb (Cell Signaling Technology), Phospho-c-Raf (Ser259) Antibody (Cell Signaling Technology), and Phospho-c-Raf (Ser338) (56A6) Rabbit mAb (Cell Signaling Technology).

#### Mek inhibitor titrations in MiaPaCa-2 and MDA-MB-453 cells

MiaPaCa-2 or MDA-MB-453 cells were plated at 75% confluence on 12 well plates in DMEM + 10% FBS and allowed to grow and adhere overnight. Cells were then exposed to cobimetinib, trametinib, binimetinib, Ro 5126766, GDC-0623, **1**, **2**, **3**, **4**, and **5** (initial concentration 10 µM, 3-fold serial dilutions) for 4 hours at 37 °C. Cells were lysed using a modified RIPA buffer containing 50 mM Tris-HCl (pH = 7.5), 150 mM NaCl, 10% glycerol, 1% IGEPAL, phosphatase inhibitor cocktails 2 + 3 (Sigma Aldrich), and protease inhibitor tablet (ThermoScientific). Cells were incubated in the buffer for 30 min at 4 °C with agitation. The lysate was cleared via centrifugation at 17,000 × g for 10 min at 4 °C and the supernatant was collected. 4X SDS loading dye containing 200 mM Tris (pH 6.8), 8% SDS, 40% glycerol, 4% beta-mercaptoethanol, and 0.08% bromophenol blue was added to each sample followed by vortexing and heating at 95 °C for 5 min. Samples were then subjected to SDS-PAGE and Western blot analysis using a combination of p44/42 MAPK (Erk1/2) (3A7) Mouse mAb (Cell Signaling Technology)/Phospho-p44/42 MAPK (Erk1/2) D13.14.4E XP^®^Rabbit mAb (Cell Signaling Technology) or Phospho-MEK1/2 (Ser217/221) (41G9) Rabbit mAb (Cell Signaling Technology)/Purified Mouse Anti-MEK1 Clone 25/MEK1 (BD Biosciences).

#### LXH-254 and Mek inhibitor co-treatments in MiaPaCa-2 or MDA-MB-453 cells

MiaPaCa-2 cells were plated at 75% confluence on 12 well plates in DMEM + 10% FBS and allowed to grow and adhere overnight. Cells were then pretreated with cobimetinib, trametinib, binimetinib, Ro 5126766, GDC-0623, **1**, **2**, **3**, **4**, and **5** for 1 hour at 37 °C followed by addition of LXH-254 (initial concentration 10 µM, 3-fold serial dilutions) and incubation of 3 additional hours at 37 °C. Cells were lysed using a modified RIPA buffer containing 50 mM Tris-HCl (pH = 7.5), 150 mM NaCl, 10% glycerol, 1% IGEPAL, phosphatase inhibitor cocktails 2 + 3 (Sigma Aldrich), and protease inhibitor tablet (ThermoScientific). Cells were incubated in the buffer for 30 min at 4 °C with agitation. The lysate was cleared via centrifugation at 17,000 × g for 10 min at 4 °C and the supernatant was collected. 4X SDS loading dye containing 200 mM Tris (pH 6.8), 8% SDS, 40% glycerol, 4% beta-mercaptoethanol, and 0.08% bromophenol blue was added to each sample followed by vortexing and heating at 95 °C for 5 min. Samples were then subjected to SDS-PAGE and Western blot analysis using a combination of p44/42 MAPK (Erk1/2) (3A7) Mouse mAb (Cell Signaling Technology) and Phospho-p44/42 MAPK (Erk1/2) D13.14.4E XP^®^Rabbit mAb (Cell Signaling Technology).

#### Bliss Synergy Analysis

Bliss synergy analysis was performed from cobimetinib/LXH-254 combination treatment pErk titration data using SynergyFinder 3.0, a free web-based application used to analyze multi-dose drug combinations^55^.

#### Co-immunoprecipitation of CRaf with BRaf in MiaPaCa-2 cells after Mek inhibitor and LXH-254 co-treatments

MiaPaCa-2 cells were plated on 6 well plates in DMEM + 10% FBS at 75% confluence and allowed to adhere and grow overnight. Cells were then treated with Mek inhibitors (cobimetinib, binimetinib, trametinib, Ro 5126766, GDC-0623) for 1 hour at 37 °C followed by the addition of LXH-254 (initial concentration 50 µM, 5-fold serial dilutions; for cobimetinib treated samples: initial concentration 10 µM, 5-fold serial dilutions) and incubation for an additional 3 hours at 37 °C. Cells were lysed using a modified RIPA buffer containing 50 mM Tris-HCl (pH = 7.5), 150 mM NaCl, 10% glycerol, 1% IGEPAL, phosphatase inhibitor cocktails 2 + 3 (Sigma Aldrich), and protease inhibitor tablet (ThermoScientific). Cells were incubated in the buffer for 30 min at 4 °C with agitation. The lysate was cleared via centrifugation at 17,000 × g for 10 min at 4 °C and the supernatant was collected. Meanwhile, Protein G agarose beads (Cell Signaling Technology) were loaded with BRaf Antibody (F-7, Santa Cruz Biotechnology), with the following protocol: beads were quickly washed by gentle centrifugation 3 times with modified RIPA buffer on ice. Beads were then resuspended in mod. RIPA buffer, antibody (5 µg antibody/20 µL 50% bead slurry) was added and incubated with beads for 2.5 hours at 4 °C with end-over-end rotation. Beads were quickly washed followed by the addition of 200 µL of cell lysate. Samples were then incubated at 4 °C for 2 hours with end-over-end rotation. Beads were quickly washed three times with mod. RIPA buffer via centrifugation and proteins were eluted with 1X SDS loading dye containing 50 mM Tris (pH 6.8), 2% SDS, 10% glycerol, 1% beta-mercaptoethanol, and 0.02% bromophenol blue. Eluted proteins were subjected to SDS-PAGE and Western blot analysis using BRaf Antibody (F-7, Santa Cruz Biotechnology) and c-Raf (D4B3J) Rabbit mAb (Cell Signaling Technology).

#### KRas-TurboID labeling and immunoprecipitations

A plasmid containing a KRas-TurboID fusion construct was transiently transfected into a 15 cm plate of CIAR-293 cells at 75% confluence. After 24 hours, cells were then transferred to poly d-lysine coated 6-well plates at 50% confluence and allowed to grow and adhere overnight. Cells were then treated with A115 (100 nM) and DMSO (0.05% w/v), LXH-254 (10 µM), cobimetinib (500 nM), or GDC-0623 (40 nM) for 4 hours at 37 °C, followed by the addition of biotin (100 µM) and an additional 15 min incubation at 37 °C. Cells were lysed using a modified RIPA buffer containing 50 mM Tris-HCl (pH = 7.5), 150 mM NaCl, 10% glycerol, 1% IGEPAL, phosphatase inhibitor cocktails 2 + 3 (Sigma Aldrich), and protease inhibitor tablet (ThermoScientific). Cells were incubated in the buffer for 30 min at 4 °C with agitation. The lysate was cleared via centrifugation at 17,000 × g for 10 min at 4 °C and the supernatant was collected. Meanwhile, Protein G agarose beads (Cell Signaling Technology) were loaded with BRaf Antibody (F-7, Santa Cruz Biotechnology) or ARaf Antibody (A-5) Santa Cruz Biotechnology) and Protein A agarose beads were loaded with c-Raf (D4B3J) Rabbit mAb (Cell Signaling Technology) using the following protocol: beads were quickly washed by gentle centrifugation three times with modified RIPA buffer on ice. Beads were then resuspended in mod. RIPA buffer, antibody (10 µg antibody/20 µL 50% bead slurry) was added and incubated with beads for 2.5 hours at 4 °C with end-over-end rotation. Beads were quickly washed three times with mod. RIPA buffer via centrifugation followed by the addition of 200 µL cell lysates. Beads were quickly washed three times with mod. RIPA buffer via centrifugation followed by elution with 1X SDS loading dye containing 50 mM Tris (pH 6.8), 2% SDS, 10% glycerol, 1% beta-mercaptoethanol, and 0.02% bromophenol blue. Eluted proteins were subjected to SDS-PAGE and Western blot analysis using ARaf Antibody (A-5) (Santa Cruz Biotechnology), BRaf Antibody (F-7) (Santa Cruz Biotechnology) and c-Raf (D4B3J) Rabbit mAb (Cell Signaling Technology).

#### Encorafenib titration in presence of cobimetinib in MiaPaCa-2 cells

MiaPaCa-2 cells were plated at 75% confluence on 12 well plates in DMEM + 10% FBS and allowed to grow and adhere overnight. Cells were then pretreated with cobimetinib for 1 hour at 37 °C followed by addition of encorafenib (initial concentration 20 µM (for encorafenib only samples; 10 µM for cobimetinib pretreated samples), with 3-fold serial dilutions) and incubation of 3 additional hours at 37 °C. Cells were lysed using a modified RIPA buffer containing 50 mM Tris-HCl (pH = 7.5), 150 mM NaCl, 10% glycerol, 1% IGEPAL, phosphatase inhibitor cocktails 2 + 3 (Sigma Aldrich), and protease inhibitor tablet (ThermoScientific). Cells were incubated in the buffer for 30 min at 4 °C with agitation. The lysate was cleared via centrifugation at 17,000 × g for 10 min at 4 °C and the supernatant was collected. 4X SDS loading dye containing 200 mM Tris (pH 6.8), 8% SDS, 40% glycerol, 4% beta-mercaptoethanol, and 0.08% bromophenol blue was added to each sample followed by vortexing and heating at 95 °C for 5 min. Samples were then subjected to SDS-PAGE and Western blot analysis using a combination of p44/42 MAPK (Erk1/2) (3A7) Mouse mAb (Cell Signaling Technology) and Phospho-p44/42 MAPK (Erk1/2) D13.14.4E XP^®^Rabbit mAb (Cell Signaling Technology).

#### Co-immunoprecipitation of BRaf and CRaf with Mek in presence of cobimetinib and new Mek inhibitors

MiaPaCa-2 cells were plated on 6 well plates in DMEM + 10% FBS at 75% confluence and allowed to adhere and grow overnight. Cells were then exposed to cobimetinib, **1**, **2**, **3**, **4**, and **5** for 4 hours at 37 °C. Cells were lysed using a modified RIPA buffer containing 50 mM Tris-HCl (pH = 7.5), 150 mM NaCl, 10% glycerol, 1% IGEPAL, phosphatase inhibitor cocktails 2 + 3 (Sigma Aldrich), and protease inhibitor tablet (ThermoScientific). Cells were incubated in the buffer for 30 min at 4 °C with agitation. The lysate was cleared via centrifugation at 17,000 × g for 10 min at 4 °C and the supernatant was collected. Meanwhile, Protein A agarose beads (Cell Signaling Technology) were loaded with Anti-Mek1 Antibody (Sigma-Aldrich), with the following protocol: beads were quickly washed by gentle centrifugation three times with modified RIPA buffer on ice. Beads were then resuspended in mod. RIPA buffer, antibody (10 µg antibody/20 µL 50% bead slurry) was added and incubated with beads for 2.5 hours at 4 °C with end-over-end rotation. Beads were quickly washed followed by the addition of 200 µL of cell lysate. Samples were then incubated at 4 °C for 2 hours with end-over-end rotation. Beads were quickly washed three times with mod. RIPA buffer via centrifugation and proteins were eluted with 1X SDS loading dye containing 50 mM Tris (pH 6.8), 2% SDS, 10% glycerol, 1% beta-mercaptoethanol, and 0.02% bromophenol blue. Eluted proteins were subjected to SDS-PAGE and Western blot analysis using BRaf Antibody (F-7) (Santa Cruz Biotechnology) and c-Raf (D4B3J) Rabbit mAb (Cell Signaling Technology).

#### GraphPad Prism curve fitting

All curve fits associated with IC_50_s, DC_50_s, CC_50_s and PA_50_s were fit using GraphPad Prism software. The curve fit model chosen was “one site - fit logIC_50_” with a least squares regression fitting method. For pErk and pMek IC50s, only values at or under 100% were included for the curve fitting. In this context, the most relevant attribute of the inhibitors was their ability to inhibit downstream signaling. Capturing the paradoxical activation window and including these values during curve fitting drastically worsens the fit and makes the data difficult to interpret.

#### Statistical Analyses

With the exception of proteomics data, student’s t-tests and Pearson’s correlations were performed using GraphPad Prism software. For the proteomics data reported, Student’s t-tests were performed using Microsoft Excel or Perseus.

#### Software used

Software used for data analysis includes STRING, ChemDraw, GraphPad Prism, Adobe Illustrator, ChemDraw, Prism, ImageStudio, PyMol, Microsoft Excel, Perseus, MaxQuant, SynergyFinder, and Mnova.

### Synthesis Methods

All chemicals purchased from commercial suppliers were used without further purification unless otherwise stated. Reactions were monitored with thin-layer chromatography (TLC) using silica gel 60 F254 coated glass plates (EM Sciences). Compound purification was performed with a Combi-Flash Rf + (Teledyne Isco) automated flash chromatography system using pre-packed Redi-Sep Rf silica columns (Hexane/EtOAc or CH2Cl2/MeOH gradient solvent). An Agilent Eclipse XDB C18 column (250 mm x 21.2 mm), eluting with H2O/CH3CN or H2O/MeOH gradient solvent (+0.05% TFA), was used for preparatory HPLC purification. The purity of all final compounds was determined by analytical HPLC with a Varian Microsorb-MV 100-5 C18 column (4.6 mm x 150 mm), eluting with either H2O/CH3CN or H2O/MeOH gradient solvent (+0.05% TFA). Elution was monitored by a UV detector at l= 220 nm and l= 254 nm, with all final compounds displaying >95% purity. Nuclear magnetic resonance (NMR) spectra were recorded on Bruker 300 or 500 MHz NMR spectrometers at ambient temperature. Chemical shifts were reported in parts per million (ppm) and coupling constants in hertz (Hz). 1H-NMR spectra were referenced to the residual solvent peaks as internal standards (7.26 ppm for CDCl3 and and 3.34 ppm for CD3OD). Mass spectra were recorded with a Bruker Esquire Liquid Chromatograph - Ion Trap Mass Spectrometer.

**Synthesis of Vemurafenib-NH_2_**

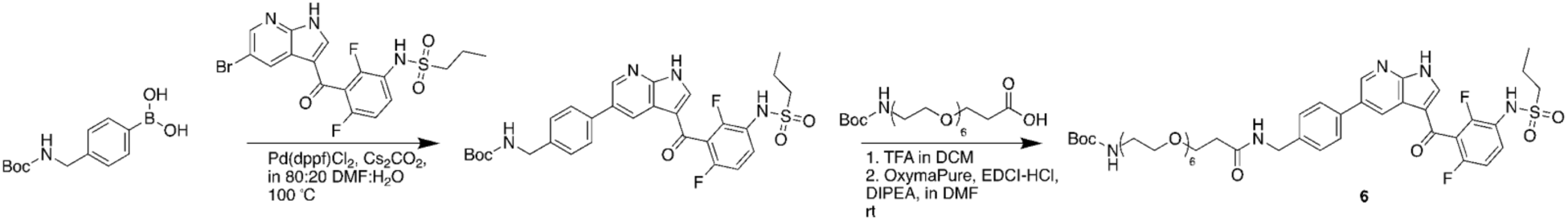

**Synthesis of carbamic acid, *N*-[[4-[3-[2,6-difluoro-3-[(propylsulfonyl)amino]benzoyl]-1*H*-pyrrolo[2,3-*b*]pyridin-5-yl]phenyl]methyl]-, 1,1-dimethylethyl ester**

To a solution of 327 mg *C*-(1,1-Dimethylethyl) *N*-[(4-boronophenyl)methyl]carbamate (Combi-Blocks) (1.30 mmol) in 5 mL 20% water in DMF in a microwave vial, 500 mg 1-propanesulfonamide, *N*-[3-[(5-bromo-1*H*-pyrrolo[2,3-*b*]pyridin-3-yl)carbonyl]-2,4-difluorophenyl] (AABlocks) (1.09 mmol) and 975 mg cesium carbonate (3.27 mmol) was added and dissolved. The resulting solution was bubbled with nitrogen for 5 minutes, following which 40 mg [1,1-Bis(diphenylphosphino)ferrocene]dichloropalladium(II) (0.055 mmol) (Sigma Aldrich) was added. The solution was microwaved (Biotage) for 10 hours at 100 °C. After cooling to room temperature, 50 mL ethyl acetate was added to the solution. The ethyl acetate solution was washed three times with brine, following which the extract was collected and dried over Na_2_SO_4_. The product was purified via silica gel chromatography (EtOAc:hexanes gradient) (RediSep Flash Chromatograph), yielding 167 mg carbamic acid, *N*-[[4-[3-[2,6-difluoro-3-[(propylsulfonyl)amino]benzoyl]-1*H*-pyrrolo[2,3-*b*]pyridin-5-yl]phenyl]methyl]-, 1,1-dimethylethyl ester (0.29 mmol) with a 26.2% yield. The product was dissolved in 3 mL 30% TFA in DCM and allowed to react for 2 hours with stirring. The solution was then co-evaporated with toluene three times via rotary evaporation. ^1^H NMR (after boc deprotection) (300 MHz, MeOD) δ 8.86 (s, 1H), 8.57 (s, 1H), 7.89 – 7.44 (m, 6H), 7.39 (s, 3H), 7.04 (t, *J* = 8.1 Hz, 1H), 4.11 (s, 3H), 3.17 – 2.90 (m, 2H), 2.03 – 1.74 (m, 2H), 1.02 (t, *J* = 7.5 Hz, 3H). MS (ESI, m/z) calculated mass = 584.19, [M+H]^+^ found 585.2.

**Synthesis of 20-{*N*-[(4-{3-[2,6-difluoro-3-(propylsulfonylamino)benzoyl]-1,7-diaza-1*H*-inden-5-yl}phenyl)methyl]carbamoyl}-3,6,9,12,15,18-hexaoxaicosyl 2-methylpropane-2-carbamate (6)**

20 mg *N*-[[4-[3-[2,6-difluoro-3-[(propylsulfonyl)amino]benzoyl]-1*H*-pyrrolo[2,3-*b*]pyridin-5-yl]phenyl]methyl]-, 1,1-dimethylethyl ester (0.034 mmol) was dissolved in 1 mL dry DMF followed by addition of 15.4 mg DIPEA (0.119 mmol) and stirring for five minutes. 18.6 mg 2,2-Dimethyl-4-oxo-3,8,11,14,17,20,23-heptaoxa-5-azahexacosan-26-oic acid (BroadPharm) (0.041 mmol) and 5.8 mg OxymaPure (0.041 mmol) (Sigma Aldrich) were then added to the reaction mixture and dissolved. Finally, 7.9 mg EDCI-HCl (0.041 mmol) (Sigma Aldrich) was added and the reaction was stirred overnight at room temperature. The next morning, product was extracted using 50 mL ethyl acetate washed twice with saturated ammonium chloride, twice with sodium bicarbonate, and twice with brine. The extract was dried over Na_2_SO_4_, decanted, evaporated, and purified via HPLC (acetonitrile:water gradient) followed by lyophilization yielding 6.4 mg 20-{*N*-[(4-{3-[2,6-difluoro-3-(propylsulfonylamino)benzoyl]-1,7-diaza-1*H*-inden-5-yl}phenyl)methyl]carbamoyl}-3,6,9,12,15,18-hexaoxaicosyl 2-methylpropane-2-carbamate (6) (20.3% yield). The product was dissolved in 3 mL 30% TFA in DCM and allowed to react for 2 hours with stirring. The solution was then co-evaporated with toluene three times via rotary evaporation. ^1^H NMR (after boc deprotection) (300 MHz, MeOD) δ 8.78 (s, 1H), 8.67 (d, *J* = 2.2 Hz, 1H), 8.01 (s, 1H), 7.89 – 7.64 (m, 3H), 7.49 (d, *J* = 7.8 Hz, 2H), 7.18 (t, *J* = 8.7 Hz, 1H), 4.51 (s, 2H), 3.92 – 3.53 (m, 24H), 3.23 – 2.96 (m, 4H), 2.59 (t, *J* = 5.8 Hz, 2H), 1.88 (q, *J* = 7.6 Hz, 2H), 1.06 (t, *J* = 7.5 Hz, 3H). MS (ESI, m/z) calculated mass = 819.33, [M+H]^+^ found 820.5.

**Synthesis of Mek inhibitors 1-4**

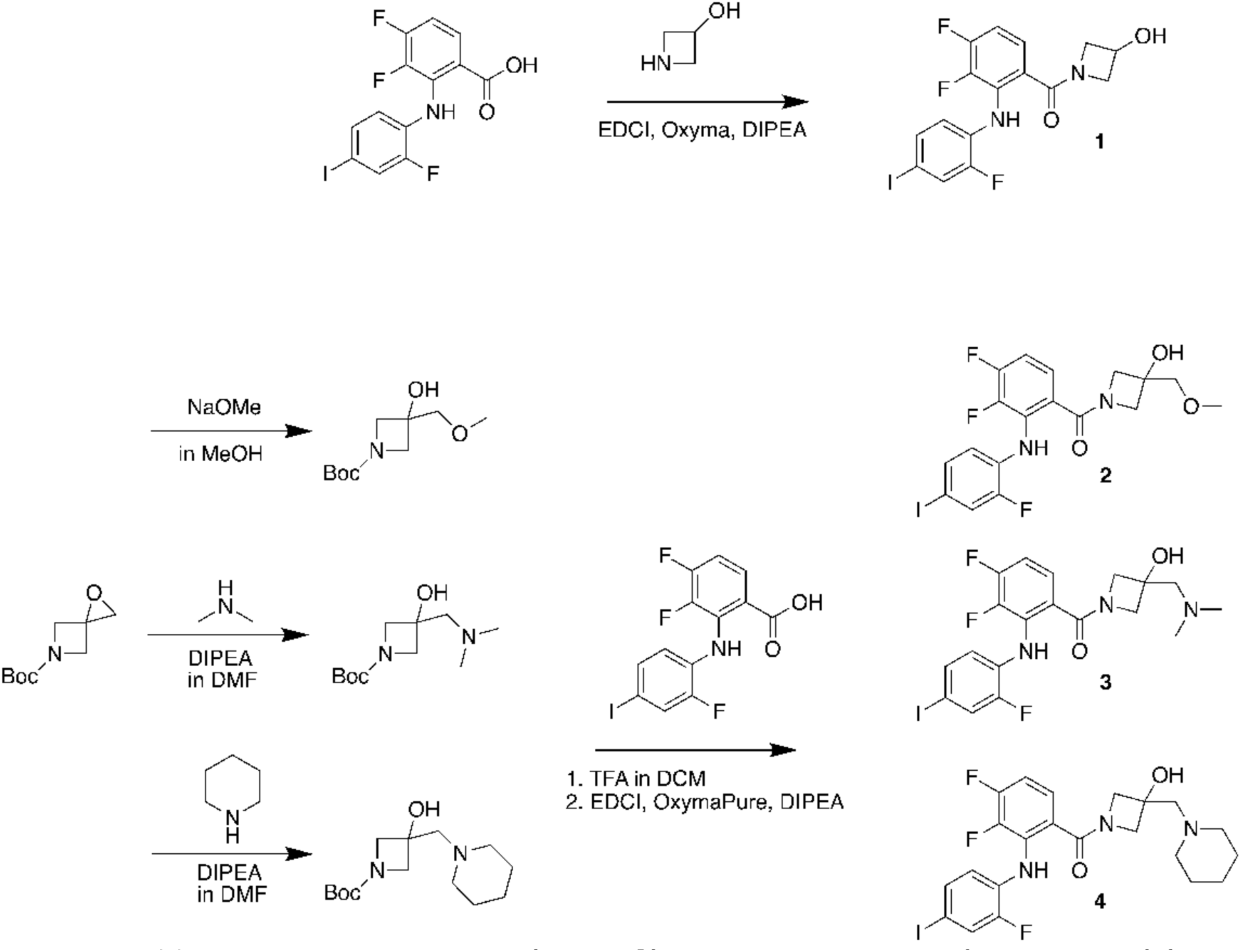

**Synthesis of fluoro-4-iodophenylamino)phenyl](3-hydroxyazetidin-1-yl)methanone (1)**

To a 0.5 mL solution of DIPEA (34 mg, 0.266 mmol) (Sigma Aldrich) in dry DMF, **3,4-difluoro-2-(2-fluoro-4-iodophenylamino)benzoic acid** was dissolved with stirring. **azetidin-3-ol hydrochloride** was then added and dissolved. OxymaPure (13 mg (0.091 mmol) (Sigma Aldrich) was added and dissolved. Lastly, EDCI-HCl (17 mg, 0.091 mmol) (Sigma Aldrich) was added to the reaction, which was stirred overnight at room temperature. 25 mL ethyl acetate was added to the reaction mixture and was washed twice with saturated ammonium chloride, twice with saturated sodium bicarbonate, and twice with brine. The resulting organic layer was dried over Na_2_SO_4_, decanted, and rotary evaporated. The crude residue was then purified by HPLC to afford **[3,4-difluoro-2-(2-fluoro-4-iodophenylamino)phenyl](3-hydroxyazetidin-1-yl)methanone (1)** (0.00982 mmol, 12.9% yield). ^1^H NMR (300 MHz, CDCl_3_) δ 8.49 (s, 1H), 7.51 – 7.30 (m, 2H), 7.14 (t, J = 7.4 Hz, 1H), 6.83 (q, J = 8.3 Hz, 1H), 6.62 (m, 1H), 4.86 – 4.64 (m, 1H), 4.52 – 4.37 (m, 2H), 4.10 (s, 2H). MS (ESI, m/z) calculated mass = 447.99, [M+H]^+^ found 448.9.

**Synthesis of [3,4-difluoro-2-(2-fluoro-4-iodophenylamino)phenyl][3-hydroxy-2-(methoxymethyl)azetidin-1-yl]methanone (2)**

To a 1 mL solution of sodium methoxide (44 mg, 0.81 mmol) in methanol, *tert*-butyl 1-oxa-5-azaspiro[2.3]hexane-5-carboxylate (Combi-Blocks) (50 mg, 0.27 mmol) was added. The reaction was stirred overnight at room temperature. The solution was then evaporated via rotary evaporation, the resulting residue was resuspended in 25 mL ethyl acetate and the organic layer was washed with 25 mL brine three times. The resulting ethyl acetate solution was then dried over Na_2_SO_4_, decanted, and evaporated to afford *tert*-butyl 3-hydroxy-3-(methoxymethyl)azetidine-1-carboxylate. The crude product was used in the next reaction without further purification.

Crude *tert*-butyl 3-hydroxy-3-(methoxymethyl)azetidine-1-carboxylate (20 mg, 0.091 mmol) was dissolved in 3 mL 30% TFA in DCM and stirred for 2 hours at room temperature. The solution was co-evaporated three times with toluene. The product was then dissolved with 3,4-difluoro-2-(2-fluoro-4-iodophenylamino)benzoic acid (35.8 mg, 0.091 mmol) in a 0.5 mL solution of DIPEA (34 mg, 0.266 mmol) (Sigma Aldrich) in dry DMF. OxymaPure (13 mg (0.091 mmol) (Sigma Aldrich) was added and dissolved. Lastly, EDCI-HCl (17 mg, 0.091 mmol) (Sigma Aldrich) was added to the reaction, which was stirred overnight at room temperature. 25 mL ethyl acetate was added to the reaction mixture and was washed twice with saturated ammonium chloride, twice with saturated sodium bicarbonate, and twice with brine. The resulting organic layer was dried over Na_2_SO_4_, decanted, and rotary evaporated. The crude residue was then purified by HPLC to afford [3,4-difluoro-2-(2-fluoro-4-iodophenylamino)phenyl][3-hydroxy-2-(methoxymethyl)azetidin-1-yl]methanone (2) (0.00772 mmol, 10.2% yield). ^1^H NMR (300 MHz, CDCl_3_) δ 8.31 (s, 1H), 7.50 – 7.29 (m, 2H), 7.14 (t, *J* = 7.4 Hz, 1H), 6.83 (q, *J* = 8.3 Hz, 1H), 6.62 (m, 1H), 4.13 (d, *J* = 9.3 Hz, 4H), 3.54 (s, 2H), 3.44 (d, *J* = 6.9 Hz, 3H). MS (ESI, m/z) calculated mass = 492.02, [M+H]^+^ found 493.0.

**Synthesis of [3,4-difluoro-2-(2-fluoro-4-iodophenylamino)phenyl]{2-[(dimethylamino)methyl]-3-hydroxyazetidin-1-yl}methanone (3)**

To a 1 mL solution of DIPEA (Sigma Aldrich) (104 mg, 0.81 mmol) in dry DMF, dimethylamine HCl (44 mg, 0.54 mmol) (Sigma Aldrich) was added and stirred until dissolved. *tert*-butyl 1-oxa-5-azaspiro[2.3]hexane-5-carboxylate (50 mg, 0.27 mmol) (Combi-Blocks) was then added and the reaction was stirred overnight at room temperature. 25 mL ethyl acetate was added and washed twice with saturated sodium bicarbonate, and twice with brine. The resulting organic layer was then dried over Na_2_SO_4_, decanted, and rotary evaporated to afford *tert*-butyl 3-[(dimethylamino)methyl]-3-hydroxyazetidine-1-carboxylate. The resulting residue was used in the next reaction without further purification.

Crude *tert*-butyl 3-[(dimethylamino)methyl]-3-hydroxyazetidine-1-carboxylate (21 mg, 0.091 mmol) was dissolved in 3 mL 30% TFA in DCM and stirred for 2 hours at room temperature. The solution was co-evaporated three times with toluene. The product was then dissolved with 3,4-difluoro-2-(2-fluoro-4-iodophenylamino)benzoic acid (35.8 mg, 0.091 mmol) in a 0.5 mL solution of DIPEA (34 mg, 0.266 mmol) (Sigma Aldrich) in dry DMF. OxymaPure (13 mg (0.091 mmol) (Sigma Aldrich) was added and dissolved. Lastly, EDCI-HCl (17 mg, 0.091 mmol) (Sigma Aldrich) was added to the reaction, which was stirred overnight at room temperature. 25 mL ethyl acetate was added to the reaction mixture and was washed twice with saturated ammonium chloride, twice with saturated sodium bicarbonate, and twice with brine. The resulting organic layer was dried over Na_2_SO_4_, decanted, and rotary evaporated. The crude residue was then purified by HPLC to afford [3,4-difluoro-2-(2-fluoro-4-iodophenylamino)phenyl]{2-[(dimethylamino)methyl]-3-hydroxyazetidin-1-yl}methanone (3) (0.0113 mmol, 14.9% yield). ^1^H NMR (300 MHz, CDCl_3_) δ 8.34 (s, 1H), 7.50 - 7.28 (m, 2H), 7.13 (t, *J* = 7.2 Hz, 1H), 6.85 (q, *J* = 8.3 Hz, 1H), 6.74 – 6.52 (m, 1H), 4.17 (s, 4H), 3.38 (s, 2H), 2.92 (s, 6H). MS (ESI, m/z) calculated mass = 505.05, [M+H]^+^ found 506.1.

**Synthesis of [3,4-difluoro-2-(2-fluoro-4-iodophenylamino)phenyl]{3-hydroxy-2-[(piperidin-1-yl)methyl]azetidin-1-yl}methanone (4)**

To a 1 mL solution of DIPEA (Sigma Aldrich) (104 mg, 0.81 mmol) in dry DMF, piperidine (46 mg, 0.54 mmol) (Sigma Aldrich) was added with stirring. *tert*-butyl 1-oxa-5-azaspiro[2.3]hexane-5-carboxylate (50 mg, 0.27 mmol) (Combi-Blocks) was then added and the reaction was stirred overnight at room temperature. 25 mL ethyl acetate was added and washed twice with saturated sodium bicarbonate, and twice with brine. The resulting organic layer was then dried over Na_2_SO_4_, decanted, and rotary evaporated to afford *tert*-butyl 3-hydroxy-3-[(piperidin-1-yl)methyl]azetidine-1-carboxylate. The resulting residue was used in the next reaction without further purification.

Crude ***tert*-butyl 3-hydroxy-3-[(piperidin-1-yl)methyl]azetidine-1-carboxylate** (25 mg, 0.091 mmol) was dissolved in 3 mL 30% TFA in DCM and stirred for 2 hours at room temperature. The solution was co-evaporated three times with toluene. The product was then dissolved with **3,4-difluoro-2-(2-fluoro-4-iodophenylamino)benzoic acid** (35.8 mg, 0.091 mmol) in a 0.5 mL solution of DIPEA (34 mg, 0.266 mmol) (Sigma Aldrich) in dry DMF. OxymaPure (13 mg (0.091 mmol) (Sigma Aldrich) was added and dissolved. Lastly, EDCI-HCl (17 mg, 0.091 mmol) (Sigma Aldrich) was added to the reaction, which was stirred overnight at room temperature. 25 mL ethyl acetate was added to the reaction mixture and was washed twice with saturated ammonium chloride, twice with saturated sodium bicarbonate, and twice with brine. The resulting organic layer was dried over Na_2_SO_4_, decanted, and rotary evaporated. The crude residue was then purified by HPLC to afford **[3,4-difluoro-2-(2-fluoro-4-iodophenylamino)phenyl]{3-hydroxy-2-[(piperidin-1-yl)methyl]azetidin-1-yl}methanone** (**4)** (0.0165 mmol, 21.7% yield). 1H NMR (300 MHz, CDCl3) δ 8.37 (s, 1H), 7.51 – 7.31 (m, 2H), 7.12 (t, *J* = 7.0 Hz, 1H), 6.94 – 6.71 (m, 1H), 6.63 (td, *J* = 8.6, 5.2 Hz, 1H), 4.38 (s, 1H), 4.13 (d, *J* = 9.6 Hz, 3H), 3.59 (d, *J* = 53.9 Hz, 4H), 3.27 (s, 2H), 2.72 (s, 2H), 2.02 (s, 1H), 1.56 – 1.22 (m, 2H).

**Synthesis of LF-268**

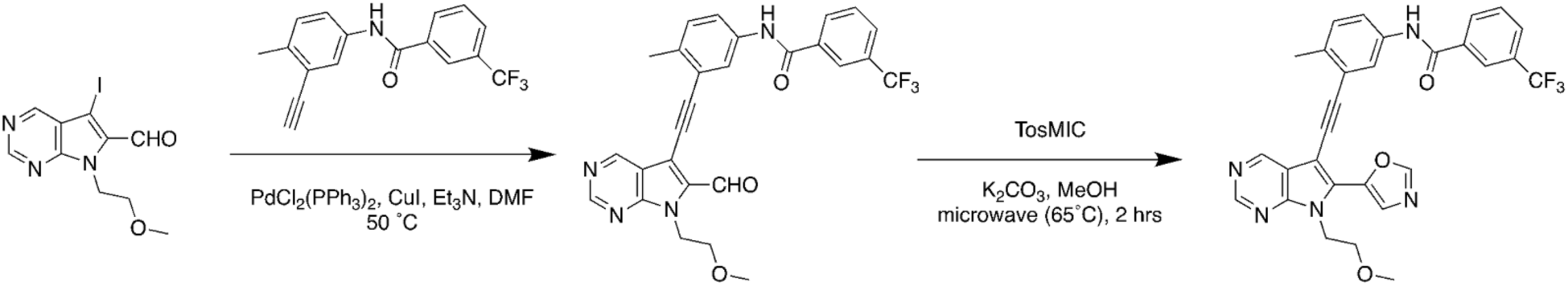

**Synthesis of *N*-(3-((6-formyl-7-(2-methoxyethyl)-7*H*-pyrrolo[2,3-*d*]pyrimidin-5-yl)ethynyl)-4-methylphenyl)-3-(trifluoromethyl)benzamide**

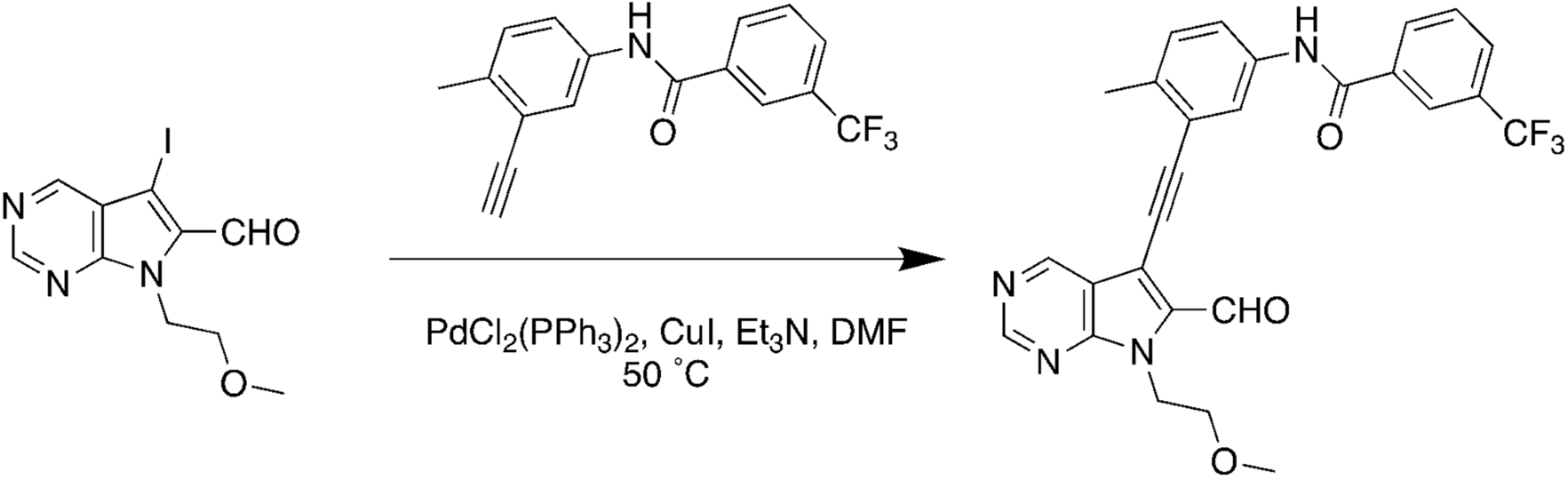

*N*-(3-ethynyl-4-methylphenyl)-3-(trifluoromethyl)-benzamide was synthesized according to a previous report (PMID: 30956043). 5-Iodo-7-(2-methoxyethyl)-7H-pyrrolo[2,3-d]pyrimidine-6-carbaldehyde (150 mg, 0.45 mmol, 1.0 equiv.) was dissolved in dry DMF (6 mL). The solution was flushed with a moderate stream of nitrogen gas for 10 minutes. Triethylamine (190 mg, 250 mL, 1.8 mmol, 4.0 equiv.), *N*-(3-ethynyl-4-methylphenyl)-3-(trifluoromethyl)benzamide (210 mg, 0.68 mmol, 1.5 equiv.), bis(triphenylphosphine)palladium(II) dichloride (16 mg, 0.023 mmol, 0.05 equiv.) and copper(I) iodide (8.6 mg, 0.045 mmol, 0.10 equiv.) were added sequentially. The resulting mixture was stirred at room temperature for 30 minutes and then maintained at 50°C for 14 hours. The solvent was removed in vacuo and the solid residue was dissolved in EtOAc (300 mL). The organic layer was washed with saturated aqueous NH4Cl, saturated aqueous NaHCO3, brine and then dried over anhydrous Na_2_SO_4_. Purification by flash chromatography on silica gel (EtOAc/Hexane = 0:100 to EtOAc/Hexane = 60:40) afforded *N*-(3-((6-formyl-7-(2-methoxyethyl)-7H-pyrrolo[2,3-d] pyrimidin-5-yl)ethynyl)4-methylphenyl)-3-(trifluoromethyl)benzamide as a pale yellow solid (189 mg, 80%). ^1^H-NMR (300 MHz, CDCl3) d = 10.37 (s, 1H), 9.29 (s, 1H), 9.10 (s, 1H), 8.16 (s, 1H), 8.11 (d, J = 7.8 Hz, 1H), 8.03 (s, 1H), 7.94 (s, 1H), 7.82 (d, J = 7.7 Hz, 1H), 7.72–7.54 (m, 2H), 7.29 (d, J = 8.3 Hz, 1H), 4.92 (t, J = 5.2 Hz, 2H), 3.77 (t, J = 5.2 Hz, 2H), 3.30 (s, 3H), 2.55 (s, 3H); ^13^C-NMR (126 MHz, CDCl3) d = 181.91, 164.40, 156.03, 152.49, 152.16, 137.08, 135.73, 135.63, 135.51, 131.14, 130.68, 130.47, 129.55, 128.64, 124.19, 123.97, 123.72, 122.55, 121.84, 118.45, 110.33, 96.83, 82.47, 71.26, 59.01, 43.15, 20.52; MS (ESI, m/z) calculated mass = 506.16, [M+H]^+^ found 508.0.

**Synthesis of *N*-(3-((7-(2-methoxyethyl)-6-(oxazol-5-yl)-7*H*-pyrrolo[2,3-*d*]pyrimidin-5-yl)ethynyl)-4-methylphenyl)-3-(trifluoromethyl)benzamide**

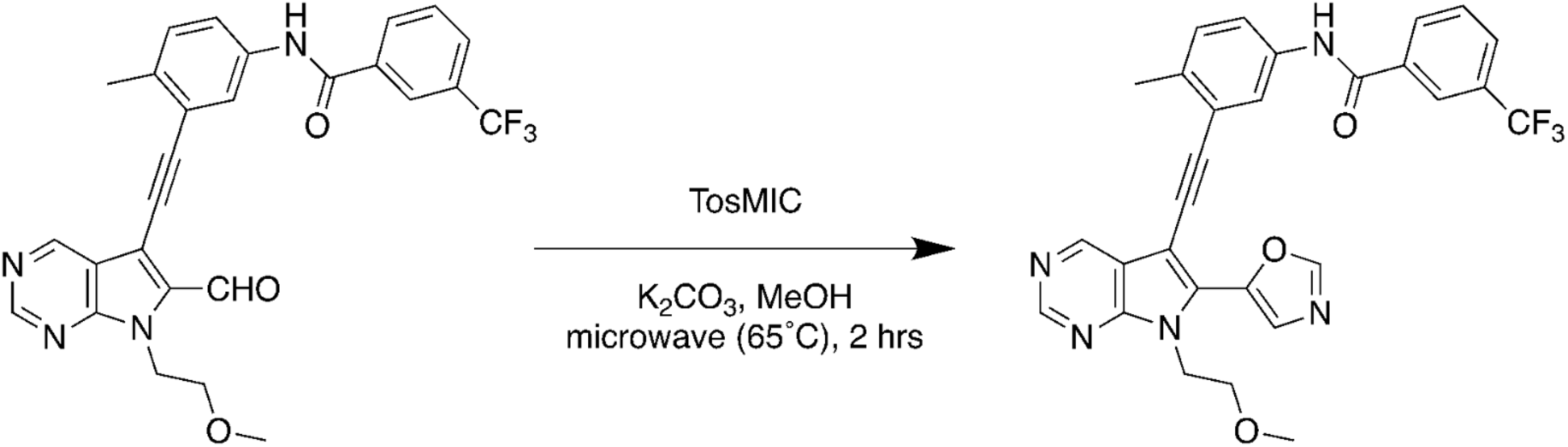

*N*-(3-((6-formyl-7-(2-methoxyethyl)-7*H*-pyrrolo[2,3-*d*]pyrimidin-5-yl)ethynyl)-4-methylphenyl)-3-(trifluoromethyl)benzamide (10 mg, 0.020 mmol, 1.0 equiv.) and toluenesulfonylmethyl isocyanide (5.8 mg, 0.030 mmol, 1.5 equiv.) were dissolved in methanol (700 µL) in a microwave reaction vial. Potassium carbonate (6.8 mg, 0.049 mmol, 2.5 equiv.) was added to the above solution sequentially. The reaction was heated at 65 °C in a microwave reactor for 2 hours. The resulting reaction mixture was diluted with ethyl acetate (10 mL) and the organic phase was washed with water (2 mL) and brine (2 mL), and then dried over anhydrous Na2SO4. Purification by flash chromatography on silica gel using a gradient of EtOAc/Hexane=0:100 to 100:0 afforded *N*-(3-((7-(2-methoxyethyl)-6-(oxazol-5-yl)-7*H*-pyrrolo[2,3-*d*]pyrimidin-5-yl)ethynyl)-4-methylphenyl)-3-(trifluoromethyl)benzamide as a pale brown solid (9.3 mg, 86%).^1^H NMR (300 MHz, CDCl_3_) δ 9.23 (s, 1H), 9.06 (s, 1H), 8.32 – 7.97 (m, 6H), 7.92 – 7.54 (m, 5H), 4.86 (t, *J* = 5.4 Hz, 2H), 3.78 (t, *J* = 5.3 Hz, 2H), 3.28 (s, 3H), 2.55 (s, 3H). MS (ESI, m/z) calculated mass = 545.1675, [M+H]^+^ found 546.1.

**Synthesis of LF-268-TCO**

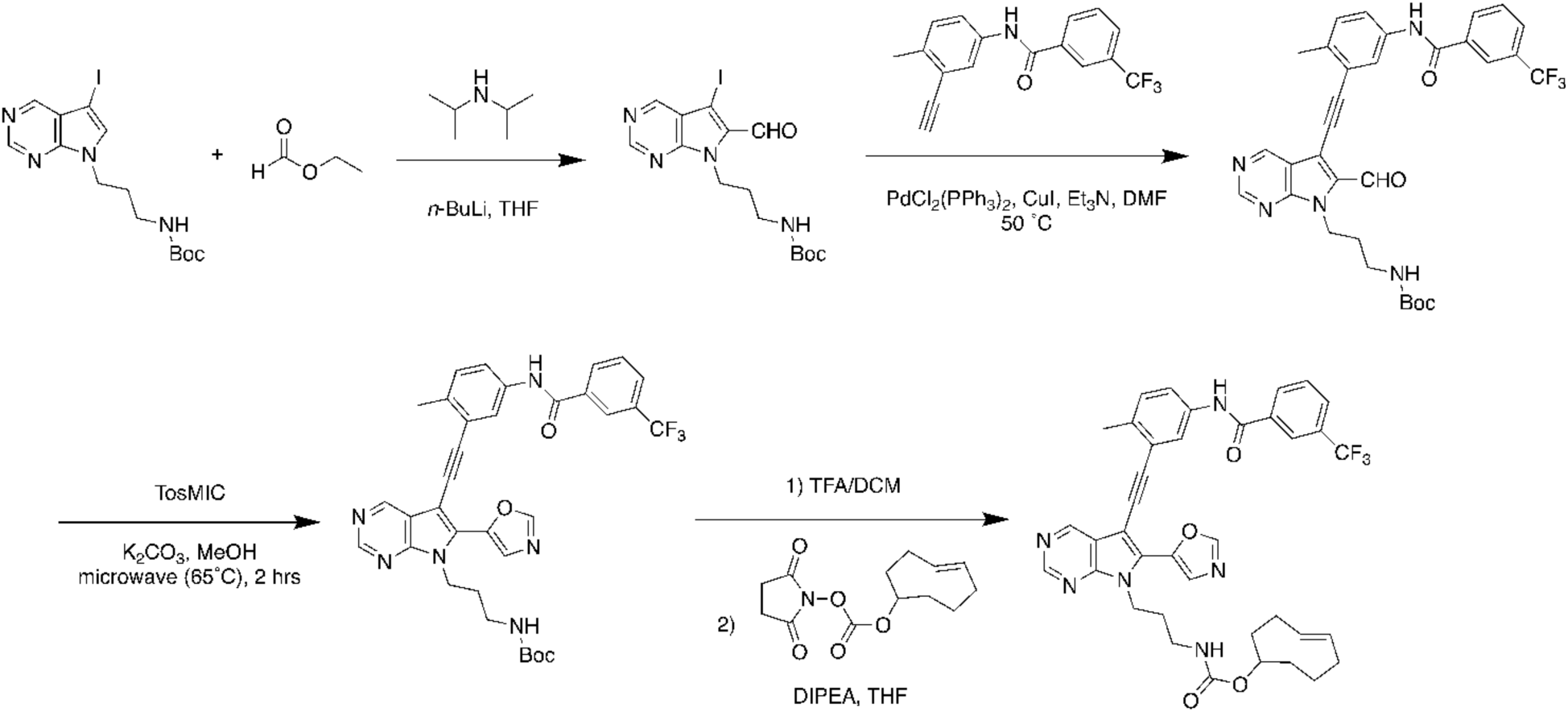

**Synthesis of *tert*-butyl (3-(6-formyl-5-iodo-7H-pyrrolo[2,3-d]pyrimidin-7-yl)propyl) carbamate**

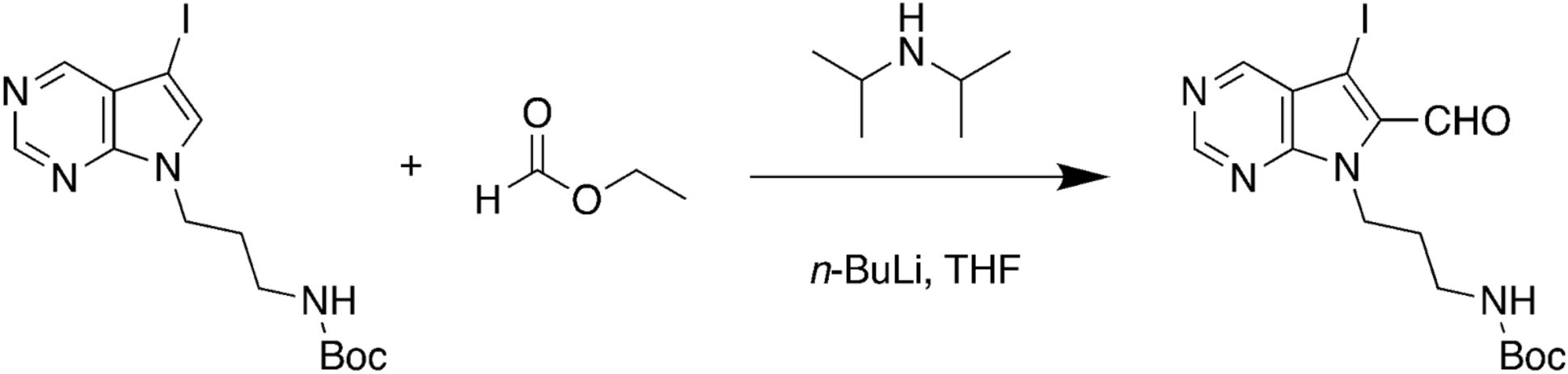

Diisopropylamine (490 mg, 680 μL, 4.9 mmol, 2.7 equiv.) was dissolved in dry THF (6 mL) and cooled to −78°C. A solution of n-BuLi (2.5 M in hexane, 1.8 mL, 4.5 mmol, 2.5 equiv.) was added dropwise to the above solution at −78 °C. The solution was stirred at 0 °C for 1 hour and then cooled to −78 °C. A solution of tert-butyl (3-(5-iodo-7H-pyrrolo[2,3-d]pyrimidin-7yl)propyl)carbamate (730 mg, 1.8 mmol, 1.0 equiv., dissolved in 3 mL dry THF) was added dropwise and the reaction mixture was then stirred at −78 °C for 1 hour. A solution of ethyl formate (536 mg, 580 µL, 7.2 mmol, 4.0 equiv., dissolved in 3 mL dry THF) was then added dropwise at −78°C. The reaction was allowed to warm to room temperature overnight. After reaction, saturated NH4Cl solution (2 mL) was added and the resulting mixture was diluted with ethyl acetate (200 mL). The organic layer was washed with saturated aqueous NaHCO3 solution, brine and dried over anhydrous Na2SO4. Purification by flash chromatography on silica gel (EtOAc: Hexane= 0:100 to EtOAc: Hexane= 3:2) afforded tert-butyl (3-(6-formyl-5-iodo-7H-pyrrolo[2,3d]pyrimidin-7-yl)propyl)carbamate as a pale brown solid (250 mg, 32%). ^1^H-NMR (300 MHz, CDCl3) δ= 10.02 (s, 1H), 9.07 (s, 1H), 8.98 (s, 1H), 5.22 (s, 1H), 4.73 (t, J = 6.6 Hz, 2H), 3.13 – 2.96 (m, 2H), 2.10 – 1.91 (m, 2H), 1.45 (s, 9H). MS (ESI, m/z) calculated mass = 430.1, [M+H]^+^ found 431.0.

**Synthesis of *tert*-butyl (3-(6-formyl-5-((2-methyl-5-(3-(trifluoromethyl)benzamido)phenyl)ethynyl)-7Hpyrrolo[2,3-d]pyrimidin-7-yl)propyl)carbamate**

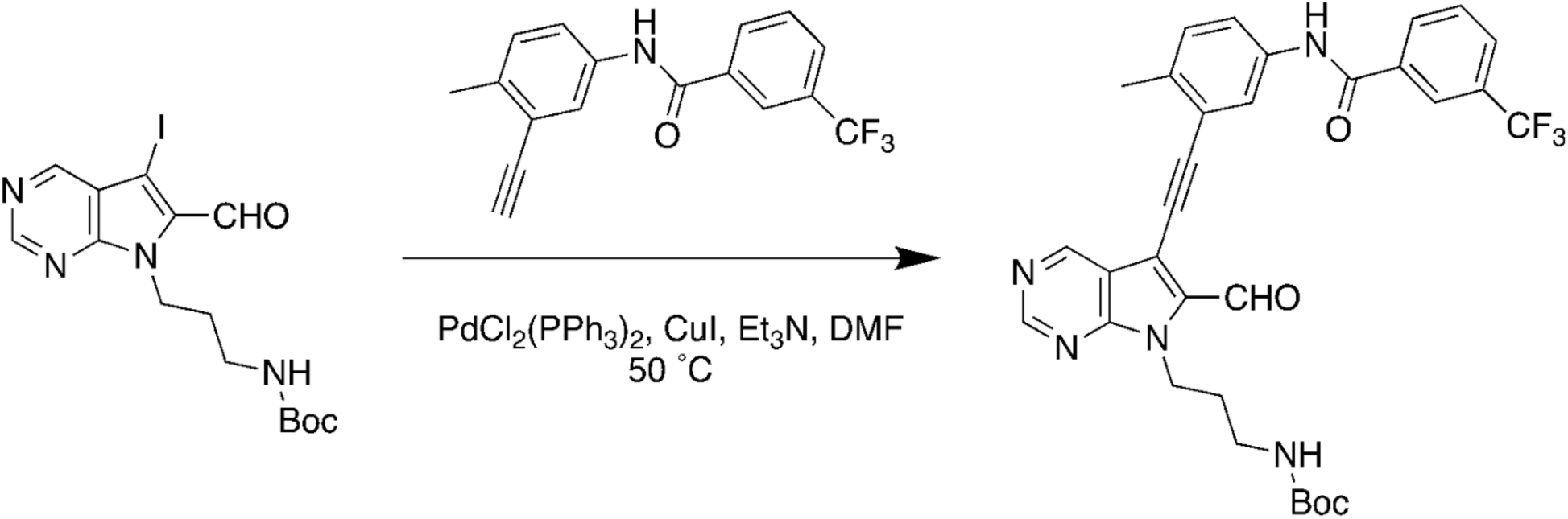

*N*-(3-ethynyl-4-methylphenyl)-3-(trifluoromethyl)-benzamide was synthesized according to previous reported (PMID: 30956043). *Tert*-butyl (3-(6-formyl-5-iodo-7H-pyrrolo[2,3-d]pyrimidin7-yl)propyl)carbamate (120 mg, 0.28 mmol, 1.0 equiv.) was dissovled in anhydrous DMF (3.5 mL) under nitrogen. Triethylamine (110 mg, 153 µL, 1.1 mmol, 4.0 equiv.), N-(3-ethynyl-4methylphenyl)-3-(trifluoromethyl)benzamide (130 mg, 0.42 mmol, 1.5 equiv.), bis(triphenylphosphine)palladium(II) dichloride (9.8 mg, 0.014 mmol, 0.05 equiv.) and copper (I) iodide (5.3 mg, 0.028 mmol, 0.1 equiv.) were added to the above solution sequentially. The reaction was heated at 50 °C for overnight and then quenched with saturated NH4Cl (1 mL). The resulting mixture was diluted with ethyl acetate (200 mL) and the organic phase was washed with saturated NaHCO3 (30 mL), brine (30 mL) and then dried over anhydrous Na2SO4. Purification by flash chromatography on silica gel using a gradient of EtOAc/Hexane (0:100 to 60:40) afforded tert-butyl (3-(6-formyl-5-((2-methyl-5-(3-(trifluoromethyl)benzamido)phenyl) ethynyl)-7Hpyrrolo[2,3-d]pyrimidin-7-yl)propyl)carbamate as a pale brown solid (65 mg, 39%; Rf= 0.42 in EtOAc:Hexane= 3:2). 1H-NMR (500 MHz, CDCl3) δ= 10.26 (s, 1H), 9.19 (s, 1H), 9.02 (s, 1H), 8.72 (s, 1H), 8.16 (s, 1H), 8.10 (d, J = 7.8 Hz, 1H), 7.92 (s, 1H), 7.76 (d, J = 7.7 Hz, 1H), 7.58 (d, J = 6.6 Hz, 1H), 7.21 (d, J = 8.3 Hz, 1H), 5.33 (s, 1H), 4.68 (t, J = 6.0 Hz, 2H), 3.12 – 2.95 (m, 2H), 2.48 (s, 3H), 2.02 – 1.91 (m, 2H), 1.43 (s, 9H). MS (ESI, m/z) calculated mass = 605.2, [M+H]^+^ found 606.2.

**Synthesis of *tert*-butyl (3-(5-((2-methyl-5-(3-(trifluoromethyl)benzamido)phenyl)ethynyl)-6-(oxazol-5-yl)-7*H*-pyrrolo[2,3-*d*]pyrimidin-7-yl)propyl)carbamate**

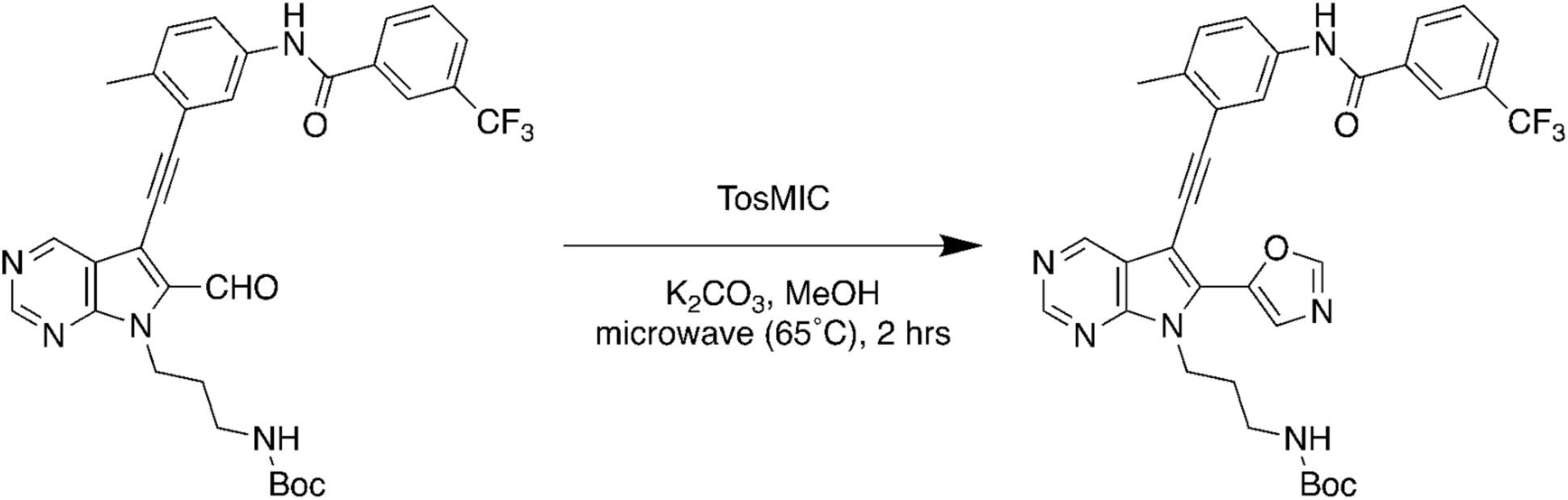

*Tert*-butyl (3-(6-formyl-5-((2-methyl-5-(3-(trifluoromethyl)benzamido)phenyl)ethynyl)-7H-pyrrolo[2,3-d]pyrimidin-7-yl)propyl)carbamate (11 mg, 0.018 mmol, 1.0 equiv.) and toluenesulfonylmethyl isocyanide (5.3 mg, 0.027 mmol, 1.5 equiv.) were dissolved in methanol (650 µL) in a microwave reaction vial. Potassium carbonate (6.3 mg, 0.045 mmol, 2.5 equiv.) was added to the above solution sequentially. The reaction was heated at 65 °C in a microwave reactor for 2 hours. The resulting reaction mixture was diluted with ethyl acetate (10 mL) and the organic phase was washed with water (2 mL) and brine (2 mL), and then dried over anhydrous Na_2_SO_4_. Purification by flash chromatography on silica gel using a gradient of EtOAc/Hexane=0:100 to 60:40 afforded *tert*-butyl (3-(5-((2-methyl-5-(3-(trifluoromethyl)benzamido)phenyl)ethynyl)-6-(oxazol-5-yl)-7H-pyrrolo[2,3-d]pyrimidin-7-yl)propyl)carbamate as a pale brown solid (9.6 mg, 83%). Rf=0.30 (EtOAc/Hexane=60:40). ^1^H NMR (500 MHz, CDCl3) δ 8.92 (s, 1H), 8.22 (s, 1H), 8.11 (s, 1H), 8.05 (d, *J* = 8.4 Hz, 2H), 7.97 (s, 1H), 7.77 – 7.65 (m, 2H), 7.62 (d, *J* = 8.3 Hz, 1H), 7.56 (t, *J* = 7.8 Hz, 1H), 7.18 (s, 1H), 5.27 (s, 1H), 4.56 (t, *J* = 6.7 Hz, 2H), 3.02 (d, *J* = 7.2 Hz, 2H), 2.45 (s, 3H), 2.04 – 1.84 (m, 2H), 1.38 (s, 9H). MS (ESI, m/z) calculated mass = 644.2359, [M+H]^+^ found 645.3.

**Synthesis of (*E*)-cyclooct-4-en-1-yl (3-(5-((2-methyl-5-(3-(trifluoromethyl)benzamido)phenyl)ethynyl)-6-(oxazol-5-yl)-7*H*-pyrrolo[2,3-*d*]pyrimidin-7-yl)propyl)carbamate (268-TCO)**

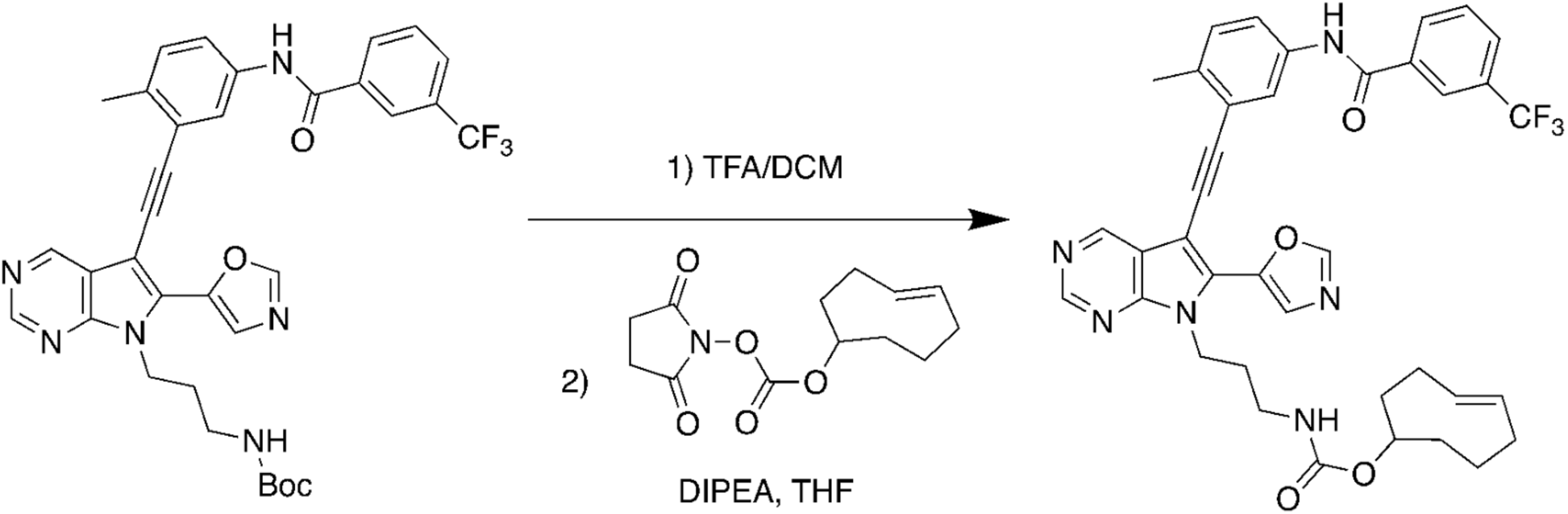

*Tert*-butyl (3-(5-((2-methyl-5-(3-(trifluoromethyl)benzamido)phenyl)ethynyl)-6-(oxazol-5-yl)-7*H*-pyrrolo[2,3-*d*]pyrimidin-7-yl)propyl)carbamate (13 mg, 0.020 mmol, 1.0 equiv.) was dissolved in a mixture of trifluoacetic acid (0.3 mL) and dichloromethane (0.7 mL). The reaction solution was stirred at room temperature for 30 mins and the solvent was removed in *vacuo*. The solid residue was dissolved in anhydrous tetrahydrofuran (1 mL). To the above solution was added diisopropylethylamine (13 mg, 18 μL, 0.099 mmol, 5.0 equiv.) and (*E*)-cyclooct-4-en-1-yl (2,5-dioxopyrrolidin-1-yl) carbonate (6.9 mg, 0.026 mmol, 1.3 equiv.) sequentially. The reaction was quenched with water (1 mL) after maintaining at room temperature overnight. The resulting mixture was then diluted with ethyl acetate (30 mL). The organic phase washed with saturated NaHCO_3_ solution (5 mL), brine (5 mL), dried over anhydrous Na_2_SO_4_. Purification using preparative TLC with a EtOAc/Hexane gradient (90:10) afforded (*E*)-cyclooct-4-en-1-yl (3-(5-((2-methyl-5-(3-(trifluoromethyl)benzamido)phenyl)ethynyl)-6-(oxazol-5-yl)-7*H*-pyrrolo[2,3-*d*]pyrimidin-7-yl)propyl)carbamate as a bright yellow powder (4.0 mg, 29% for 2 steps). R_f_= 0.5 in pure EtOAc/Hexane=90:10). ^1^H NMR (500 MHz, CDCl_3_) δ 9.05 (s, 1H), 8.92 (s, 1H), 8.58 (s, 1H), 8.19 (s, 1H), 8.12 (d, *J* = 7.9 Hz, 1H), 8.09 (s, 1H), 7.99 (s, 1H), 7.78 (d, *J* = 8.4 Hz, 2H), 7.67 (d, *J* = 8.3 Hz, 1H), 7.60 (t, *J* = 7.8 Hz, 1H), 7.22 (d, *J* = 8.3 Hz, 1H), 5.62 – 5.38 (m, 3H), 4.59 (t, *J* = 6.7 Hz, 2H), 3.34 (s, 0H), 3.15 – 3.02 (m, 2H), 2.94 (s, 1H), 2.85 (s, 1H), 2.49 (s, 3H), 2.37 – 2.24 (m, 3H), 2.08 – 1.84 (m, 5H), 1.81 – 1.49 (m, 3H).; MS (ESI, m/z) calculated mass = 696.2672, [M+H]^+^ found 697.1.

**carbamic acid, *N*-[[4-[3-[2,6-difluoro-3-[(propylsulfonyl)amino]benzoyl]-1*H*-pyrrolo[2,3-*b*]pyridin-5-yl]phenyl]methyl]-, 1,1-dimethylethyl ester**

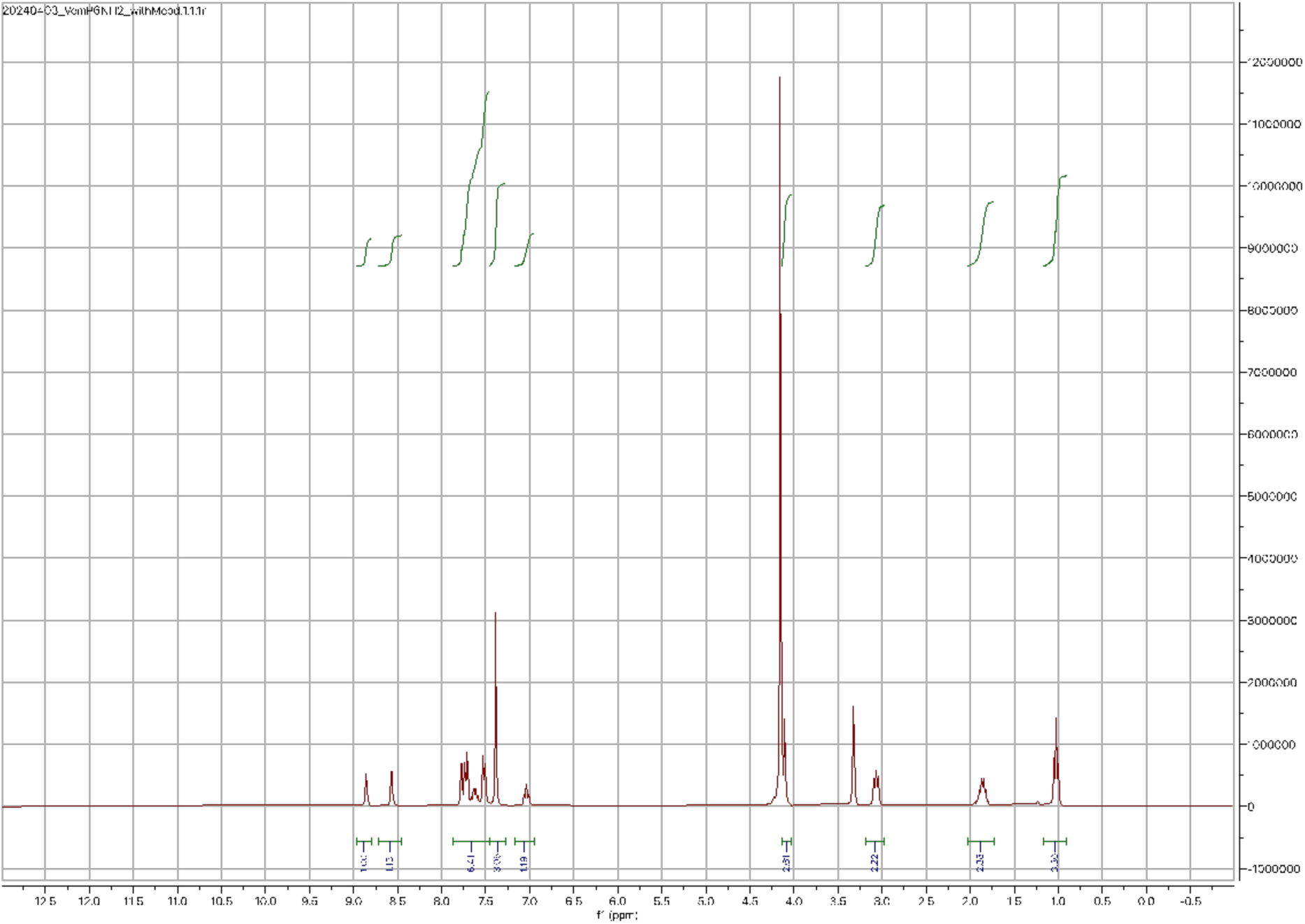

**20-{N-[(4-{3-[2,6-difluoro-3-(propylsulfonylamino)benzoyl]-1,7-diaza-1H-inden-5-yl}phenyl)methyl]carbamoyl}-3,6,9,12,15,18-hexaoxaicosyl 2-methylpropane-2-carbamate (6)**

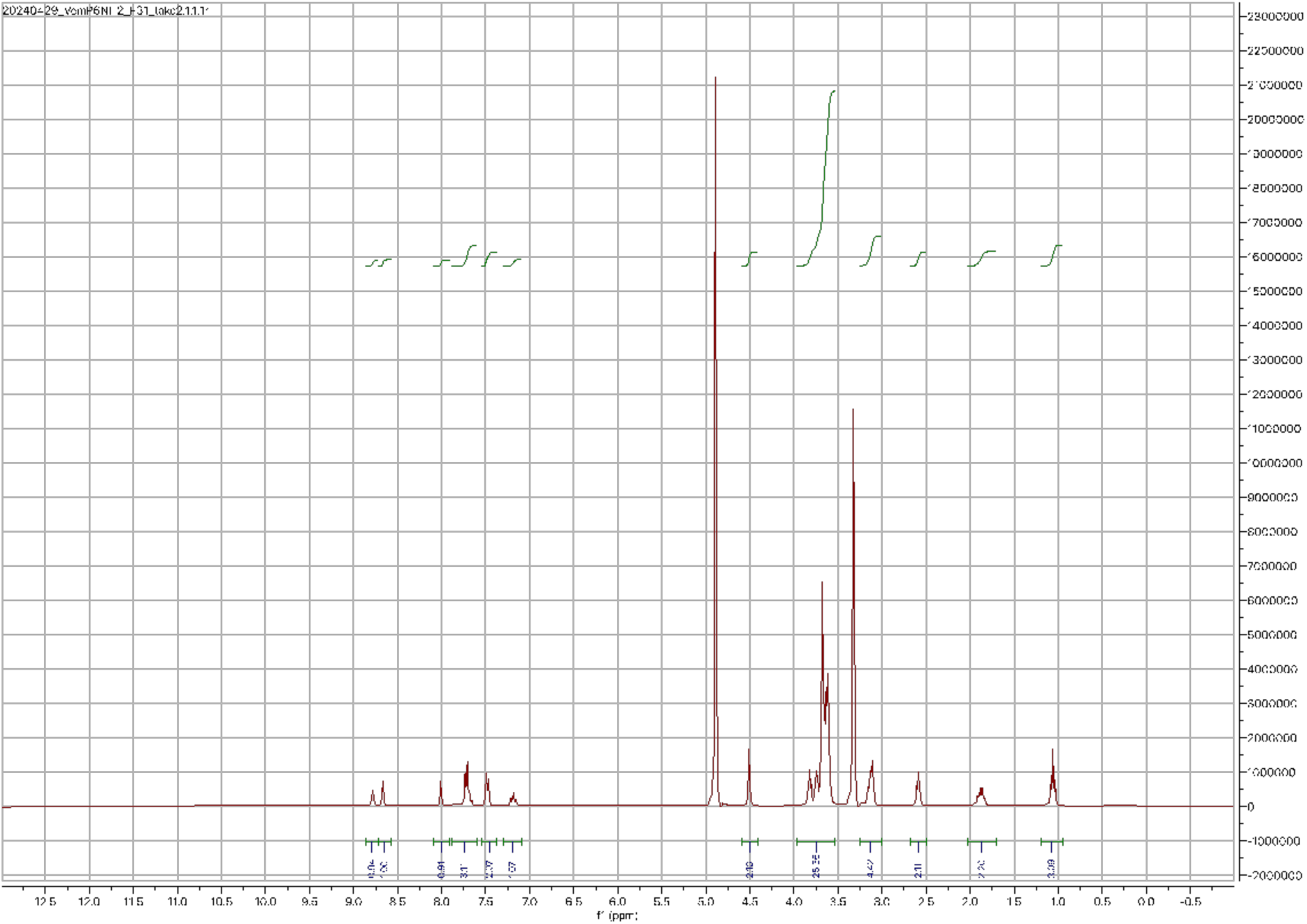

**[3,4-difluoro-2-(2-fluoro-4-iodophenylamino)phenyl](3-hydroxyazetidin-1-yl)methanone (1)**

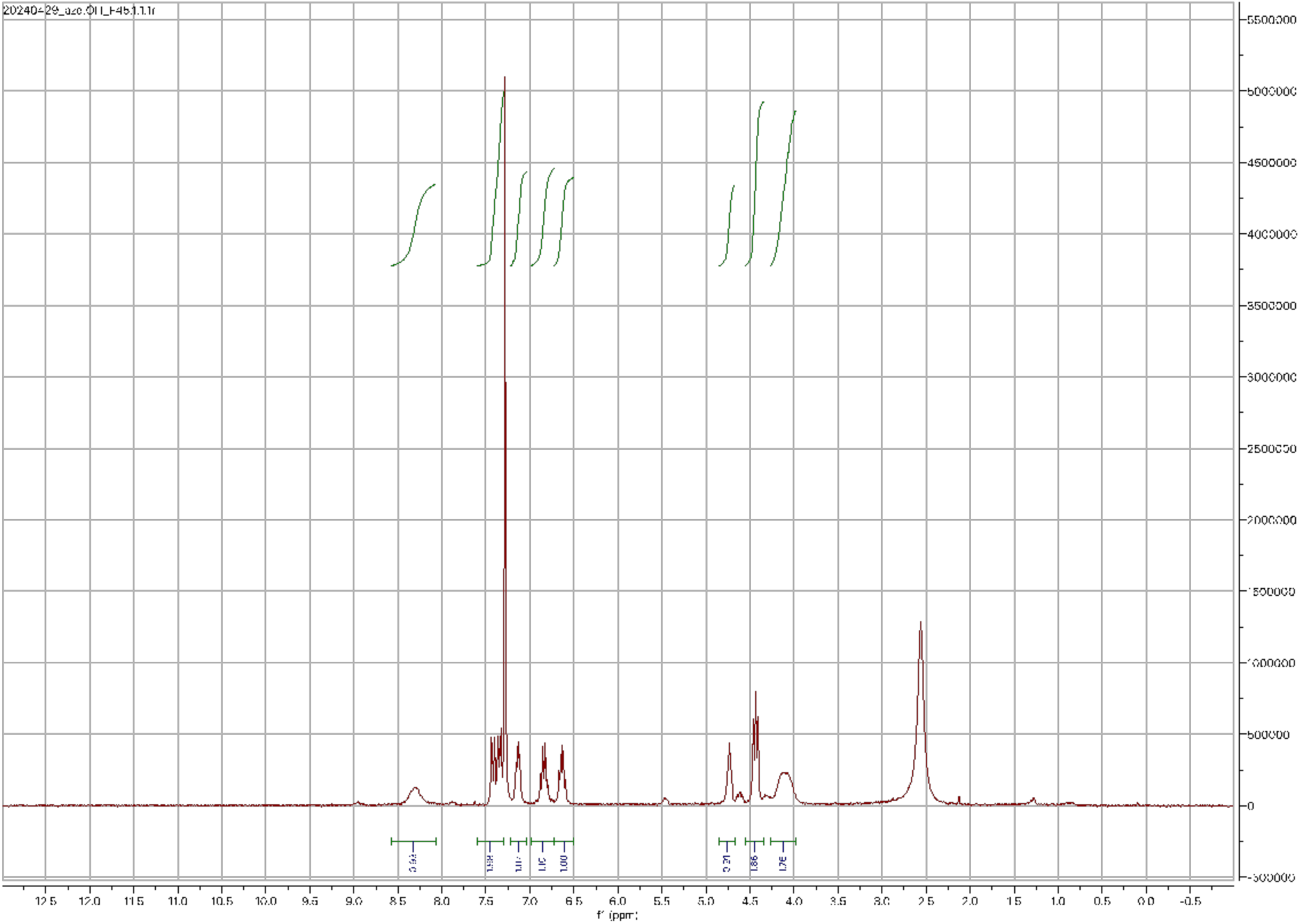

**[3,4-difluoro-2-(2-fluoro-4-iodophenylamino)phenyl][3-hydroxy-2-(methoxymethyl)azetidin-1-yl]methanone (2)**

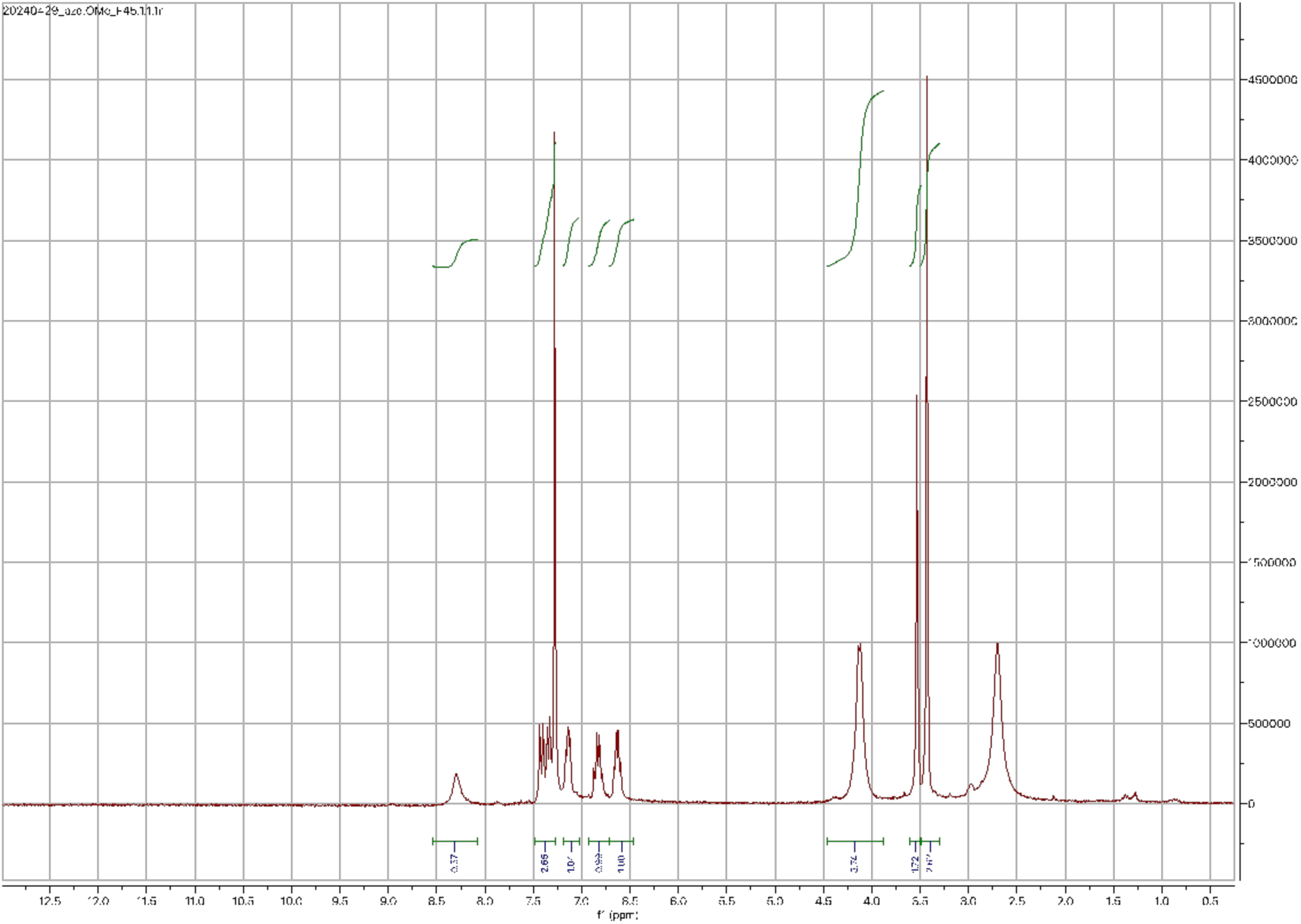

**[3,4-difluoro-2-(2-fluoro-4-iodophenylamino)phenyl]{2-[(dimethylamino)methyl]-3-hydroxyazetidin-1-yl}methanone** (**3)**

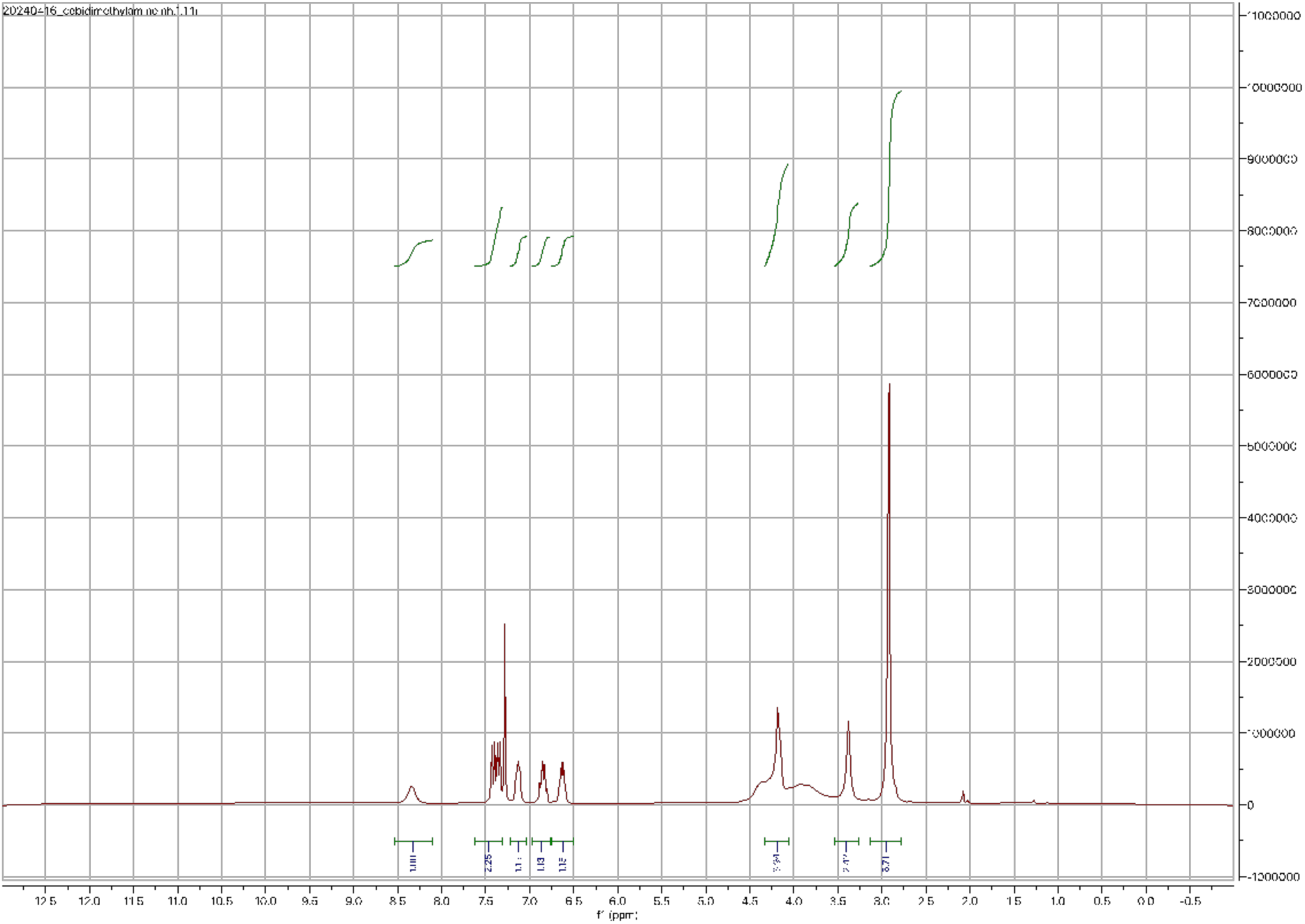

**[3,4-difluoro-2-(2-fluoro-4-iodophenylamino)phenyl]{3-hydroxy-2-[(piperidin-1-yl)methyl]azetidin-1-yl}methanone** (**4)**

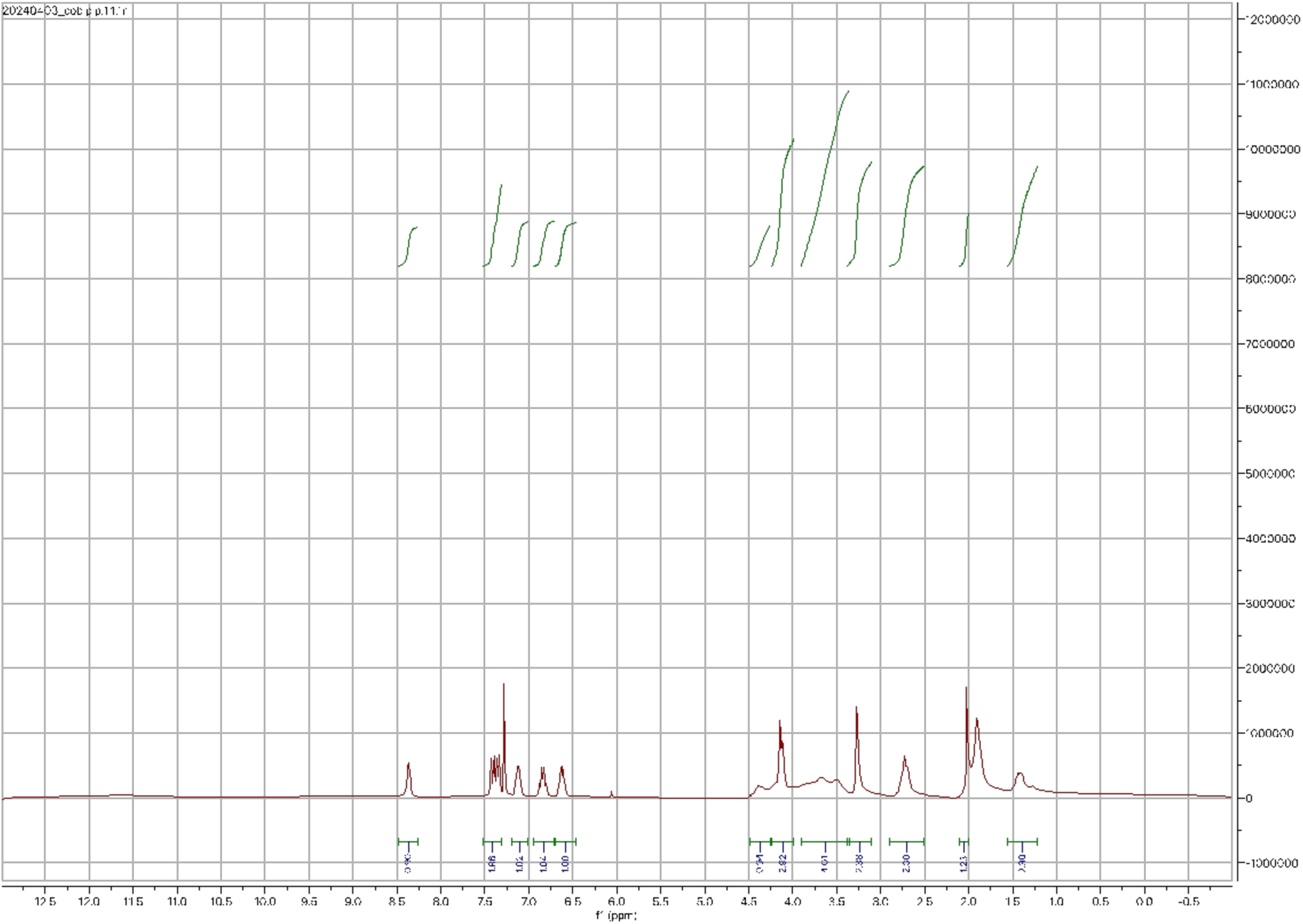

***N*-(3-((6-formyl-7-(2-methoxyethyl)-7*H*-pyrrolo[2,3-*d*]pyrimidin-5-yl)ethynyl)-4-methylphenyl)-3-(trifluoromethyl)benzamide**

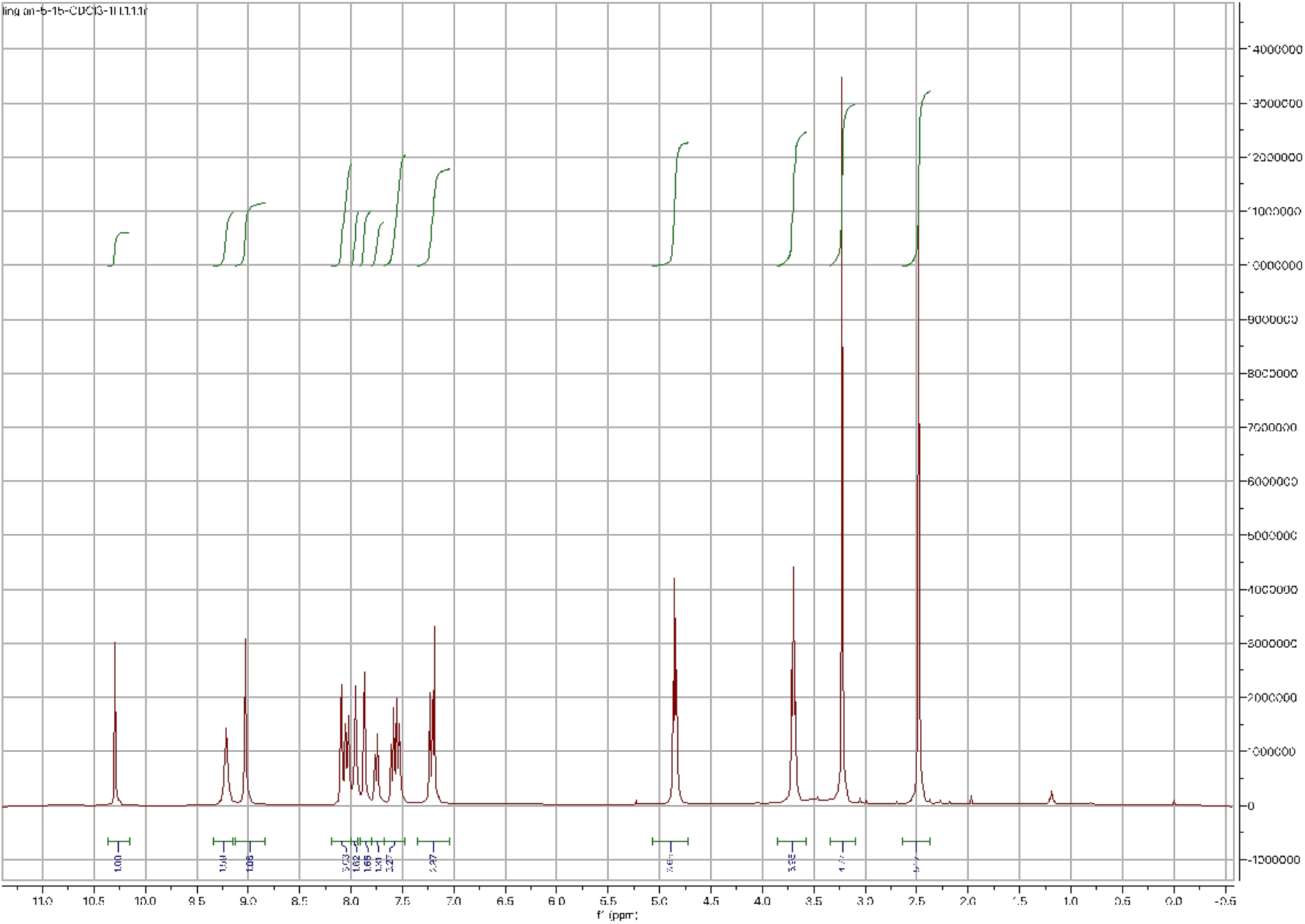

***N*-(3-((7-(2-methoxyethyl)-6-(oxazol-5-yl)-7*H*-pyrrolo[2,3-*d*]pyrimidin-5-yl)ethynyl)-4-methylphenyl)-3-(trifluoromethyl)benzamide**

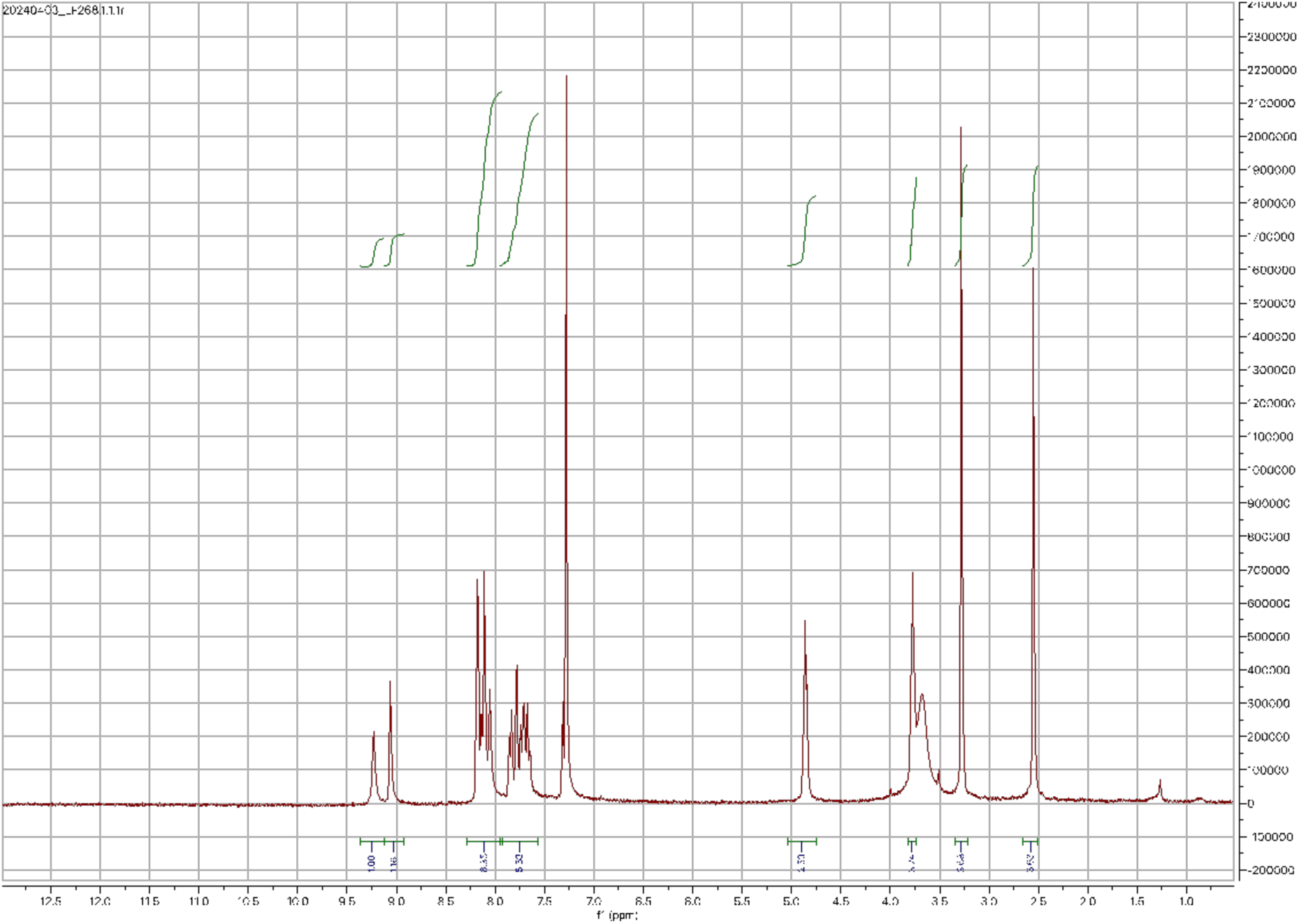

***tert*-butyl (3-(6-formyl-5-((2-methyl-5-(3-(trifluoromethyl)benzamido)phenyl)ethynyl)-7Hpyrrolo[2,3-d]pyrimidin-7-yl)propyl)carbamate**

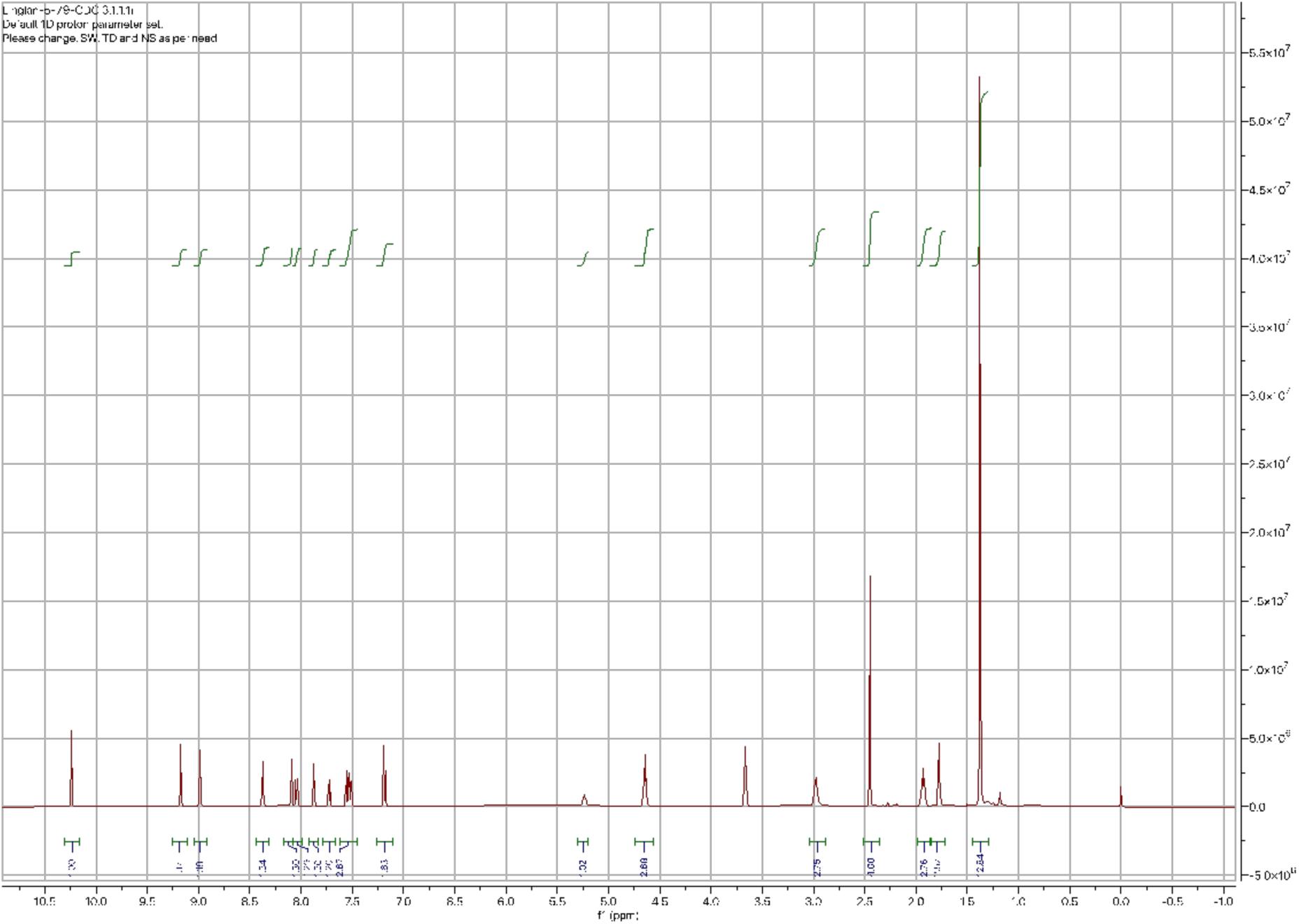

***tert*-butyl (3-(5-((2-methyl-5-(3-(trifluoromethyl)benzamido)phenyl)ethynyl)-6-(oxazol-5-yl)-7*H*-pyrrolo[2,3-*d*]pyrimidin-7-yl)propyl)carbamate**

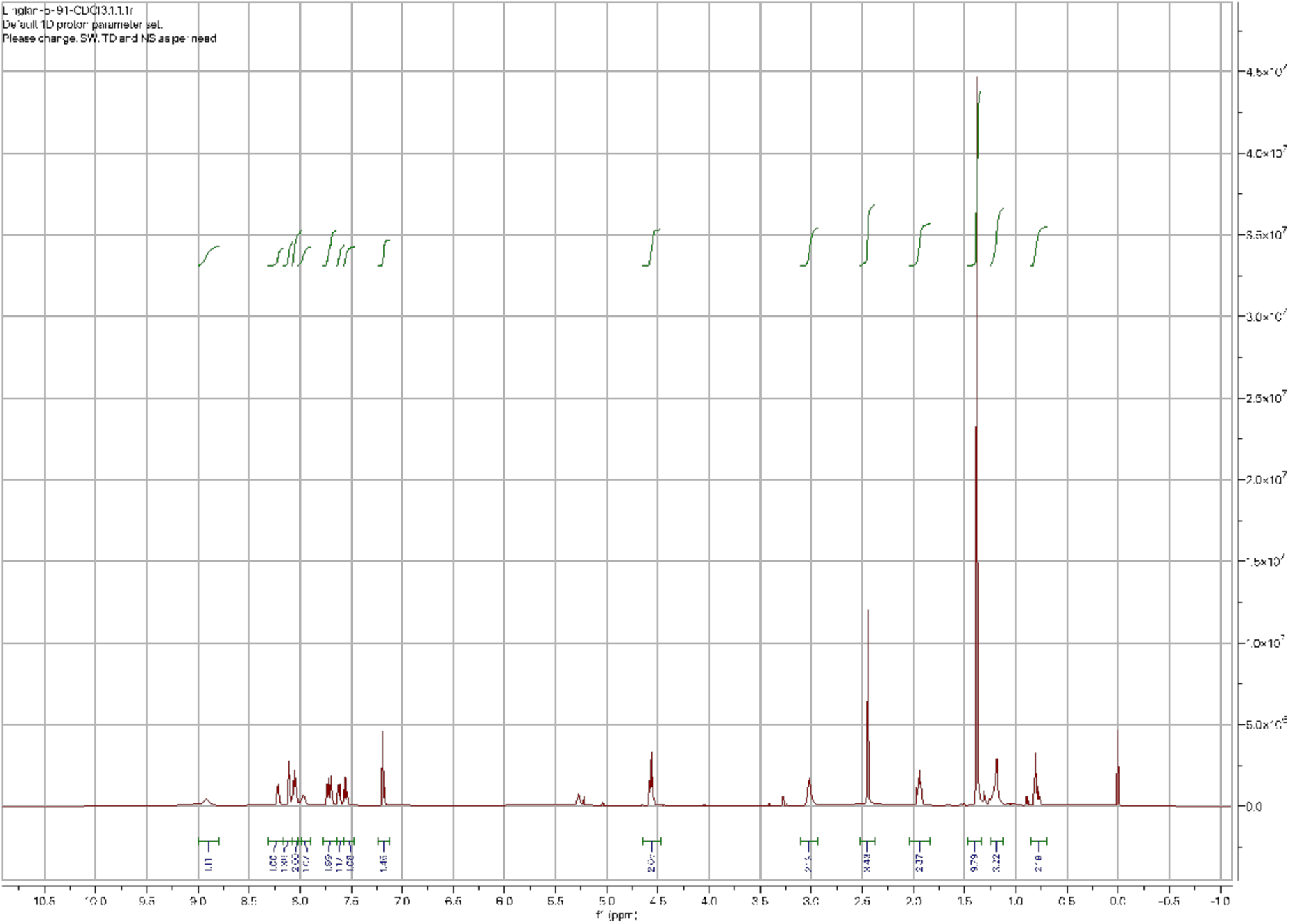

***tert*-butyl (3-(5-((2-methyl-5-(3-(trifluoromethyl)benzamido)phenyl)ethynyl)-6-(oxazol-5-yl)-7H-pyrrolo[2,3-d]pyrimidin-7-yl)propyl)carbamate**

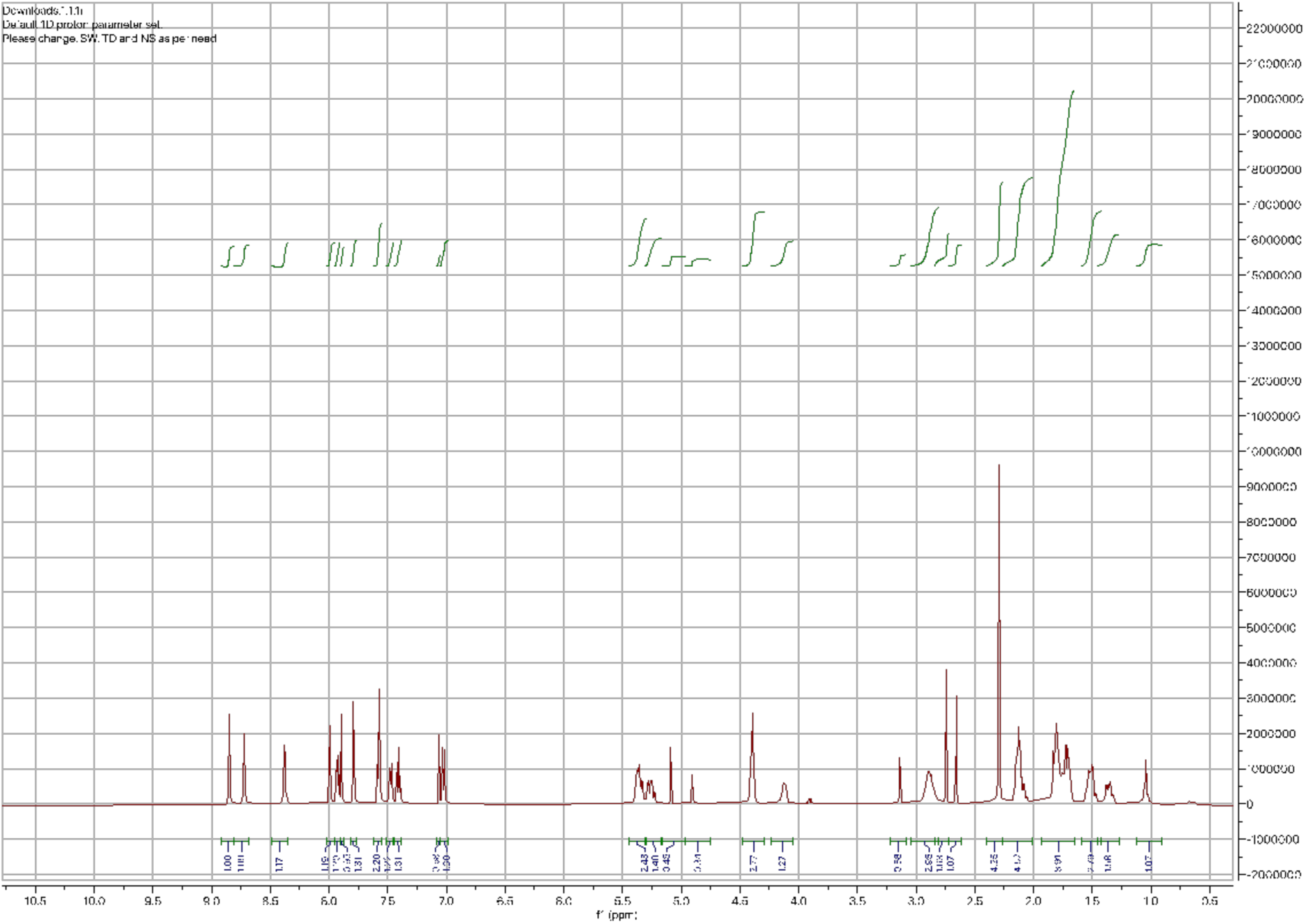

## Acknowledgements

This work was supported by NIH grant no. R01GM145011 (NIGMS) and R01GM086858 (NIGMS) awarded to D.J.M.

## Ethics Declarations

The authors declare no competing interests.

